# Reflectance spectra capture temporal variation in functional traits and their integration across leaf phenology

**DOI:** 10.64898/2026.04.21.719921

**Authors:** Cornelius Onyedikachi Nichodemus, Jose Eduardo Meireles

## Abstract

- Plant function depends not on single traits but on how traits covary, yet trait integration is typically estimated from a peak-season snapshot and treated as fixed. Currently, we do not understand if integration has a phenology and how it affects spectral trait models that depend on trait covariance.
- We measured traits and reflectance spectra weekly across a growing season in seven temperate species (7,515 spectra), compared trait and spectral correlation structure across three phenophases, and tested PLSR models trained on the full season or peak season alone, alongside a widely used published model.
- Traits followed species-specific trajectories, with spectral lability concentrated in the visible region. Correlation structure shifted with phenology: LMA–EWT, C–N, and N–QY held across phenophases, whereas LMA–N and VIS–NIR spectral correlations reversed sign. Full-season models tracked trait dynamics, whereas peak-season and published models failed outside their window, producing biologically impossible estimates.
- Traits and their integration have themselves a phenology, and a peak-season correlation matrix does not fully capture trait integration. Spectra recover trait dynamics only with phenologically representative training data, since models calibrated in one phenophase inherit its correlation structure — a shift that coincides with, and may underlie, their failure outside it.

## INTRODUCTION

Plant functional traits — measurable characteristics that underlie how plants interact with and respond to their environment (Violle *et al*., 2007) — provide an integrative framework for understanding fitness (McGill *et al*., 2006), ecosystem properties, and variation in function and distribution across spatio-temporal scales (Wright *et al*., 2005; Šímová *et al*., 2018). However, plant performance is not a function of any single trait. Traits covary, and the structure of that covariation — phenotypic integration — constrains which trait combinations are attainable and therefore how plants respond to their environment (Pigliucci, 2003; Díaz *et al*., 2016). The leaf economic spectrum is the canonical description of such structure in leaves, since it embodies trade-offs between resource acquisition and conservation and serves as a proxy for ecological strategy (Wright *et al*., 2004; Kraft *et al*., 2015). Leaf mass per area (LMA), for example, tracks investment in structure, defense, and longevity, whereas leaf nitrogen (N), much of which is allocated to Rubisco and chlorophyll, tracks photosynthetic capacity, and the negative relationship between the two is the axis along which most variation in leaf strategy is arrayed (Wright *et al*., 2004; de Bello *et al*., 2010; Reich, 2014).

Importantly, we know that leaf traits are not fixed over time: leaves develop, mature, and senesce over a growing season, and the traits that define leaf economics change with them. LMA increases as leaves expand, thicken, and accumulate structural material, and declines during senescence as complex compounds are broken down, and dry mass is lost (Morais *et al*., 2022). Leaf N peaks early in the growing season, when leaves invest heavily in Rubisco for carbon fixation, and declines late in the season as leaves senesce, and plants reabsorb nitrogen before abscission (Muller *et al*., 2011). Direct measures of physiology follow suit: the quantum yield of photosystem II (QY) rises toward peak growing season and falls as chlorophyll degrades (Skillman, 2008; Jiang *et al*., 2024; Yu *et al*., 2025). While these trajectories differ in timing, rate, and direction, trait relationships that describe plant strategy are commonly estimated as if they did not.

Because traits change at different rates and in different directions, their correlations should change as well. That is, trait integration should itself have a phenology. The same expectation follows from the environment, given that the correlation structure of functional traits varies across environmental gradients (Laughlin *et al*., 2017) and the growing season is itself a gradient in temperature, light, and water availability. Trait correlation structure is known to shift with ontogeny (Damián *et al*., 2018), between years (Pahadi *et al*., 2024), and between sampling dates within a season (Mason *et al*., 2020). Trait networks have been compared between coarse seasons (Rao *et al*., 2023), and summer and winter leaves of an evergreen differ in their trait relationships (Puglielli & Varone, 2018). Seasonal variation in trait means is well documented (McKown *et al*., 2013; Fajardo & Siefert, 2016; Yang *et al*., 2016; Chlus & Townsend, 2022; Niittynen *et al*., 2026) — in fact, Niittynen *et al*., (2026) report that species rankings are largely preserved through the season — but such studies do not ask whether the relationships among traits are equally stable. How integration changes across the phenophases of a single leaf cohort, from emergence to senescence, remains largely uncharacterized. This matters beyond pure description, since trait-based inference built on one phenophase inherits the correlation structure of that phenophase, and may therefore carry a systematic bias that replication within the phenophase cannot reveal.

Most functional trait research nevertheless relies on snapshot measurements taken during peak growing season, a consequence of logistical constraints, since examining traits across a full season requires recurrent wet-lab based trait measurements that are both time consuming and costly. Leaf reflectance spectroscopy addresses those constraints. Leaves reflect light differently across the solar spectrum (400–2500 nm) depending on their structure and chemistry, enabling phenotypic properties to be inferred from the spectral signature (Curran, 1989; Ustin *et al*., 2009; Ustin & Jacquemoud, 2020; Burnett *et al*., 2021b; Cavender-Bares *et al*., 2025, 2026). The visible region (VIS: 400–700 nm) is dominated by pigments — chlorophylls, carotenoids, and anthocyanins — whose pools turn over rapidly during leaf development and senescence, while the near-infrared (NIR: 700–1100 nm) and shortwave-infrared (SWIR: 1100–2500 nm) regions capture structural features, water, and secondary compounds such as phenolics, lignin, and cellulose, which turn over slowly (Curran, 1989; Kokaly *et al*., 2009; Ustin *et al*., 2009; Asner *et al*., 2014; Ustin & Jacquemoud, 2020). If spectral change tracks the underlying trait dynamics, the VIS should be more labile across phenology while the NIR and SWIR should be comparatively conserved. Remote sensing has long tracked canopy phenology (Caparros-Santiago *et al*., 2021; Gong *et al*., 2024), yet few studies have examined how phenology affects leaf-level full-range spectra (Chavana-Bryant *et al*., 2017, 2019; Chlus & Townsend, 2022), which spectral regions are conserved versus labile across phenophases, and how spectral change maps onto trait change.

Spectra are themselves strongly correlated signals. Neighboring and functionally related bands covary, and that covariance is leveraged by empirical models to estimate traits. Most trait models are derived from these data using partial least squares regression (PLSR; Wold *et al*., 2001), which relates observed traits to the full spectrum through a small number of latent components (Serbin *et al*., 2014; Yang *et al*., 2016; Couture *et al*., 2016; Ely *et al*., 2019; Kothari *et al*., 2023). These empirical models are typically trained on spectra and traits collected during peak growing season (but see Yang *et al*., 2016 at the leaf level, and Chlus & Townsend, 2022; Ji *et al*., 2026 for imaging spectroscopy). These facts, the spectral autocovariance and restricted temporal sampling, highlight two important knowledge gaps: (1) whether the covariance structure of the full leaf spectrum is stable across phenophases, and (2) whether trait models trained on peak-season data transfer across phenophases, given that they are not exposed to the full range of phenologically driven variation in spectra and traits.

Similarly, empirical spectral models also exploit a second covariance structure: the covariance among traits. A trait with no distinct optical signature of its own can still be predicted accurately when it covaries with a trait that has one — the constellation effect, named by Nunes *et al*., (2017) following Chadwick & Asner, (2016). We can think of traits lying on a continuum from core, with a direct influence on the reflectance spectrum, to peripheral, whose association with the spectrum is mediated by their correlations with core traits (Kothari & Schweiger, 2022). The consequence is stated plainly by Kothari *et al*., (2023): if trait covariance patterns vary among datasets, overfitting to the training data could cause such models to produce inaccurate trait estimates from new spectral data. That risk has so far been framed spatially, across sites and species assemblages. If trait integration itself varies across phenophases, then the spectral–trait map is non-stationary since a model calibrated in one phenophase encodes a correlation structure that no longer holds in another. As a consequence, we expect models calibrated at peak season to produce biased predictions.

In this study, we measured leaf-level reflectance spectra and functional traits weekly across a full growing season for seven temperate tree and shrub species, generating 7,515 spectra from leaf emergence to senescence. We used these data to characterize the phenology of traits, spectra, and their correlation structure, and to build and test trait prediction models. We developed three categories of PLSR models — *all-season* (trained on data from all leaf phenological stages), *week-as-covariate* (as *all-season*, with collection week added as a predictor), and *peak-season* (trained on peak-growing-season data only) — and evaluated them alongside published coefficients from a widely used model (Kothari *et al*., 2023). Specifically, we ask:

1. How do traits and spectra change across taxa and leaf phenological stages, and which spectral regions are conserved or labile?
2. How does the correlation structure — among traits, and among spectral bands — vary across leaf phenological stages?
3. Can spectra predict the temporal dynamics of traits, and how do models trained on a narrow time slice perform across the full season?
4. Do trait estimates differ among phenophases, and are the departures quantitative or qualitative?

## MATERIALS AND METHODS

### Taxon sampling and experimental setup

We sampled seven (7) temperate species (*Acer platanoides*, *A. rubrum*, *Betula papyrifera*, *Prunus nigra*, *Quercus rubra*, *Rhododendron catawbiense*, and *R. maximum* — Table S1) within the University of Maine, Orono campus (44°52’58’’N, 68°40’21’’W), representing trees and shrubs. We monitored changes in leaf traits and spectra weekly throughout the growing season, from shortly after leaf emergence, on May 31st, to old/senescent on November 6th, 2023, for a total of 24 weeks. To account for phenotypic variation, we sampled 27 leaves per species per week (3 leaves per branch, 3 branches per individual tree, 3 individuals). We stored one leaf/branch/week at -80 °C for chemical analysis. Leaves were collected in plastic bags with moist paper towels to reduce moisture loss before being transferred to the lab for further measurements.

### Leaf spectra and trait measurement

We measured 7,515 leaf-level reflectance spectra from leaves using a full-range (350 - 2500 nm) field spectroradiometer—an SVC HR-1024i (Spectra Vista Corp., Poughkeepsie, NY, USA)—with a leaf clip and artificial light sources. Leaf spectral measurements were taken immediately after field collection in the laboratory from one leaf per branch. We took 5 random scans on each adaxial surface to capture variation in leaf orientation and relative position to the light source, and averaged the leaf measurements. We adopted the measurement protocol (Burnett *et al*., 2021a), processed, and trimmed the spectral data to 400 - 2400 nm using the SPECTROLAB R package (Meireles *et al*., 2017). To assess the degree of temporal variability in spectra, we first computed the average spectrum for each species per week. Second, we used the 24 resulting spectra —which capture how the average spectral signature changes over time— to compute the coefficient of variation (CV; standard deviation divided by the mean) per wavelength. Spectral regions with higher CV are more labile over time than regions with lower CV. We also assessed variation in spectral space occupied at different phenological stages using the representative spectra mean per week.

Immediately after field collection, we measured each leaf’s fresh mass (g) using a Quintix 124-1S Sartorius weighing balance and scanned the leaf area with a Canon 300 table scanner. We processed the scanned leaf images using a custom R script and the magick R package (Ooms, 2026) and computed leaf area (cm^2^ and m^2^). We dried the leaves in a drying oven at 50 °C for 14 days, and measured their oven-dry mass (g). We then computed leaf dry mass per area (LMA - kg/m^2^) using equation 5 (Poorter *et al*., 2009) and equivalent water thickness (EWT - g/cm^2^) using equation 2 (Féret *et al*., 2019). We used Fluorpen 1.1.0 to measure quantum yield (QY - α, [Photosystem II]) of chlorophyll in leaves.

### Tissue chemistry

We selected 3 leaves per species from 5 sampling weeks (22, 26, 32, 39, and 44) to measure percent carbon (C - %) and nitrogen (N - %), for a total of 102 and 104 data points, respectively. We pulverized oven-dried leaves using a SPEX Sample Prep high-throughput grinder. We then weighed 3 mg of pulverized tissue into a 9 mm tin capsule following the sample preparation protocol for the Central Appalachians Stable Isotope Facility (CASIF, University of Maryland Center for Environmental Sciences, MD, USA) and submitted samples for analysis.

To ensure that our measured traits are within the range of globally accepted trait values, we further compared our trait values for LMA, EWT, C, and N with the TRY database (Kattge *et al*., 2020) and visually assessed trait ranges from the Canadian Airborne Biodiversity Observatory database (CABO) (Kothari *et al*., 2023). Our trait comparison in TRY did not include EWT due to insufficient data. Also, we did not compare our QY values with TRY and CABO due to the absence of data.

### Spectral and trait correlation

We assessed correlation in spectra and traits across leaf phenological stages by selecting three weeks (22, 32, and 44) to represent early (young), peak (mature), and late (old/senescent) growing seasons. We tested the significant differences (p < 0.05) between phenological stages (for example, young versus mature and mature versus old/senescent) and squared the differences in correlation to assess how each phenological stage differs from the other. Because carbon and nitrogen were measured on a subset of sampling weeks, trait correlations involving those two traits rest on substantially fewer observations per phenophase than those involving LMA, EWT, and quantum yield, and we interpret them accordingly.

### Fitting models to predict traits from spectra

We used a PLSR modeling approach to develop trait prediction models from the processed spectral data, given its ability to handle high-dimensional and multicollinear data (Wold *et al*., 2001). We used the trimmed reflectance spectra (400-2400 nm) as predictors to build trait models for LMA, EWT, C, N, and QY, and followed established statistical protocols and examples (Serbin *et al*., 2014; Burnett *et al*., 2021a; Kothari *et al*., 2023).

We developed three models, namely: **(1) *all-season* model**, trained on the full spectral and trait data across the growing seasons; **(2) *week-as-covariate* model,** similar to the *all-season* data, but also incorporates the ‘week’ of collection as a covariate alongside the spectral data as a predictor; and **(3) *peak-season* model**, trained only on peak-growing season spectra and trait data. Broadly, these trait models can be grouped into “complete leaf growing season” models encompassing *all-season* and *week-as-covariate* models with 24 weeks of data collection and “models trained on one phenophase” with 10 weeks of data, which covers weeks 28 to 37.

Across all models, we split the dataset into 75% calibration and 25% validation using the createDataPartition function in the caret R package (Kuhn, 2008), balancing the subsampling using the branch information (BranchID) for each individual in the weekly measurements. We tuned the model by fitting PLSR to the calibration subset using the pls R package (Mevik & Wehrens, 2007; Liland *et al*., 2026) with a maximum of 30 latent components. We used cross-validation to estimate model performance for each number of components with the ‘oscorespls’ method and selected the optimal number of components using the ‘onsigma’ method, which chooses the simplest model whose cross-validation root mean square error (RMSE) is within 1 standard error of the minimum error. The optimum number of components varied considerably, from 3 to 21 components (Table S2), which could be attributed to differences in data size across traits. We performed 1000 bootstrap iterations on the calibration dataset to obtain model coefficients. To assess the most important spectral regions for trait prediction, we calculated the variable influence on projection for each trait (Wold *et al*., 2001). We then extracted model coefficients across the three model categories to predict traits—LMA, EWT, C, N, and QY— and used them for downstream analyses.

We assessed the performance of our fitted models—*all-season*, *week-as-covariate*, and *peak-season*— predictions on the 25% validation dataset using four different metrics. We calculated coefficient of determination (R^2^) to measure the proportion of variance in the predicted trait values that is explained in the spectra data; root mean square error (RMSE) to measure the magnitude of error in our model predictions; percentage root mean square error (%RMSE) to calculate the normalized RMSE; and bias (average difference between observations and predictions) to show how likely our model could over- or underestimate the measured traits.

### Assessing model performance over time

We then assessed the performance of our model predictions—*all-season*, *week-as-covariate*, and *peak-season*—on measured traits. We assessed their performance over time by calculating R², RMSE, %RMSE, and bias. We were explicitly interested in how each model predicts traits across phenological stages, including how models trained on one phenophase —*peak-season*—predict early and late season traits. We generated time-series plots using ggplot2 (Wickham, 2016) for predicted and measured traits to display trait temporal dynamics and patterns for each species.

In addition to the above-mentioned trait models, we used widely adopted trait model coefficients (Kothari *et al*., 2023), hereafter Kothari 2023) to estimate the following traits: LMA, EWT, C, N, leaf dry matter content (LDMC), ChlA, ChlB, carotenoids, lignin, cellulose, hemicellulose, soluble fractions, phosphorus, and potassium. We calculated total chlorophyll by summing the chlorophyll predictions of ChlA + ChlB. We assessed the performance of LMA, EWT, C, and N on the measured traits over time, and observed the temporal variation and patterns of other traits across species. We scaled EWT (mm) in (Kothari *et al*., 2023) to g/cm^2^ for uniformity in trait units. We chose to use (Kothari *et al*., 2023) because it allows us to assess multiple traits in addition to the four traits that are central to our study. We further tested the proportion of trait predictions that are mathematically valid and are biologically meaningful for explaining plant function. That is, we quantified trait concentrations that are ≥ 0 and trait percentages that are < 100%.

### Assessing significant differences in traits

We selected three weeks (22, 32, and 44) to represent early, peak, and late growing seasons to test whether the measured traits differ significantly among species. We also tested whether the predicted traits differed significantly from the measured traits by species at different phenological stages. We performed analysis of variance (ANOVA) to test whether there are statistical differences in trait means for each species within the datasets, and a post hoc analysis—Tukey HSD test—for multiple trait comparison to assess which phenophases differed (McHugh, 2011). We marked levels of significance with asterisks (*, **, or ***) at p-values ≤ 0.05, 0.01, and 0.001, respectively.

## RESULTS

### Spectral and trait phenology

Reflectance spectroscopy captured the different patterns in leaf phenological change across taxa (Fig. 1a,b, Fig. S1). The visible region (VIS, 400 - 700 nm) — which is associated with leaf pigments (Curran, 1989) — was more labile across seasons, as indicated by the higher coefficient of variation (CV) of the average weekly spectra. In contrast, the near-infrared (NIR, 700-1100 nm) and short-wave infrared (SWIR, 1100-2400 nm) regions — associated with leaf water, structure, and secondary compounds — showed overall lower CV of the weekly spectra, and are therefore more conserved across leaf phenology (Fig 1b). Different species showed varying degrees of spectral lability across seasons (Fig 1b). For example, the VIS in *R. maximum* showed less temporal lability than in other species, and a similar degree of lability in the water absorption feature (1940 nm) in the SWIR. *A. platanoides* and *R. catawbiense* showed the highest lability in the VIS region.

**Fig. 1.**
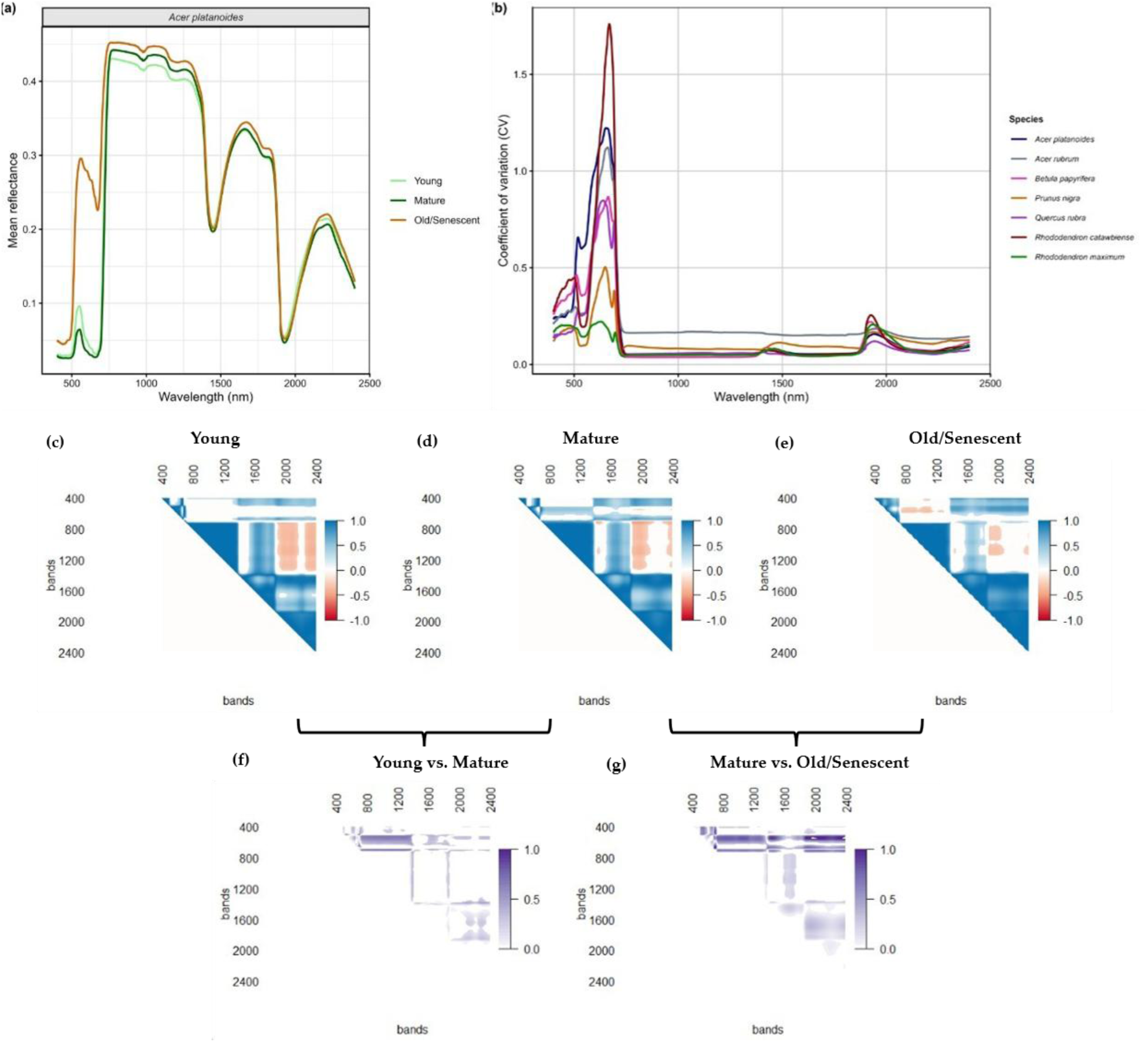
Variation in leaf spectra, coefficient of variation (CV), and correlation across species. (a) Spectral signatures of leaf phenological stages (young, mature, and senescent), which represent three seasons (early, peak, and late season) for the select species. (b) CV across species showing spectral regions that are highly labile (400-700 nm) vs. conserved (700-2400 nm) across the growing season. (c-e) Spectral correlation across phenological stages. (f-g) Difference in spectral correlation between phenological stages. Spectral bands ranged from 400-2400 nm.

Leaf traits — leaf dry mass per area (LMA - kg/m^2^), equivalent water thickness (EWT - g/cm^2^), carbon (C - %), nitrogen (N - %), and quantum yield (QY - α)— varied significantly across phenological stages in most species (Fig. 2a-e, Table S3). ANOVA assessment on mean trait values showed that LMA, C, and QY varied significantly across the growing seasons for all species. N changed significantly with phenology, except for *R. maximum*. Similarly, EWT also varied significantly across seasons, except in *R. maximum* and *Q. rubra*. Trait temporal patterns differed among taxa from early (young) to late (old/senescent) season (Fig. 2). For example, *R. catawbiense* and *R. maximum* experienced a sharp decrease in LMA from early to peak season, followed by a gradual increase from peak to late season. In contrast, most other species’ LMA increased from early to peak season and decreased from peak to late season, while *B. papyrifera*’s LMA was relatively stable over time. EWT increased from early to peak season in *R. maximum* and *P. nigra* but decreased to various degrees in the other species. QY increased from early to peak season but decreased in late season, except for *R. maximum*, which increased in late season. Despite higher uncertainty in C and N estimates, both show an overall bimodal pattern of trait change and decline towards senescence (Fig. 2). In general, both C and N increased in the first few weeks, then decreased towards peak season. In the second half of the growing season, C and N again increased towards early Fall and then decreased towards senescence. Despite seasonal variation in traits, the values are within the global range in TRY and CABO databases (Table S4).

**Fig. 2.**
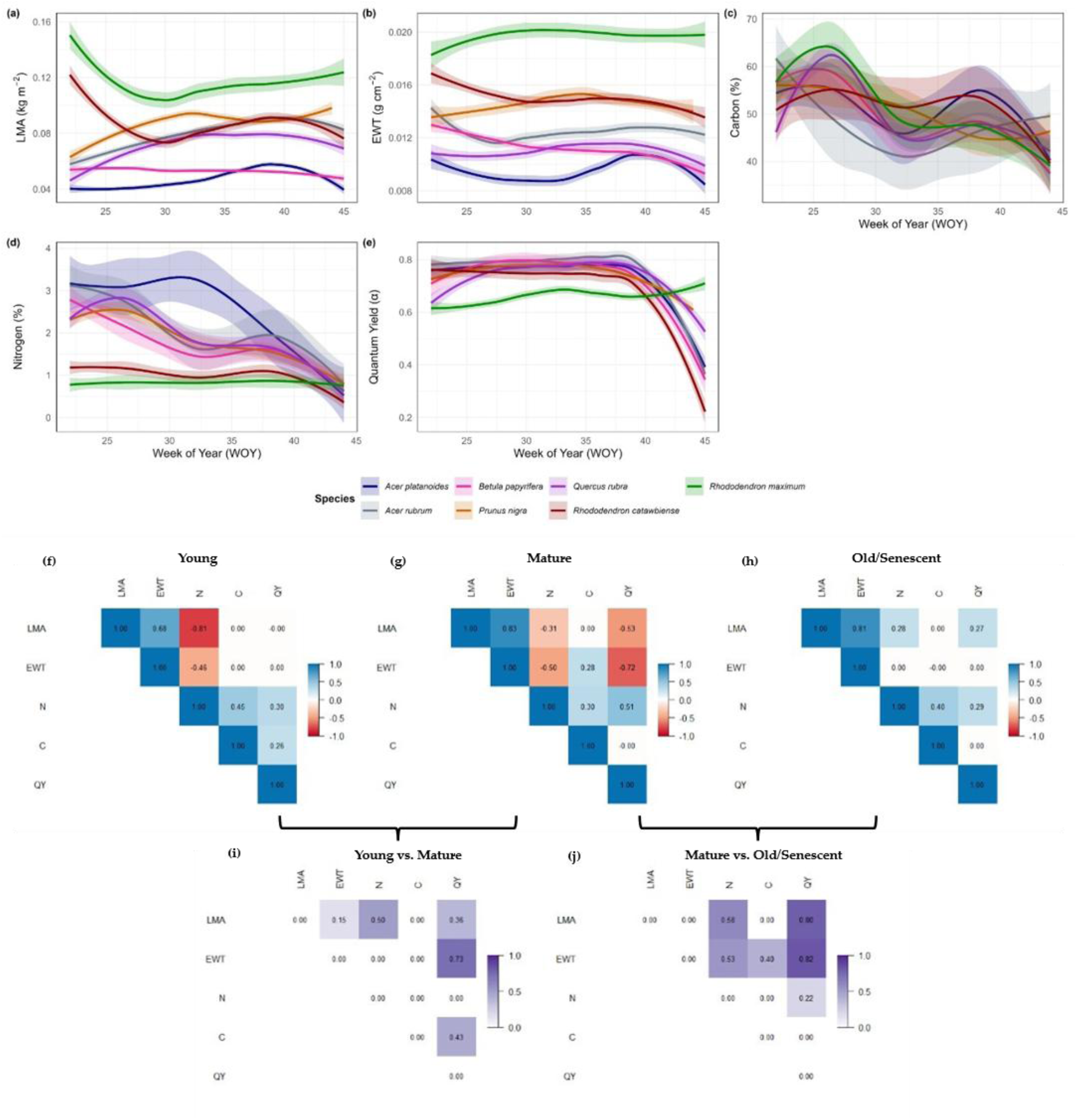
Temporal dynamics of leaf traits and their correlation across species. (a) LMA, leaf mass per area (kg/m^2^), (b) EWT, equivalent water thickness (g/cm^2^), (c) Carbon (%), (d) Nitrogen (%), and (e) QY, Quantum Yield (α). (f-h) Trait correlation across leaf phenological stages, (i-j) Difference in trait correlation between leaf phenological stages. Solid lines are the smooth curve from LOESS (locally estimated scatterplot smoothing) regression, and the shaded area depicts the 95% confidence interval of the smooth curve. Weekly measurement (22-45) covers from May 31st (shortly after leaf emergence) to November 6th, 2023 (old/senescent).

### Spectral and trait correlation

Correlation in spectral and traits varied across leaf phenological stages and were significantly different (Figs. 1,2). Across leaf phenological stages, the NIR—SWIR regions maintained a negative correlation, while the correlation of VIS—NIR varied from zero to positive to negative (Fig. 1c-e). Between phenological stages (young versus mature and mature versus old/senescent), the differences in spectral correlation were significant, particularly for VIS—NIR and VIS—SWIR regions (Fig. 1f,g).

The strength and direction of trait correlations varied across phenophases, for example, while LMA—N were negatively correlated in young and mature leaf stages, both traits were slightly positively correlated in old/senescent leaves (Fig. 2f-h). This variation in trait correlation across leaf phenological stages was significantly different when assessed between phenophases (Fig. 2i,j). Other trait correlations, such as LMA—QY and EWT—QY, also showed significant differences from young to mature and to old/senescent leaves. In contrast to this variation, some traits maintained positive correlations across leaf phenological stages, for example, LMA—EWT, C—N, and N—QY (Fig. 2f-h).

### Assessing model performance across leaf phenology

We show that PLSR can build models for predicting traits from reflectance spectra, despite variation in data sizes and predictor variables. Performance metrics for the three fitting models—*all-season*, *week-as-covariate,* and *peak-season*—show high accuracy for LMA and EWT (R^2^ = 0.81 - 0.94) and low to intermediate accuracy for C, N, and QY (R^2^ = 0.11 - 0.66) (Fig. S2, S3; Table S4). The VIP metrics across the three models show the critical spectral regions for predicting the traits (Figs. S4; Results S1).

We assessed the performance of four models trained using different phenological stages. Model classes—either complete leaf phenology or models trained on one phenophase—show similar accuracies. For example, *all-season* and *week-as-covariate* models show high accuracy for LMA and EWT (R^2^ = 0.85 - 0.93, %RMSE = 4.48 - 4.77), intermediate accuracy for N and QY (R^2^ = 0.6 - 0.66, %RMSE = 9.13 - 10.71), and low accuracy for C (R^2^ = 0.08 - 0.26, %RMSE = 17.83 - 19.92) (Fig. 3, S5, S9; Table 1, Table S5). Training set size differed markedly among traits (see Methods), which bears on the comparison across traits. *Peak-season* and *Kothari 2023* models showed low accuracy for all traits, for example, LMA (R^2^ = 0.03 - 0.25, %RMSE = 39.05 - 89.83), EWT (R^2^ = 0.15 - 0.25, %RMSE = 16.34 - 27.28), C (R^2^ = 0.00, %RMSE = 22.59 - 23.31), N (R^2^ = 0.11 - 0.14, %RMSE = 50.00 - 89.88), and QY (R^2^ = 0.23, %RMSE = 49.14) (Fig. 3; Fig. S6, S9; Table 1). Predictions and model performance metrics for the *week-as-covariate* model (Figs. S7-S9, Table S5), which included time (week of collection) as a covariate to help explain temporal patterns, were statistically indistinguishable from those of the *all-season* model (and therefore only shown in the supplement). This result is noteworthy because it shows that spectra alone can capture the leaf phenological dynamics.

To further characterize model performance, we ran a linear regression between predicted and measured traits for each model. The slope of the regression in unbiased models should overlap 1 (i.e., coincide with the 1:1 line). Across our model categories, we observed a significant difference in slope between complete leaf phenology models and models trained on one phenophase for all traits (Fig. 3), and the 95% confidence interval of the slope for complete leaf phenology models overlaps with the 1:1 regression line. However, in models trained on one phenophase, the 95% confidence interval of the slope does not overlap with the 1:1 regression line. Overall, models trained with complete leaf phenology data performed better than those trained with one phenophase data.

**Fig. 3.**
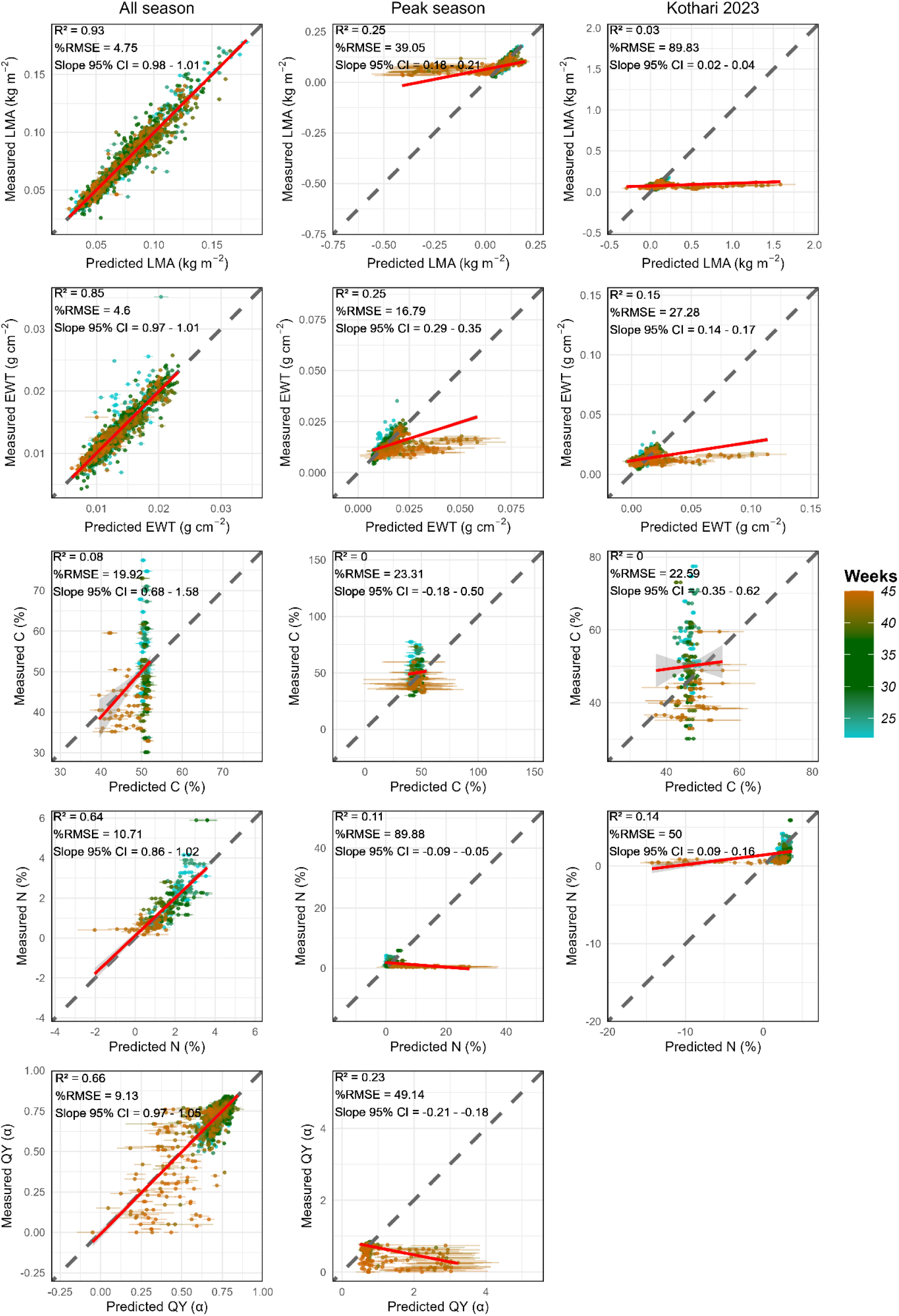
Validation results for predicted traits—LMA, EWT, C, N, and QY—from model coefficients of *all-season*, *peak-season*, and *Kothari 2023* assessed on measured traits by week. Weeks = Week of Year (WOY). Weekly measurement (22-45) covers from May 31st (shortly after leaf emergence) to November 6th, 2023 (old/senescence). The dark grey dashed line in each panel is the 1:1 line, and the red line is the fit of a linear regression between predicted and measured traits. The Slope 95% confidence interval (CI) shows how different the regression line is from the 1:1 line. *Kothari 2023* model coefficients for trait prediction were obtained from(Kothari *et al*., 2023). The validation results for *week-as-covariate* are shown in the supplementary information (Fig. S7) and species validation results (Fig. S5).

**Table 1.** Summary statistics for model performances over time on measured traits.

|  | Complete leaf phenology model |  |  |  | Models trained on one phenophase |  |  |  |  |  |  |  |
| --- | --- | --- | --- | --- | --- | --- | --- | --- | --- | --- | --- | --- |
|  | All-season |  |  |  | Peak-season |  |  |  | Kothari 2023 |  |  |  |
| Trait | *n* | R<sup>2</sup> | %RMSE | BIAS | *n* | R<sup>2</sup> | %RMSE | BIAS | *n* | R<sup>2</sup> | %RMSE | BIAS |
| Leaf mass per area (kg/m<sup>2</sup>) | 1500 | 0.93 | 4.75 | 0.00 | 1500 | 0.25 | 39.05 | 0.01 | 1500 | 0.03 | 89.83 | -0.03 |
| Equivalent water thickness (g/cm<sup>2</sup>) | 1500 | 0.85 | 4.60 | 0.00 | 1500 | 0.25 | 16.79 | -0.00 | 1500 | 0.15 | 27.28 | 0.001 |
| Carbon (%) | 304 | 0.08 | 19.92 | 0.11 | 304 | 0.00 | 23.31 | 4.27 | 304 | 0.00 | 22.59 | 3.80 |
| Nitrogen (%) | 310 | 0.64 | 10.71 | 0.03 | 310 | 0.11 | 89.88 | -1.52 | 310 | 0.14 | 50.00 | -0.15 |
| Quantum yield ($\alpha$) | 1500 | 0.66 | 9.13 | 0.00 | 1500 | 0.23 | 49.14 | -0.09 | NA | NA | NA | NA |

NA refers to trait information not available.

We observed that the temporal variation and patterns between measured and predicted traits are similar for complete leaf phenology models, but different for models trained on one phenophase, particularly in the late season (Fig. 4, S8, S10-12). Across all models, trait values were similar to the measured dataset in early and peak seasons. In the late season, models trained on one phenophase either over- or under-predicted traits with negative values across taxa (Fig. 4). These predictions for LMA, EWT, C, N, and QY produced trait values that are beyond the measured range and may not reflect ideal plant functional and ecosystem properties. This shows that models trained on one phenophase may not be ideal for temporal analysis of trait phenology because they are forced to predict traits outside their training data range. Predictions from models trained with complete leaf phenology data were more similar to the measured traits and TRY dataset compared to those from models trained on one phenophase, which had either high or low predictions with negative values for LMA, EWT, QY, and N, except for C.

**Fig. 4.**
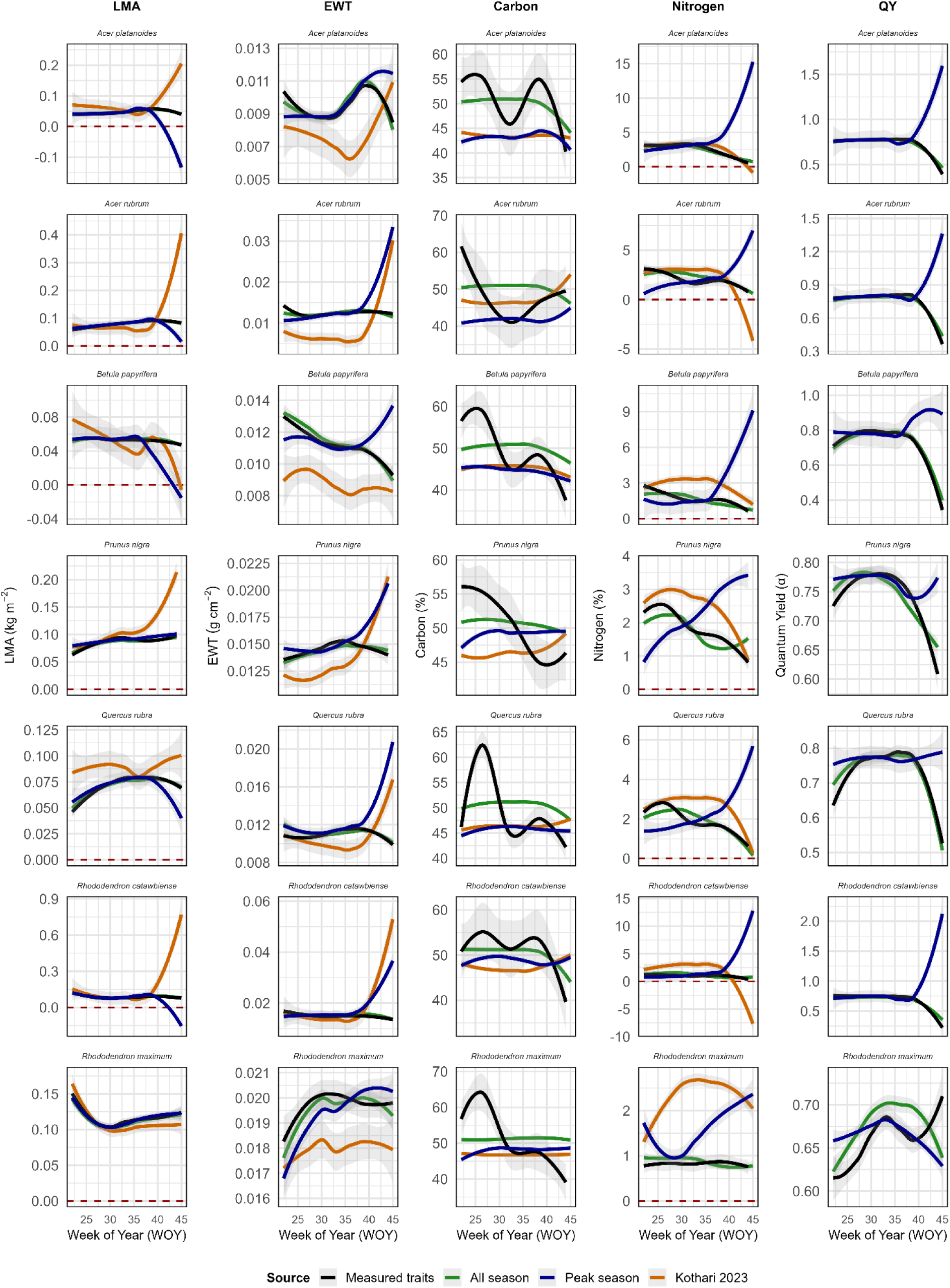
Temporal trends in LMA, EWT, Carbon, Nitrogen, and QY across trait datasets—measured vs predicted traits. Leaf mass per area (LMA); equivalent water thickness (EWT); quantum yield (QY). Kothari 2023, model coefficient from(Kothari *et al*., 2023). Solid lines are the smooth curve from LOESS (locally estimated scatterplot smoothing) method, and the shaded area depicts the 95% confidence interval of the smooth curve. Weekly measurement (22-45) covers from May 31st (shortly after leaf emergence) to November 6th, 2023 (senescence). Predictions from *all-season*, *peak-season*, and *Kothari 2023* are from trait model coefficients that cover different stages of leaf phenology. Trait temporal trends for *week-as-covariate* model are in Fig. S8.

In addition, we used model coefficients from (Kothari *et al*., 2023) to estimate traits we did not measure, including pigment concentrations, carbon-soluble fractions, and micronutrients. We assessed the quality of these predictions very conservatively by asking if they were mathematically viable, that is, if concentrations were greater than 0 (≥ 0) and if percentages fell between 0 and 100 % (Figs. 5, S10 - S12). We show that the proportion of mathematically reasonable trait predictions is high in peak growing season, with 98-100%, but often low in late season (senescence), with 33-97% (Fig. 5). There was considerable variation in how unrealistic trait estimates for late season were. For example, only 3% of the estimates for K were not mathematically reasonable, whereas 67% of the hemicellulose concentration estimates were unrealistic. Still, ⅓ of the estimates were unreasonable for 6 out of 9 traits (Fig. 5).

We tested the significant differences between predicted and measured traits across early, peak, and late seasons. We show that in *all-season* and *week-as-covariate* models, there are no significant differences from measured traits for LMA and EWT across all species and leaf phenophases (Table S6). In models trained on one phenophase, such as *peak-season* and *Kothari 2023*, predicted traits differed significantly from measured traits, particularly in the early (young) and late (old/senescent) seasons (Table S7). However, the significance levels differed across species and traits. For example, all species except *R. maximum* showed significant differences in LMA and C at early and late seasons. Across leaf phenophases, our results show that trait predictions were stronger in peak season. This variation in significant differences across trait models and seasons underscores the importance of capturing variation in leaf phenophases.

**Fig. 5.**
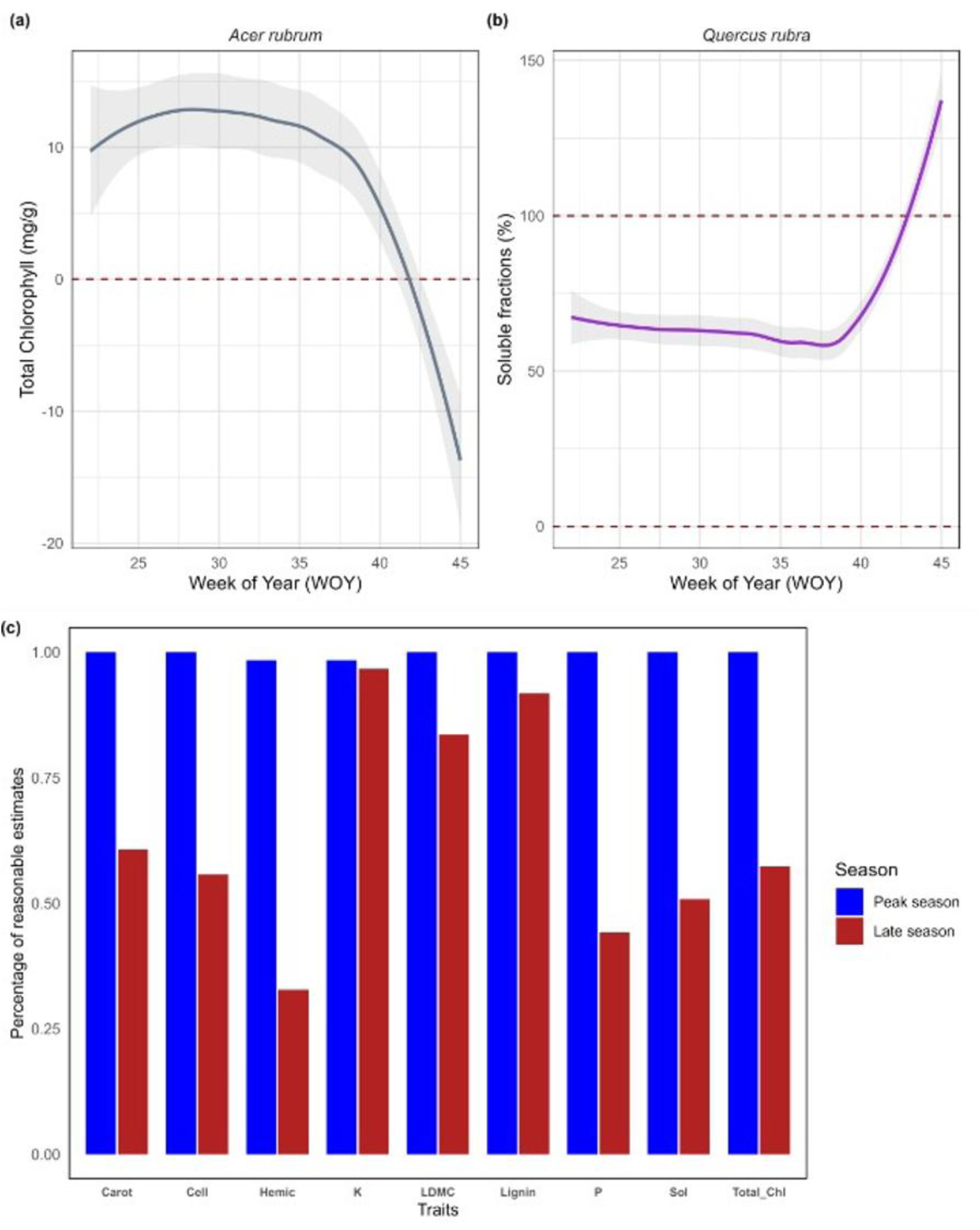
Predictions of additional traits from *Kothari 2023* from May 31st (shortly after leaf emergence, week 22) to November 6th, 2023 (senescence, week 45). (a) and (b) Temporal trends in predicted Total Chlorophyll and Soluble fractions, showing reasonable estimates over most of leaf phenology but unreasonable trait estimates in late season (concentrations must be > 0 and percentages fall between 0 and 100). (c) Percentage of peak and late season trait estimates that are reasonable for Carotenoid (Carot); Cellulose (Cell); Hemicellulose (Hemic); Potassium (K); Leaf dry matter content (LDMC); Lignin; Phosphorus (P); Soluble (Sol); Total Chlorophyll (Total_Chl).

## DISCUSSION

Models trained on complete phenological data tracked trait dynamics accurately throughout the growing season, confirming the strong potential of reflectance spectroscopy for temporal trait assessment when training data are representative (Figs 3, 4). In contrast, models trained on one phenophase — including a widely used published model (Kothari *et al*., 2023) — not only underperformed quantitatively (Figs 3, 4) but produced biologically unrealistic predictions that may systematically bias ecological inference (Fig. 5). Both trait and spectral correlation structure also shifted across phenophases (Figs. 1,2). We first interpret the temporal patterns in spectra, traits, and their correlations. Then we examine how and where the models failed and consider what the shift in correlation structure can and cannot explain about that failure. Finally, we discuss what a phenologically explicit view of plant function offers ecology and evolution.

### Spectral and trait phenology

The differential temporal lability of spectral regions across phenology reflects the distinct responsiveness of their underlying leaf properties to seasonal change (Yang *et al*., 2016; Ji *et al*., 2026). The VIS region — associated with pigment pools that turn over rapidly during leaf development and senescence (Curran, 1989; Ustin *et al*., 2009; Ustin & Jacquemoud, 2020) — showed high CV across the season, consistent with the degradation of chlorophyll (Hörtensteiner, 2006; Tanaka & Ito, 2025), the decline in photosynthetic activity (Luo *et al*., 2017; Palm *et al*., 2022; Ji *et al*., 2026), and the synthesis of anthocyanins and carotenoids during senescence (Zhao *et al*., 2021). The NIR and SWIR, which capture structural features, water content, and secondary compounds (Couture *et al*., 2016), showed lower CV, consistent with the slower turnover of cell wall and structural constituents. The two evergreen *Rhododendron* species, however, had contrasting spectral responses: while *R. maximum* showed low lability in the VIS, *R. catawbiense* had the highest VIS lability of any species, consistent with our field observations of its leaves turning red, turning yellow, or remaining green at the end of the growing season. Changes in CV at 1400 and 1940 nm correspond to water absorption features (Curran, 1989), indicating that water-related spectral signals also shift with seasonal changes in plant water status.

The contrasting trait trajectories we observed across species are consistent with distinct developmental patterns and functional strategies rather than variation around a seasonal mean. In deciduous species such as *A. platanoides*, *A. rubrum*, and *Q. rubra*, LMA increased from early to peak season and declined during senescence, a pattern consistent with the accumulation and subsequent reabsorption of structural carbon compounds (Fajardo & Siefert, 2016). In contrast, the evergreen *Rhododendron* species showed continuous increases in LMA from peak to late season, likely reflecting ongoing structural investment in long-lived leaves. Similarly, the stability of N and QY in *R. maximum* relative to the broad late-season decline in the deciduous taxa suggests that evergreen species may maintain photosynthetic nitrogen pools across seasons, whereas the decrease in N toward senescence in deciduous species is consistent with nitrogen reabsorption before abscission, a well-documented pattern in temperate deciduous taxa (McKown *et al*., 2013; Fajardo & Siefert, 2016). These species-specific temporal profiles are not simply variation around a peak-season value, but instead seem to encode life-history information, specifically contrasting strategies of leaf longevity, resource investment, and reabsorption, that peak-season snapshots systematically exclude.

### The phenology of correlation structure

Spectral correlation structure was not stationary across phenophases (Fig. 1c-g). The NIR–SWIR regions maintained a negative correlation across all three phenophases, reflecting the joint dependence of both regions on leaf water status and structure (Slaton *et al*., 2001; Ustin & Jacquemoud, 2020). In contrast, the VIS–NIR correlation varied from near zero to positive to negative, which is consistent with a decline in chlorophyll and potentially anatomical changes, in particular in the spacing of the spongy mesophyll as leaves aged (Curran, 1989; Vogelmann, 1993; Ustin *et al*., 2009; Ustin & Jacquemoud, 2020). The differences between phenophases were significant and concentrated in the VIS–NIR and VIS–SWIR (Fig. 1f,g), the same regions in which spectral lability was highest (Fig. 1b).

Likewise, trait correlation structure also had a phenology, and its components were not equally labile (Fig. 2f-j). Some relationships held across all three phenophases — LMA–EWT, C–N, and N–QY remained positively correlated from young to senescent leaves — whereas others changed in strength or sign. The sharpest case is LMA–N, which was negatively correlated in young and mature leaves, as the leaf economic spectrum predicts (Wright *et al*., 2004), but weakly positive in senescent leaves, since both traits decline together as plants shift from acquisition to reabsorption. Correlations between structural and photosynthetic traits, LMA–QY and EWT–QY, also differed significantly between phenophases (Fig. 2i,j). Trait correlation structure is known to shift with ontogeny (Damián *et al*., 2018), between years (Pahadi *et al*., 2024), between sampling dates (Mason *et al*., 2020), and between coarse seasons (Rao *et al*., 2023). Our contribution resolves that shift within a single leaf cohort across the phenophases of one growing season, and shows that the integration described by a peak-season correlation matrix is a property of that phenophase rather than of the species. We acknowledge, however, that correlations involving C and N are based on far fewer observations per phenophase than those involving LMA, EWT, and QY (see Methods), and the magnitude of their change should be interpreted accordingly.

### Model performance across phenology

Complete leaf phenology models achieved high accuracy for LMA and EWT, intermediate accuracy for N and QY, and low accuracy for C (Fig. 3, Table 1). The lower accuracy for C and N across all model types likely reflects the smaller training sets available for those traits (see Methods). Moreover, carbon may additionally be an intrinsically peripheral trait since it is present in nearly all organic leaf constituents and therefore lacks the distinct clear absorption features that water and chlorophyll possess. Advancing the generality of spectral trait models across systems remains a key challenge (Kothari *et al*., 2023), and previous work has shown that such models may not generalize across sites, biomes, or species assemblages when calibration data do not represent the target conditions (Kothari *et al*., 2023; Wang *et al*., 2023). Our results extend that transferability problem to the temporal dimension. Even within a single site and species set, models trained on one phenophase performed poorly across all traits (R² ≤ 0.25; Fig. 3, Table 1) and systematically over- or under-predicted traits in the late season (Fig. 4), while their regression slopes departed significantly from the 1:1 line that the complete-phenology models matched (Fig. 3). Adding collection week as a covariate did not improve the model performance (Figs S7-S9, Table S5), which indicates that spectra alone carry the phenological information. That is, the temporal coverage of the training data, not an explicit time term, is what determines model performance.

Models trained on one phenophase — including the widely used (Kothari *et al*., 2023) model — did not simply lose precision outside their training window: they produced biologically unreasonable predictions (Fig. 5, S10-S12). These included negative concentrations for total chlorophyll, carotenoids, cellulose, hemicellulose, and phosphorus; soluble fractions exceeding 100%; and predicted increases in N during senescence, when measurements and the literature demonstrate a decline (Escudero & Mediavilla, 2003). Such errors are consequential because a researcher relying on these predictions may “learn” that foliar N rises toward senescence (Fig. 3) and draw ecological conclusions from an artifact. We are not implying by any means that the Kothari 2023 model is at fault. We think the problem is that we stretched the model to time windows it was not built to cover. Our results support this interpretation, since 98–100% of the predictions from the Kothari 2023 model fell within mathematically valid and biologically meaningful bounds for all traits, whereas only 33–97% did so in the late season (Fig. 5).

When direct trait measurements are unavailable for validation — as may be the case in applied settings — our results suggest that mathematical and biological bounds can serve as practical, if very conservative, post-hoc diagnostic criteria for assessing models. Concentrations must be non-negative, percentages must not exceed 100%, and predicted temporal trends should not contradict well-established physiological patterns. Applying these criteria revealed that models trained on one phenophase frequently produced values outside valid bounds in the late season, whereas complete leaf phenology models did not. We propose that researchers routinely apply such bounds-checking when using spectral models outside their original calibration conditions, since it provides a minimum safeguard against biologically misleading inference even in the absence of ground truth data.

### Why phenologically narrow models fail

Empirical spectral models leverage covariance, and both covariances depend on shifted across phenophases. PLSR uses the correlation among spectral bands to reduce the spectrum to a few latent components (Wold *et al*., 2001), and it recovers traits partly through their covariance with traits that have direct absorption features (Chadwick & Asner, 2016; Nunes *et al*., 2017), under which peripheral traits are predicted through their correlations with core traits rather than through optical signatures of their own (Kothari & Schweiger, 2022). As a consequence, a model calibrated at peak season, encodes the peak-season spectral and trait correlation structure alongside the spectral–trait relationships themselves. When applied to senescent leaves, the model meets spectra whose VIS has shifted enough to alter band covariance (Fig. 1f,g), forcing it to extrapolate into regions of latent-variable space where the calibrated relationships were never fitted, and it applies trait relationships that have weakened or changed sign (Fig. 2i,j). (Kothari *et al*., 2023) state the risk explicitly for their own models: if trait covariance patterns vary among datasets, overfitting to the training data could cause such models to produce inaccurate trait estimates from new spectral data. Our data document the two premises of this argument — correlation structure changed, and models failed where it changed — but we did not test the link between them. Our design does not isolate covariance shift from the alternatives considered next, and the uniformity of failure across core and peripheral traits (Table 1) suggests the data cannot rank traits by their dependence on covariance.

Several mechanisms may contribute to late-season failure, and they are not mutually exclusive. Beyond a shift in covariance structure, models applied to senescent leaves must extrapolate beyond the range of trait values on which they were trained, and range extrapolation is a recognized failure mode of empirical spectral models. Measurement uncertainty could be higher in senescing tissue. Distinguishing these would require training sets that hold trait range constant while varying correlation structure, or the reverse — a design our sampling was not built to support. We raise the alternatives because the correlation explanation is the one our data motivate, not because they are the only possibilities. Whatever the cause, PLSR appears especially exposed to it: it has worse transferability than more mechanistic models (Kothari *et al*., 2023; Wang *et al*., 2023), is prone to overfitting, and yields uncertainty estimates poorly calibrated to true prediction error, whereas approaches such as Bayesian model averaging that account for model uncertainty and perform variable selection reduce overfitting and improve generalization (Zhao *et al*., 2013; Ji *et al*., 2024). Such approaches may prove more robust when models must generalize across phenological conditions absent from the training data.

### A dynamic view of plant function

Plant functional traits encode ecological and evolutionary strategies across scales from individual physiology to macroevolutionary diversification (Wright *et al*., 2004; Violle *et al*., 2007; Cornwell *et al*., 2014; Díaz *et al*., 2016), and leaf reflectance can estimate key traits (Serbin *et al*., 2014; Kothari *et al*., 2023), recover phylogenetic signal in functional variation (Meireles *et al*., 2020; Stasinski *et al*., 2021), and enable biodiversity monitoring at landscape scales (Turner *et al*., 2003; Aguirre-Gutiérrez *et al*., 2025). That potential has been realized primarily under frameworks that treat plant function as adequately captured by a single temporal snapshot, and our results challenge this assumption. Established trait-based frameworks, such as the leaf economic spectrum (Wright *et al*., 2004), describe correlations and trade-offs among traits at one point in time and are therefore a single-time-point projection of a trait space through which each species moves over the season. In at least some species, those seasonal trajectories were distinctive enough to suggest that phenological profiles constitute a dimension of functional differentiation among taxa rather than a uniform response to shared environmental cues (Fig. 4; Cope *et al*., 2022): species occupying similar LMA positions at peak season diverged in their LMA trajectories, and trade-offs between LMA and other traits changed over the season (Fig. 2). Incorporating leaf phenology into functional assessments is therefore necessary not only to improve trait estimation but to capture the rate and direction of seasonal change — dimensions of plant strategy that static measurements cannot represent (Violle *et al*., 2012; Reich, 2014; Siefert *et al*., 2015; Cleland & Wolkovich, 2024).

A dynamic view of plant function has direct consequences for community ecology and for evolutionary biology. Because trait differences underlie both the fitness differences that drive competitive exclusion and the stabilizing niche differences that promote coexistence (Mayfield & Levine, 2010; Kraft *et al*., 2015), species-specific seasonal trajectories imply that the strength, and potentially the sign, of interactions among co-occurring species change across the growing season, consistent with the demonstration by Rudolf, (2019) that coexistence conditions depend strongly on differences in relative phenology. Research in plant genetics, adaptation, and macroevolution already represents phenotypic variation as reaction norms expressed across environmental gradients (Gomulkiewicz & Kirkpatrick, 1992; Sultan, 2000), and phenologically capable spectral models offer a tractable way to estimate such norms across the season at the frequency phenology demands. The seasonal lability of traits, and of their integration, may in turn be modeled as a trait in its own right at different phylogenetic scales (Meireles *et al*., 2020; Stasinski *et al*., 2021).

From a methodological perspective, we suggest that researchers building spectral trait models treat the temporal coverage of training data as a first-order design criterion, on par with taxonomic and functional breadth. Complete-phenology models substantially outperformed peak-season alternatives here, while the minimum phenological sampling required to capture the key temporal dynamics of spectra and traits remains to be established. We have shown that PLSR can recover trait dynamics when its training data are representative, whereas its documented shortcomings under domain shift (Zhao *et al*., 2013; Ji *et al*., 2024) argue for continued development of physics-based and Bayesian alternatives that may generalize better to phenophases absent from calibration.

Three limitations bound our conclusions. Our sampling covers a single site, a single growing season, and seven species, so we cannot say how far the patterns we describe in trait trajectories, in correlation structure, or in the divergence between peak-season and late-season spectral feature space generalize to other biomes, growth forms, or climates. Carbon and nitrogen were measured at five sampling weeks against twenty-four for LMA, EWT, and quantum yield, which constrains both the models fitted to those traits and the precision of the correlations we estimate for them at each phenophase. Correlation structure was assessed at three representative weeks rather than continuously, so we characterize its phenology as a contrast among phenophases rather than as a full trajectory. Despite these shortcomings, we are confident that they do not invalidate our findings.

## CONCLUSION

We demonstrate that leaf reflectance spectroscopy can reliably capture the temporal dynamics of functional traits across the full growing season, but only when trait prediction models are trained on phenologically representative data. Models trained on one phenophase — including widely used published coefficients — fail not only quantitatively but qualitatively when applied outside the phenological window of their training data, producing trait estimates that can mislead ecological inference in ways that are not always detectable without direct validation.

Trait integration itself has a phenology: the correlation structure among traits, and among spectral bands, differed across phenophases, and a model calibrated in one phenophase carries that phenophase with it. Whether the shift in correlation structure is the major cause of model failure or one of several contributors remains to be tested. By demonstrating that models trained on complete phenological data substantially outperform peak-season alternatives, our study provides both a cautionary tale and a path forward. Developing a phenologically robust modeling framework for spectroscopy and plant function will be essential for advancing our understanding of plant function at global scales.

## ACKNOWLEDGEMENTS

This study was conducted in the University of Maine, Orono campus, located on Marsh Island in the homeland of the Penobscot and the Wabanaki Tribal Nations. We thank Brian McGill for statistical suggestions and the MMRRG lab group for helpful comments. JEM was supported by the National Science Foundation awards DBI 2021898 and DEB-2442433. The authors thank the editor and the anonymous reviewers for their comments that helped shape this manuscript.

## AUTHOR CONTRIBUTIONS

CON and JEM conceptualized the research; CON collected the traits and spectral data; CON and JEM performed the data analyses, visualization, developed the models, and interpreted the results; CON and JEM wrote the initial draft and final version of the manuscript.

## DATA AVAILABILITY

All spectra and traits used in model calibration and validation are available on Figshare (Nichodemus & Meireles, (2026) (10.6084/m9.figshare.32060352)). R scripts generated for spectral analysis and modeling in this study are available on GitHub (https://github.com/Cornelius-Nic/Spectra_Phenology).

## SUPPORTING INFORMATION

**Fig. S1.**
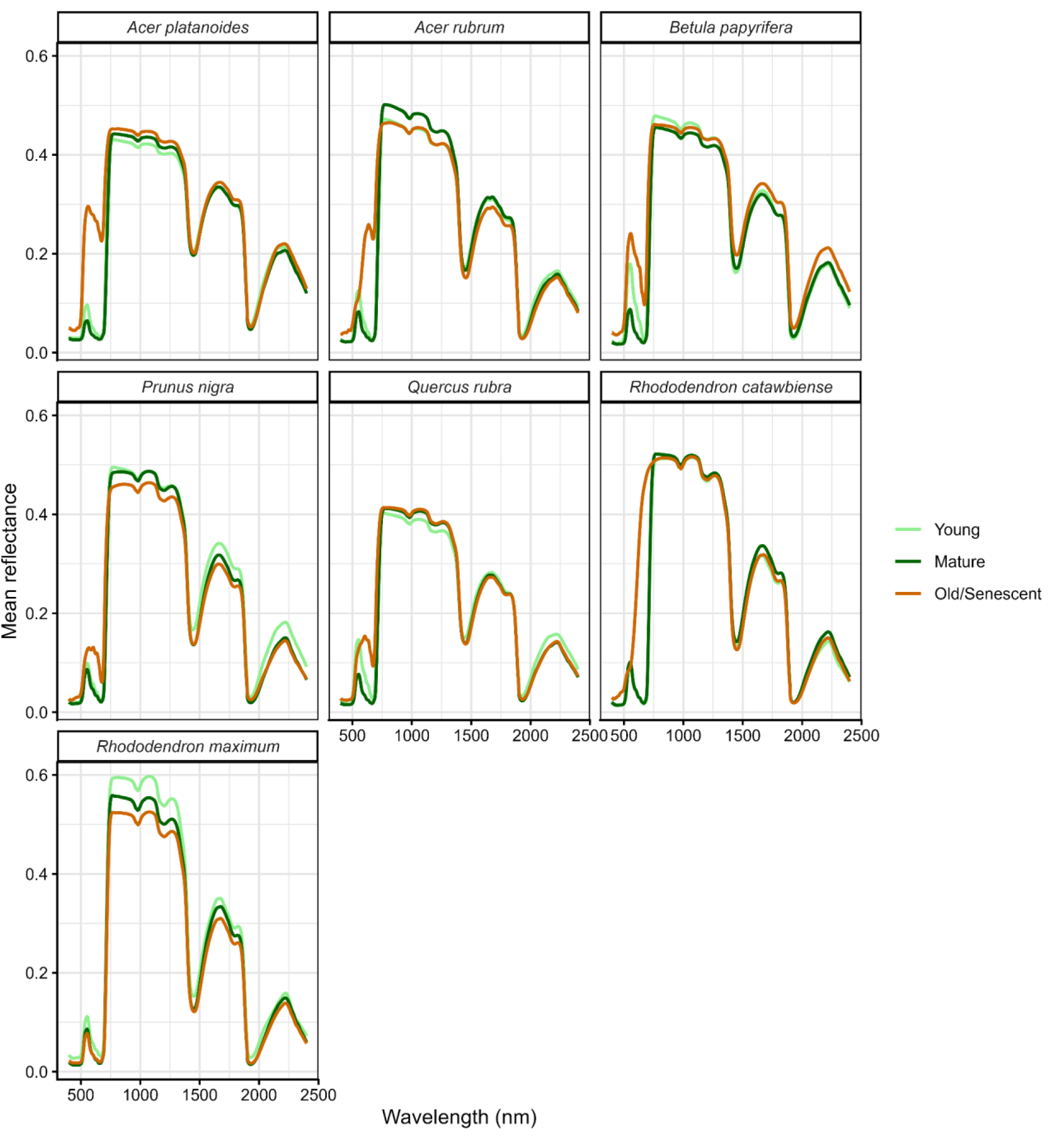
Fresh leaf spectral signatures across taxa for different leaf phenological stages. Colors denote stages in leaf phenology, where young = early season, mature = peak season, and old/senescence = late season.

**Fig. S2.**
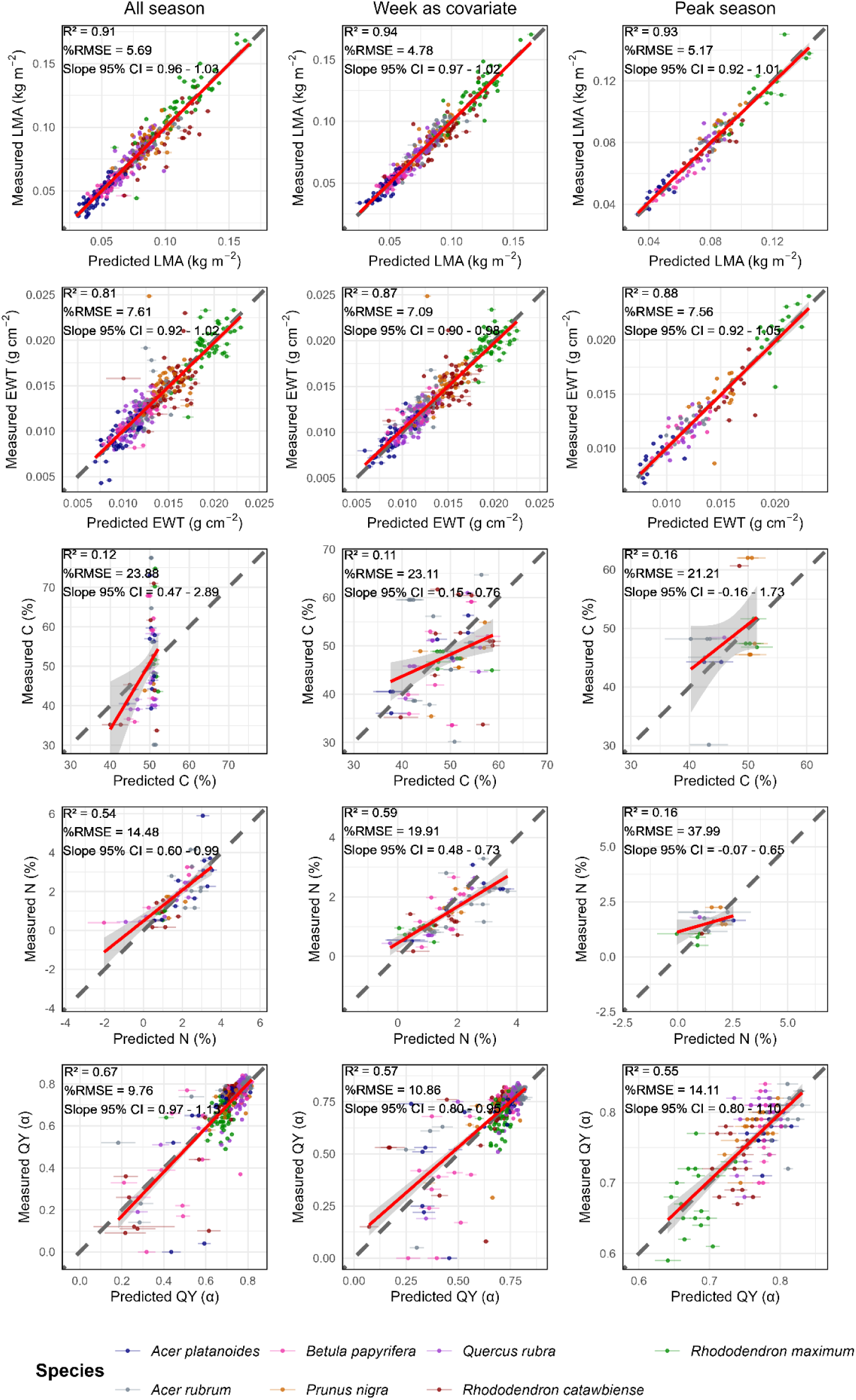
Validation results for the three models—*all-season*, *week-as-covariate*, and *peak-season—*fitting. The dark grey dashed line in each panel is the 1:1 line. Leaf mass per area (LMA); Equivalent water thickness (EWT); Carbon (C); Nitrogen (N); Quantum Yield (QY). The 95% confidence interval (CI) for the slope shows how significantly different the regression line is from the 1:1 line.

**Fig. S3.**
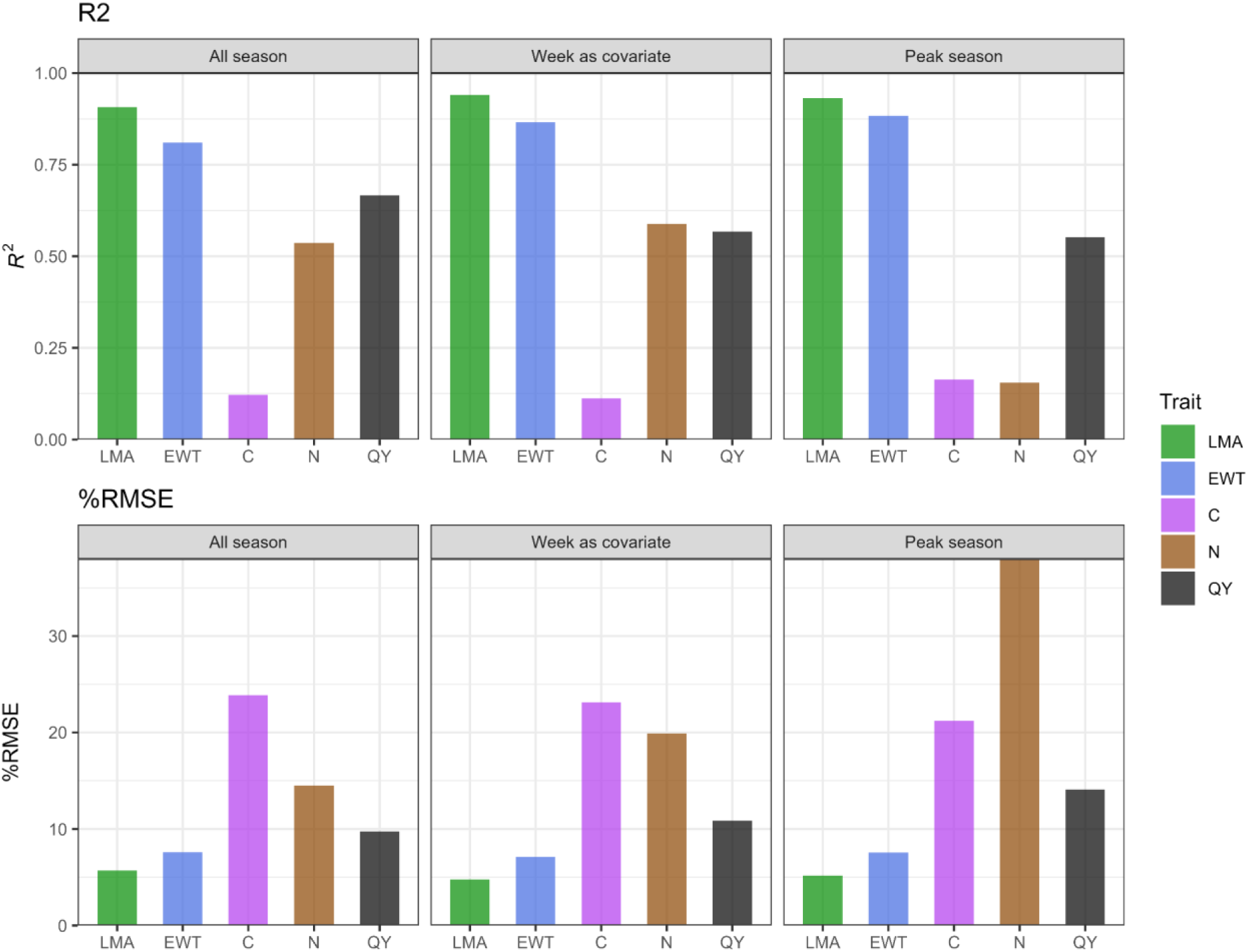
Comparison of model fitting performances. Leaf mass per area (LMA); Equivalent water thickness (EWT); Carbon (C); Nitrogen (N); Quantum Yield (QY).

**Fig. S4.**
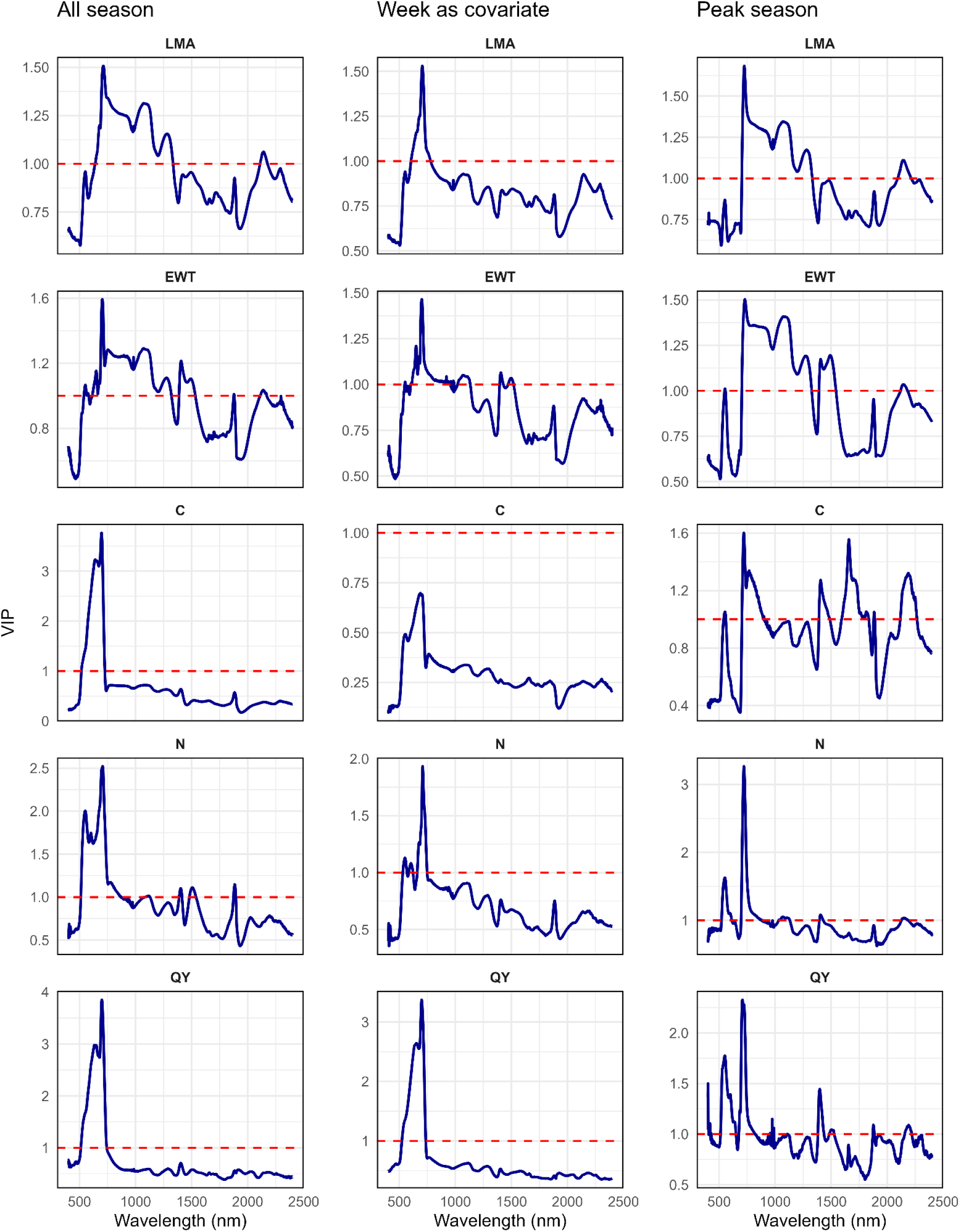
Variable influence on projection (VIP) for *all-season, week-as-covariate*, and *peak-season* models. Leaf mass per area (LMA); Equivalent water thickness (EWT); Carbon (C); Nitrogen (N); Quantum Yield (QY).

**Fig. S5.**
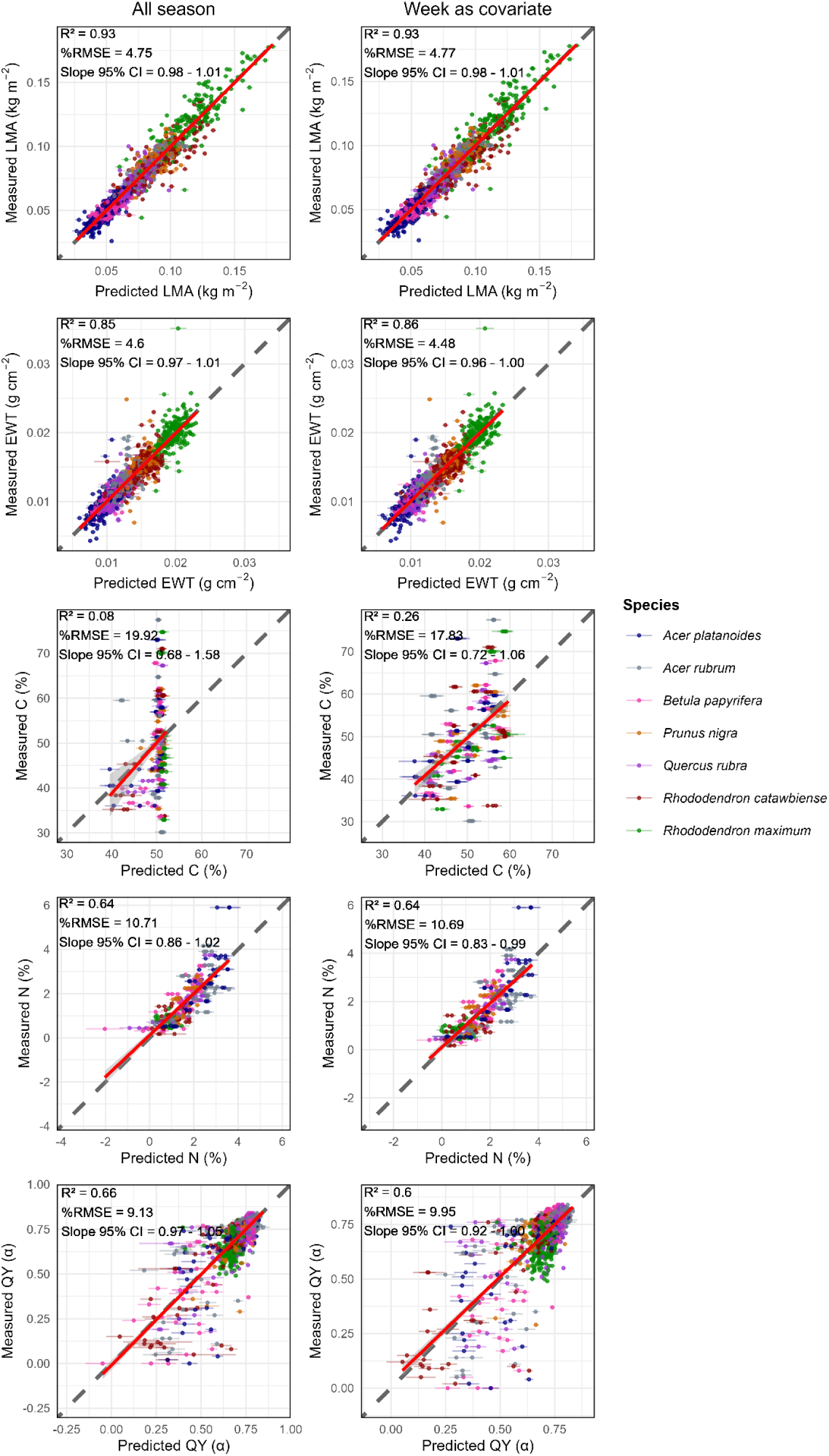
Validation results for complete leaf phenology model coefficients assessed on measured traits by species. The dark grey dashed line in each panel is the 1:1 line. The 95% confidence interval (CI) for the slope shows how significantly different the regression line is from the 1:1 line. Leaf mass per area (LMA); Equivalent water thickness (EWT); Carbon (C); Nitrogen (N); Quantum Yield (QY).

**Fig. S6.**
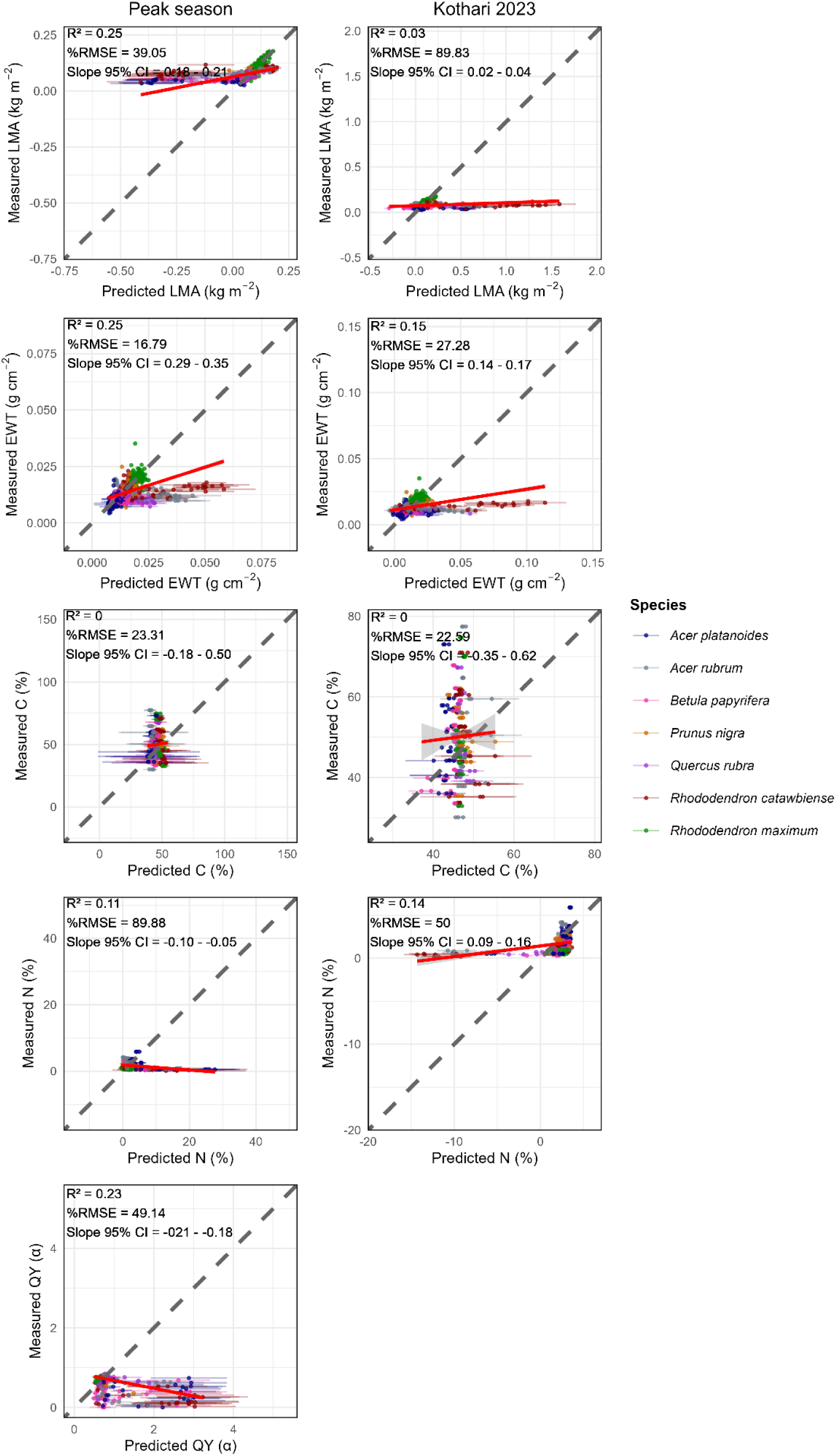
Validation results for models trained on one phenophase coefficient assessed on measured traits by species. The dark grey dashed line in each panel is the 1:1 line. The 95% confidence interval (CI) for the slope shows how significantly different the regression line is from the 1:1 line. Leaf mass per area (LMA); Equivalent water thickness (EWT); Carbon (C); Nitrogen (N); Quantum Yield (QY). *Kothari 2023* model coefficients for trait prediction were obtained from (Kothari *et al*., 2023).

**Fig. S7.**
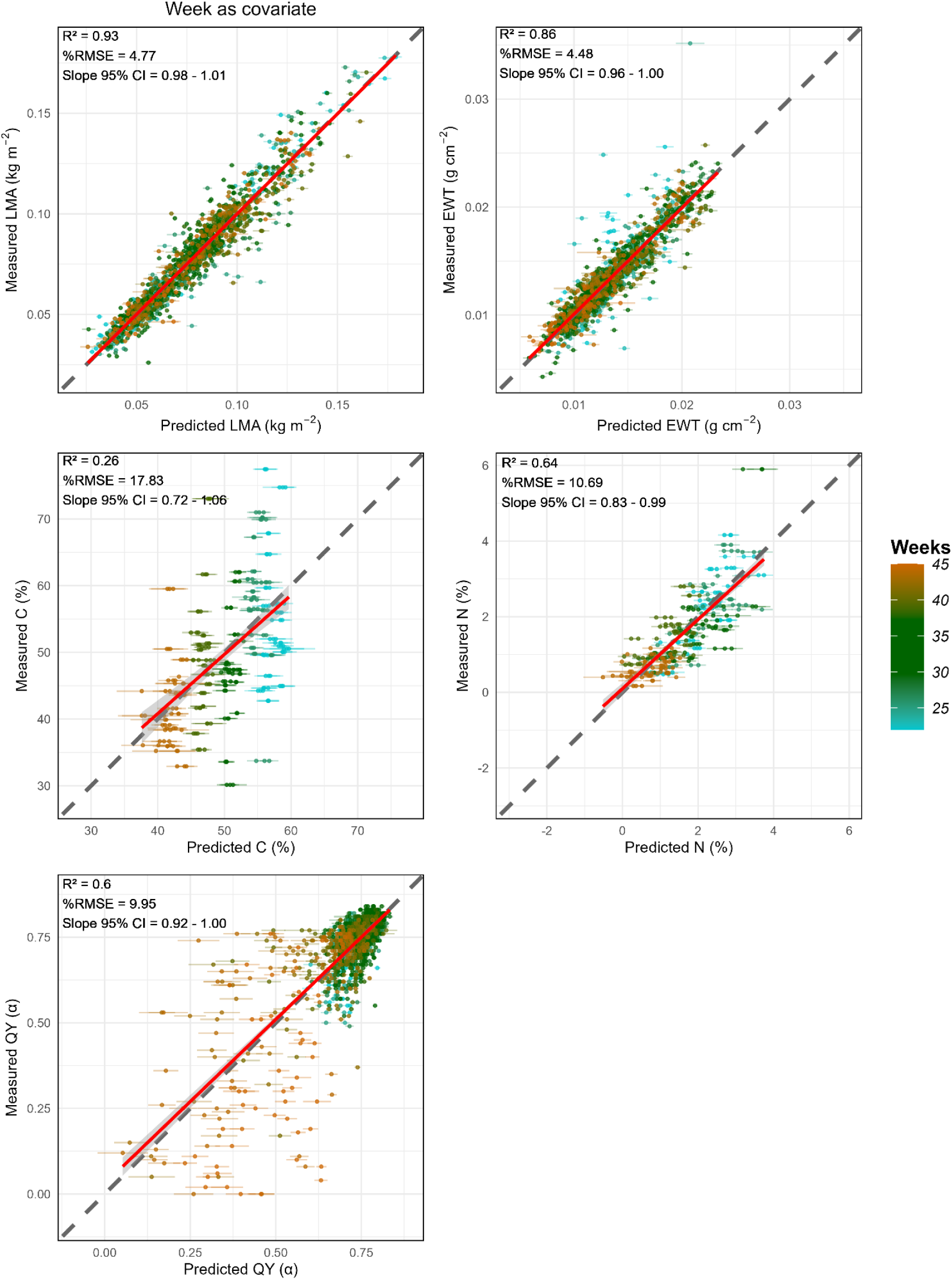
Validation results for *week-as-covariate* model coefficients assessed on measured traits by week. The dark grey dashed line in each panel is the 1:1 line. The 95% confidence interval (CI) for the slope shows how significantly different the regression line is from the 1:1 line. Weeks = Week of Year (WOY). Weekly measurement (22-45) covers from May 31st (shortly after leaf emergence) to November 6th, 2023 (senescence). Leaf mass per area (LMA); Equivalent water thickness (EWT); Carbon (C); Nitrogen (N); Quantum Yield (QY).

**Fig. S8.**
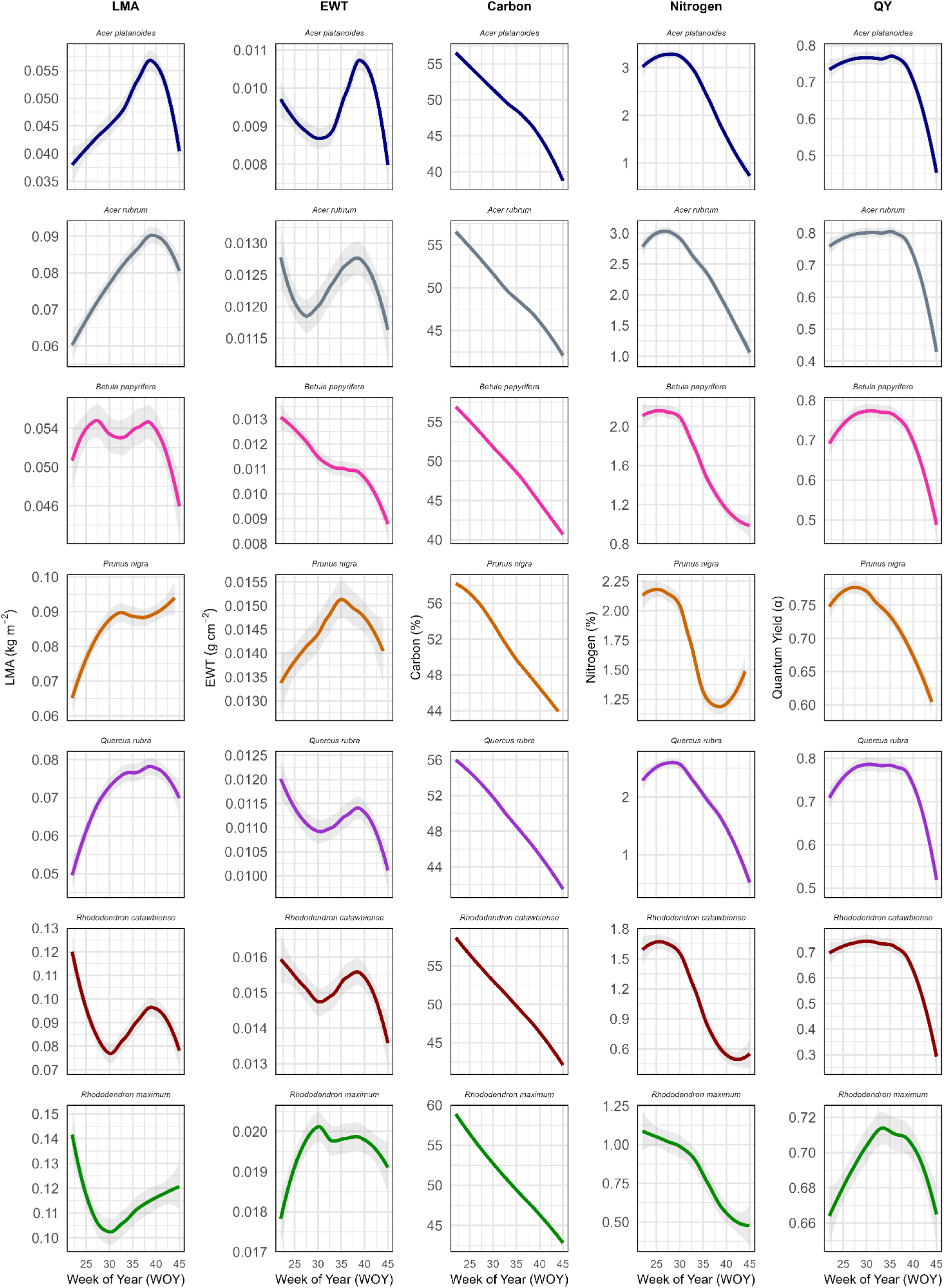
Trait temporal dynamics for LMA, EWT, Carbon, Nitrogen, and QY from *week-as-covariate* model. LMA, Leaf mass per area; EWT, Equivalent water thickness; QY, Quantum Yield. Weekly prediction (22-45) covers from May 31st (shortly after leaf emergence) to November 6th, 2023 (senescence).

**Fig. S9.**
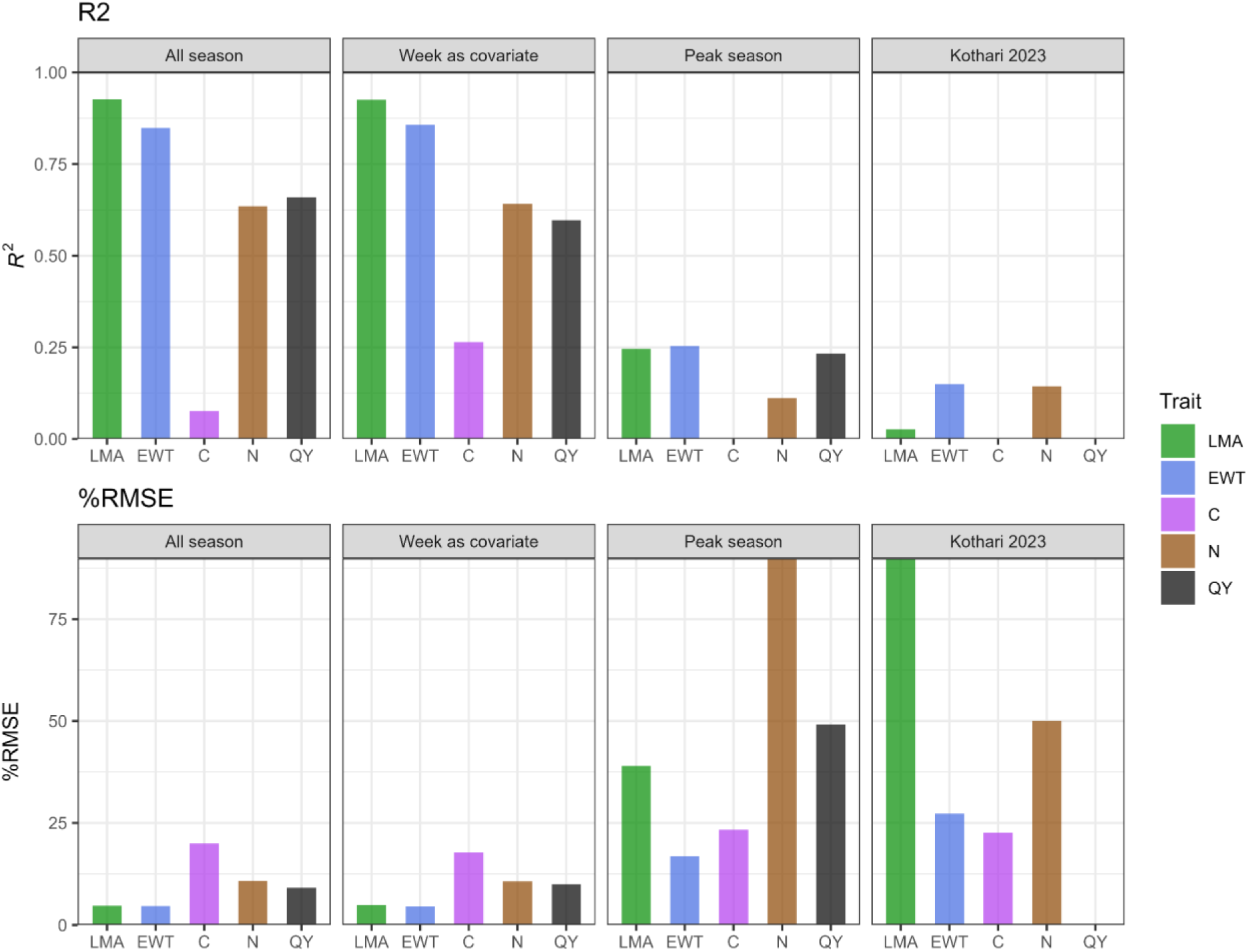
Comparison of performance metrics across models on measured datasets. Leaf mass per area (LMA); Equivalent water thickness (EWT); Carbon (C); Nitrogen (N); Quantum Yield (QY). *Kothari 2023* model coefficients for trait prediction were obtained from (Kothari *et al*., 2023).

**Fig. S10.**
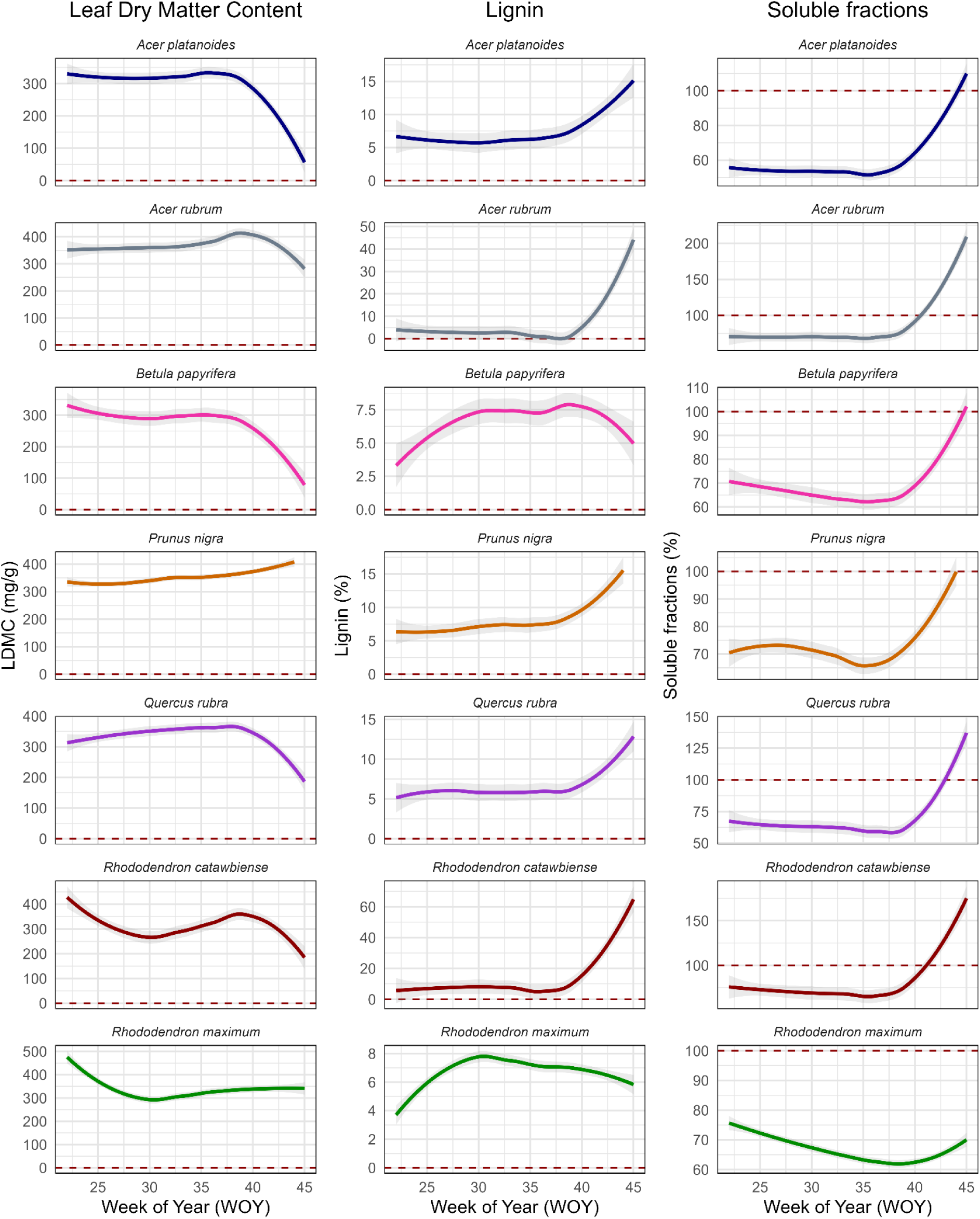
Temporal assessment of LDMC, Lignin, and Soluble fractions predictions from. (Kothari *et al*., 2023) **model coefficients.** Leaf dry matter content (LDMC). Weekly prediction (22-45) covers from May 31st (shortly after leaf emergence) to November 6th, 2023 (senescence).

**Fig. S11.**
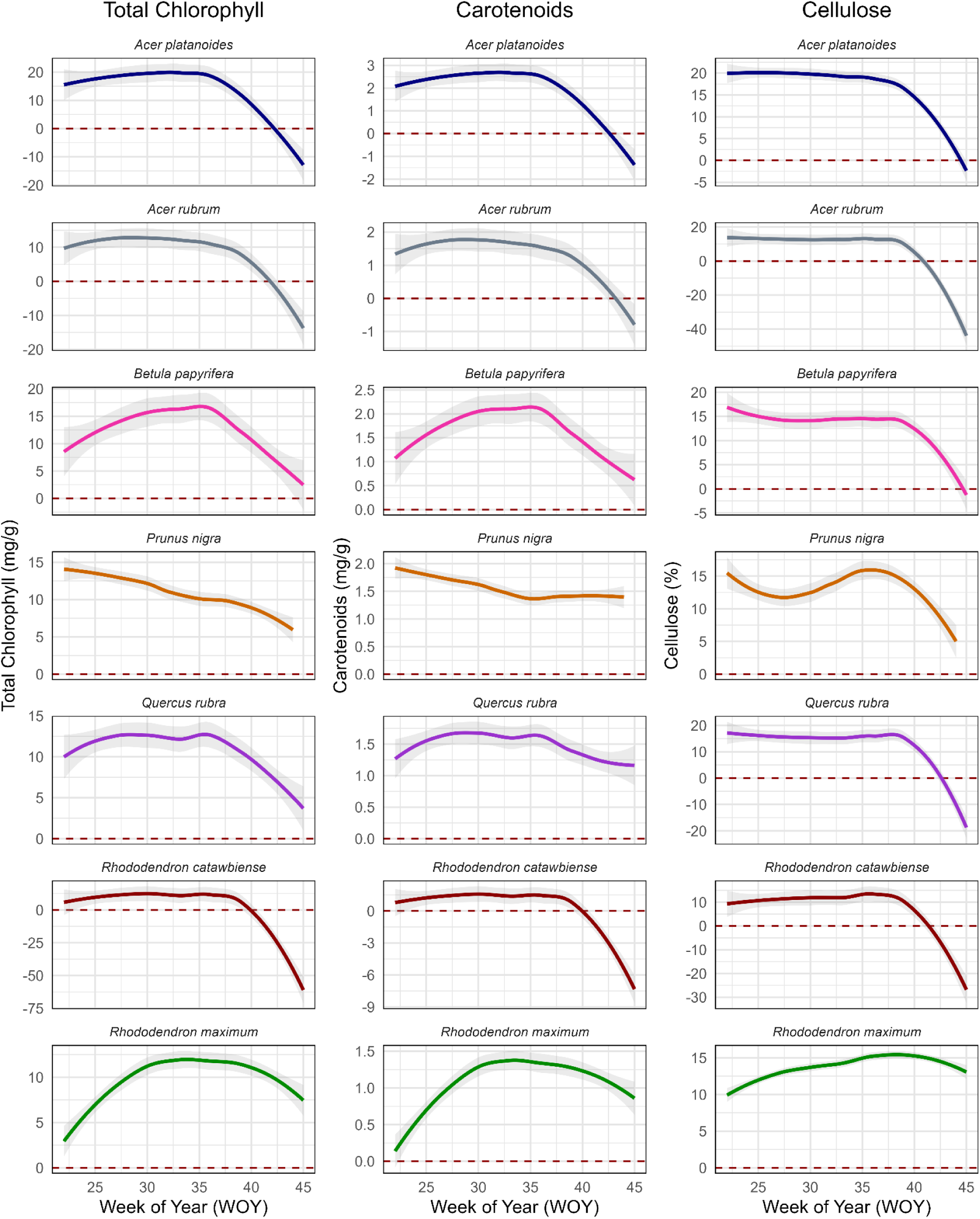
Temporal assessment of Total Chlorophyll, Carotenoids, and Cellulose prediction from. (Kothari *et al*., 2023) **model coefficients.** Weekly prediction (22-45) covers from May 31st (shortly after leaf emergence) to November 6th, 2023 (senescence).

**Fig. S12.**
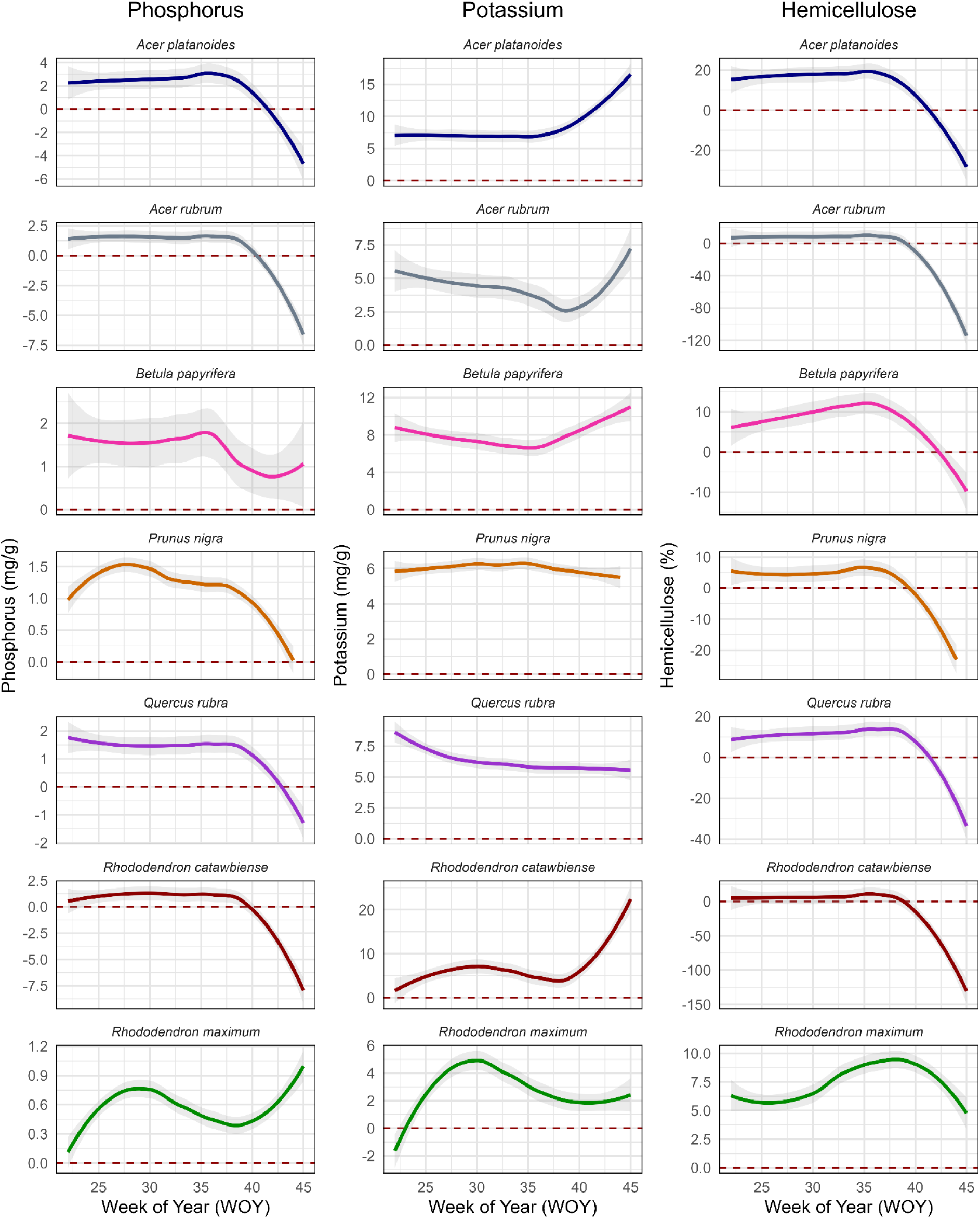
Temporal assessment of Phosphorus, Potassium, and Hemicellulose predictions from. (Kothari *et al*., 2023) **model coefficients.** Weekly prediction (22-45) covers from May 31st (shortly after leaf emergence) to November 6th, 2023 (senescence).

### Result S1. Trait prediction from model fitting

We show that PLSR can build models for predicting trait temporal dynamics from reflectance spectra, which vary in data sizes and predictor variables. The performance metrics for the three models—*all-season*, *week-as-covariate,* and *peak-season*—show higher accuracy for LMA and EWT (R^2^ = 0.81 - 0.94, %RMSE = 4.78 - 7.61), intermediate accuracy for N and QY (R^2^ = 0.54 - 0.67, %RMSE = 9.76 - 19.91), and low accuracy for C (R^2^ = 0.11 - 0.16, %RMSE = 21.21 - 23.88) (Fig. S2, S3, Table S4). The low to intermediate accuracy for C and N can be attributed to its small data size compared to LMA, EWT, and QY.

The VIP metrics across the three models show the critical regions of the spectrum that were important in predicting the traits (Fig. S4). Across the three models, we observed that (i) the VIP metrics for models are similar but not entirely the same. (ii) The VIS, NIR, and SWIR influenced LMA and EWT prediction in *all-season* and *peak-season*. Similarly, the same regions influenced the prediction of C and N, but in smaller wavelengths. Interestingly, only the VIS region influenced the prediction of QY. (iii) In *week-as-covariate*, the VIS region was important in predicting all traits. (iv) The important spectrum regions for trait prediction in the *week-as-covariate* model are smaller compared to *all-season* and *peak-season*. (v) In *week-as-covariate*, the VIP metrics for C had no important spectrum region at VIP = 1, while all other models and traits had important spectrum regions for trait prediction (Fig. S4). The lack of important spectral regions for C and smaller wavelength ranges for other traits in the *week-as-covariate* model could be attributed to the influence of ‘week’ as a predictor alongside spectra.

**Table S1.** Selected taxa and summary of sampling design for monitoring leaf phenology.

| Species | Family | Habit | *n* weeks | *n* spectra |
| --- | --- | --- | --- | --- |
| *Acer platanoides* L. | Sapindaceae | Tree - deciduous | 24 | 1080 |
| *A. rubrum* L. | Sapindaceae | Tree - deciduous | 24 | 1080 |
| *Betula papyrifera* Marshall | Myrtaceae | Tree - deciduous | 24 | 1080 |
| *Prunus nigra* Aiton | Rosaceae | Tree - deciduous | 24 | 1080 |
| *Quercus rubra* L. | Fagaceae | Tree - deciduous | 23 | 1035 |
| *Rhododendron catawbiense* Michx. | Ericaceae | Shrub - evergreen | 24 | 1080 |
| *R. maximum* L. | Ericaceae | Shrub - evergreen | 24 | 1080 |

**Table S2.**
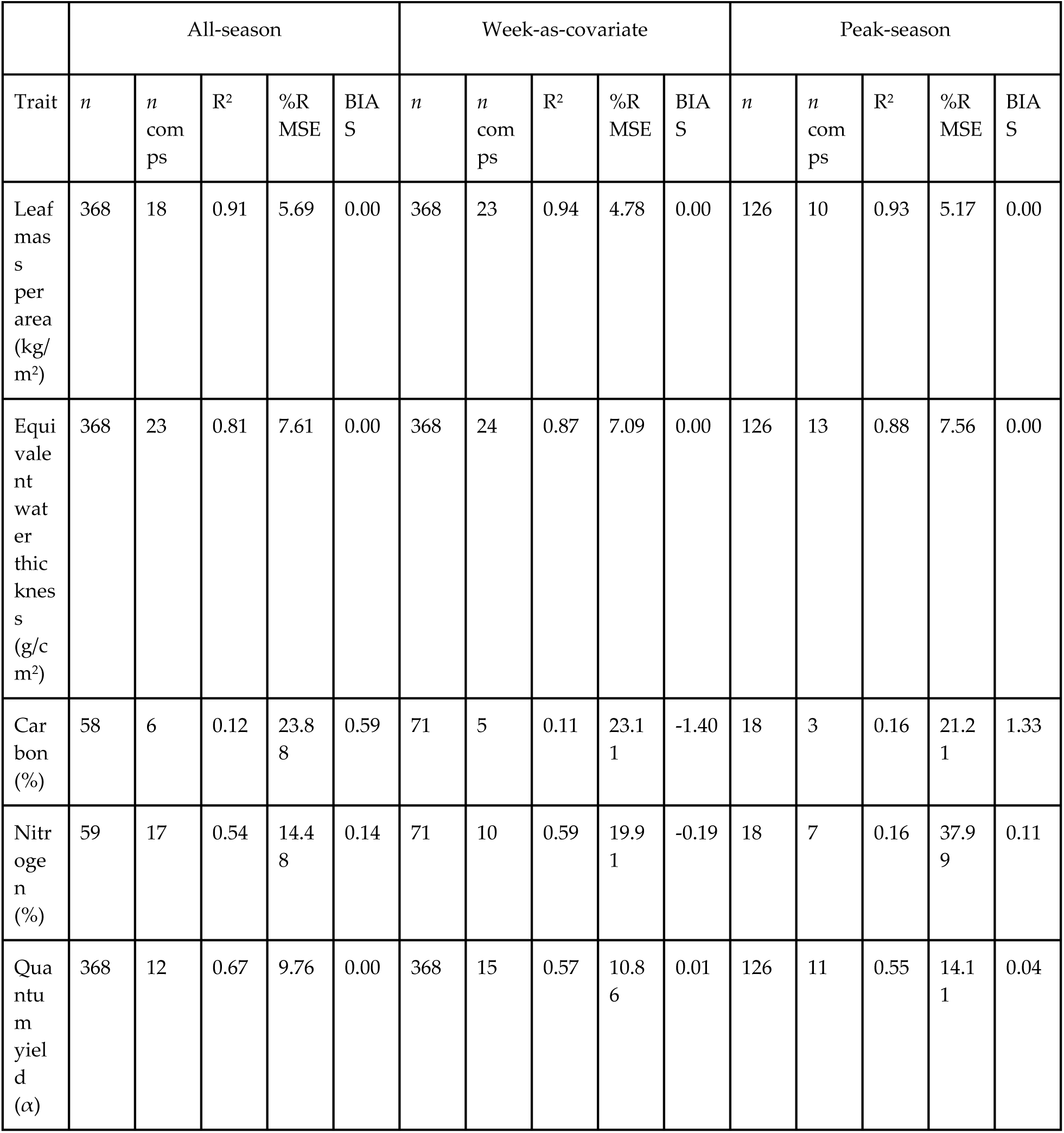
Model fitting summary statistics for fresh leaf spectra trait models averaged over 1000 iterations.

**Table S3.** Significant differences across leaf phenological stages for measured traits.

| Trait | Species | Term | Df | Sum.Sq | Mean.Sq | F.value | Pr..F. |
| --- | --- | --- | --- | --- | --- | --- | --- |
| LMA | Quercus rubra | Week | 2 | 0.006893 | 0.003447 | 35.41695 | 6.90E-08 |
| LMA | Quercus rubra | Residuals | 24 | 0.002336 | 9.73E-05 | NA | NA |
| LMA | Betula papyrifera | Week | 2 | 0.000264 | 0.000132 | 5.817546 | 0.00871 |
| LMA | Betula papyrifera | Residuals | 24 | 0.000545 | 2.27E-05 | NA | NA |
| LMA | Prunus nigra | Week | 2 | 0.006085 | 0.003042 | 40.63348 | 4.10E-08 |
| LMA | Prunus nigra | Residuals | 22 | 0.001647 | 7.49E-05 | NA | NA |
| LMA | Acer rubrum | Week | 2 | 0.003446 | 0.001723 | 16.74004 | 2.81E-05 |
| LMA | Acer rubrum | Residuals | 24 | 0.00247 | 0.000103 | NA | NA |
| LMA | Rhododendron maximum | Week | 2 | 0.008833 | 0.004416 | 12.72962 | 0.00017 |
| LMA | Rhododendron maximum | Residuals | 24 | 0.008326 | 0.000347 | NA | NA |
| LMA | Rhododendron catawbiense | Week | 2 | 0.008171 | 0.004085 | 26.28456 | 8.99E-07 |
| LMA | Rhododendron catawbiense | Residuals | 24 | 0.00373 | 0.000155 | NA | NA |
| LMA | Acer platanoides | Week | 2 | 0.000263 | 0.000132 | 4.059908 | 0.030287 |
| LMA | Acer platanoides | Residuals | 24 | 0.000779 | 3.24E-05 | NA | NA |
| EWT | Quercus rubra | Week | 2 | 2.54E-06 | 1.27E-06 | 0.453285 | 0.640867 |
| EWT | Quercus rubra | Residuals | 24 | 6.71E-05 | 2.80E-06 | NA | NA |
| EWT | Betula papyrifera | Week | 2 | 7.74E-05 | 3.87E-05 | 44.20522 | 8.97E-09 |
| EWT | Betula papyrifera | Residuals | 24 | 2.10E-05 | 8.75E-07 | NA | NA |
| EWT | Prunus nigra | Week | 2 | 3.09E-05 | 1.54E-05 | 6.327771 | 0.006748 |
| EWT | Prunus nigra | Residuals | 22 | 5.37E-05 | 2.44E-06 | NA | NA |
| EWT | Acer rubrum | Week | 2 | 0.000182 | 9.09E-05 | 41.18267 | 1.74E-08 |
| EWT | Acer rubrum | Residuals | 24 | 5.30E-05 | 2.21E-06 | NA | NA |
| EWT | Rhododendron maximum | Week | 2 | 1.49E-05 | 7.44E-06 | 1.419768 | 0.261357 |
| EWT | Rhododendron maximum | Residuals | 24 | 0.000126 | 5.24E-06 | NA | NA |
| EWT | Rhododendron catawbiense | Week | 2 | 3.03E-05 | 1.52E-05 | 5.025839 | 0.015027 |
| EWT | Rhododendron catawbiense | Residuals | 24 | 7.24E-05 | 3.01E-06 | NA | NA |
| EWT | Acer platanoides | Week | 2 | 1.02E-05 | 5.08E-06 | 3.964254 | 0.032538 |
| EWT | Acer platanoides | Residuals | 24 | 3.07E-05 | 1.28E-06 | NA | NA |
| N | Quercus rubra | Week | 2 | 13.88207 | 6.941033 | 155.1647 | 1.87E-14 |
| N | Quercus rubra | Residuals | 24 | 1.0736 | 0.044733 | NA | NA |
| N | Betula papyrifera | Week | 2 | 21.2906 | 10.6453 | 50.63364 | 2.45E-09 |
| N | Betula papyrifera | Residuals | 24 | 5.0458 | 0.210242 | NA | NA |
| N | Prunus nigra | Week | 2 | 8.996093 | 4.498046 | 81.77107 | 6.51E-11 |
| N | Prunus nigra | Residuals | 22 | 1.210171 | 0.055008 | NA | NA |
| N | Acer rubrum | Week | 2 | 25.1282 | 12.5641 | 33.51618 | 1.13E-07 |
| N | Acer rubrum | Residuals | 24 | 8.9968 | 0.374867 | NA | NA |
| N | Rhododendron maximum | Week | 2 | 0.018467 | 0.009233 | 0.097889 | 0.90711 |
| N | Rhododendron maximum | Residuals | 24 | 2.2638 | 0.094325 | NA | NA |
| N | Rhododendron catawbiense | Week | 2 | 3.2342 | 1.6171 | 87.56859 | 9.39E-12 |
| N | Rhododendron catawbiense | Residuals | 24 | 0.4432 | 0.018467 | NA | NA |
| N | Acer platanoides | Week | 2 | 43.68327 | 21.84163 | 16.09968 | 3.68E-05 |
| N | Acer platanoides | Residuals | 24 | 32.5596 | 1.35665 | NA | NA |
| C | Quercus rubra | Week | 2 | 78.80207 | 39.40103 | 3.900912 | 0.034128 |
| C | Quercus rubra | Residuals | 24 | 242.4112 | 10.10047 | NA | NA |
| C | Betula papyrifera | Week | 2 | 1637.413 | 818.7066 | 13.84569 | 0.0001 |
| C | Betula papyrifera | Residuals | 24 | 1419.139 | 59.13077 | NA | NA |
| C | Prunus nigra | Week | 2 | 367.4119 | 183.7059 | 7.526417 | 0.003233 |
| C | Prunus nigra | Residuals | 22 | 536.9794 | 24.40815 | NA | NA |
| C | Acer rubrum | Week | 2 | 1912.017 | 956.0084 | 7.47579 | 0.002994 |
| C | Acer rubrum | Residuals | 24 | 3069.134 | 127.8806 | NA | NA |
| C | Rhododendron maximum | Week | 2 | 1402.604 | 701.3022 | 9.720621 | 0.000809 |
| C | Rhododendron maximum | Residuals | 24 | 1731.5 | 72.14583 | NA | NA |
| C | Rhododendron catawbiense | Week | 2 | 791.1261 | 395.563 | 12.56037 | 0.000185 |
| C | Rhododendron catawbiense | Residuals | 24 | 755.8306 | 31.49294 | NA | NA |
| C | Acer platanoides | Week | 2 | 905.6474 | 452.8237 | 23.37687 | 4.56E-06 |
| C | Acer platanoides | Residuals | 21 | 406.7823 | 19.37059 | NA | NA |
| QY | Quercus rubra | Week | 2 | 0.22343 | 0.111715 | 71.26521 | 8.03E-11 |
| QY | Quercus rubra | Residuals | 24 | 0.037622 | 0.001568 | NA | NA |
| QY | Betula papyrifera | Week | 2 | 0.915556 | 0.457778 | 13.01875 | 0.000148 |
| QY | Betula papyrifera | Residuals | 24 | 0.843911 | 0.035163 | NA | NA |
| QY | Prunus nigra | Week | 2 | 0.237425 | 0.118712 | 14.15456 | 0.000112 |
| QY | Prunus nigra | Residuals | 22 | 0.184511 | 0.008387 | NA | NA |
| QY | Acer rubrum | Week | 2 | 0.69983 | 0.349915 | 17.7555 | 1.85E-05 |
| QY | Acer rubrum | Residuals | 24 | 0.472978 | 0.019707 | NA | NA |
| QY | Rhododendron maximum | Week | 2 | 0.065163 | 0.032581 | 15.83618 | 4.12E-05 |
| QY | Rhododendron maximum | Residuals | 24 | 0.049378 | 0.002057 | NA | NA |
| QY | Rhododendron catawbiense | Week | 2 | 1.702067 | 0.851033 | 109.4968 | 8.62E-13 |
| QY | Rhododendron catawbiense | Residuals | 24 | 0.186533 | 0.007772 | NA | NA |
| QY | Acer platanoides | Week | 2 | 0.360689 | 0.180344 | 13.28867 | 0.00013 |
| QY | Acer platanoides | Residuals | 24 | 0.325711 | 0.013571 | NA | NA |

**Table S4.**
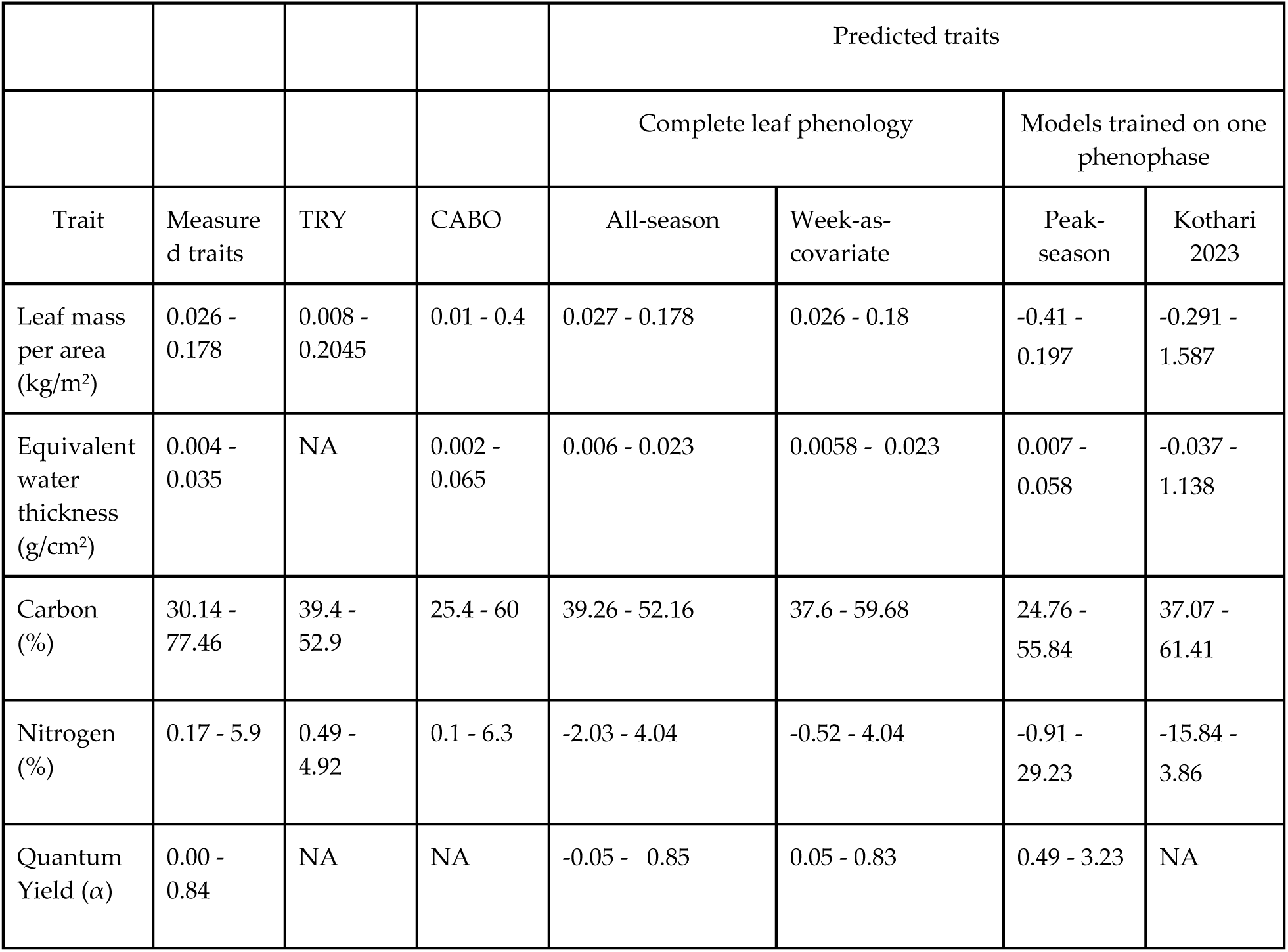
Comparison of trait range across datasets—measured, model predictions, TRY, and CABO database. Equivalent water thickness was only obtained from the Canadian Airborne Biodiversity Observatory (CABO) (Fig. 1, (Kothari *et al*., 2023), as TRY had fewer data points (Kattge *et al*., 2020).

**Table S5.** Summary statistics for *week-as-covariate* model performance over time on measured traits.

| Complete leaf phenology model |  |  |  |  |
| --- | --- | --- | --- | --- |
| Week-as-covariate |  |  |  |  |
| Traits | *n* | R<sup>2</sup> | %RMSE | BIAS |
| Leaf mass per area (kg/m<sup>2</sup>) | 1500 | 0.93 | 4.77 | 0.00 |
| Equivalent water thickness (g/cm<sup>2</sup>) | 1500 | 0.86 | 4.48 | 0.00 |
| Carbon (%) | 304 | 0.26 | 17.83 | -0.35 |
| Nitrogen (%) | 310 | 0.64 | 10.69 | -0.05 |
| Quantum yield ($\alpha$) | 1500 | 0.6 | 9.95 | 0.08 |

**Table S6.** Significant differences across leaf phenophases between measured and predicted traits from complete leaf phenology trait models.

| Trait | Species | Phenophase | Comparison | p adj | Significance | ANOVA_F | ANOVA_p | ANOVA_sig |
| --- | --- | --- | --- | --- | --- | --- | --- | --- |
| LMA | Quercus rubra | Young | Measured-All_season | 0.142639 | ns | 2.397352 | 0.112404 | ns |
| LMA | Quercus rubra | Young | Week_covariate-All_season | 0.988226 | ns | 2.397352 | 0.112404 | ns |
| LMA | Quercus rubra | Young | Week_covariate-Measured | 0.184929 | ns | 2.397352 | 0.112404 | ns |
| LMA | Quercus rubra | Mature | Measured-All_season | 0.946984 | ns | 0.059495 | 0.942379 | ns |
| LMA | Quercus rubra | Mature | Week_covariate-All_season | 0.999322 | ns | 0.059495 | 0.942379 | ns |
| LMA | Quercus rubra | Mature | Week_covariate-Measured | 0.957881 | ns | 0.059495 | 0.942379 | ns |
| LMA | Quercus rubra | Old/Senescent | Measured-All_season | 0.777527 | ns | 0.379773 | 0.688054 | ns |
| L<br>M<br>A | Quercus rubra | Old/Senescent | Week_covariate-All_season | 0.9903<br>17 | ns | 0.3797<br>73 | 0.6880<br>54 | ns |
| L<br>M<br>A | Quercus rubra | Old/Senescent | Week_covariate-Measured | 0.6991<br>23 | ns | 0.3797<br>73 | 0.6880<br>54 | ns |
| L<br>M<br>A | Betula papyrifera | Young | Measured-All_season | 0.6006<br>88 | ns | 0.7717<br>34 | 0.4733<br>44 | ns |
| L<br>M<br>A | Betula papyrifera | Young | Week_covariate-All_season | 0.9817<br>53 | ns | 0.7717<br>34 | 0.4733<br>44 | ns |
| L<br>M<br>A | Betula papyrifera | Young | Week_covariate-Measured | 0.4903<br>62 | ns | 0.7717<br>34 | 0.4733<br>44 | ns |
| L<br>M<br>A | Betula papyrifera | Mature | Measured-All_season | 0.8207<br>37 | ns | 0.2175<br>1 | 0.8060<br>88 | ns |
| L<br>M<br>A | Betula papyrifera | Mature | Week_covariate-All_season | 0.9976<br>04 | ns | 0.2175<br>1 | 0.8060<br>88 | ns |
| L<br>M<br>A | Betula papyrifera | Mature | Week_covariate-Measured | 0.8548<br>08 | ns | 0.2175<br>1 | 0.8060<br>88 | ns |
| L<br>M<br>A | Betula papyrifera | Old/Senescent | Measured-All_season | 0.8410<br>2 | ns | 0.1582<br>84 | 0.8544<br>91 | ns |
| L<br>M<br>A | Betula papyrifera | Old/Senescent | Week_covariate-All_season | 0.9587<br>06 | ns | 0.1582<br>84 | 0.8544<br>91 | ns |
| L<br>M<br>A | Betula papyrifera | Old/Senescent | Week_covariate-Measured | 0.9560<br>58 | ns | 0.1582<br>84 | 0.8544<br>91 | ns |
| L<br>M<br>A | Prunus nigra | Young | Measured-All_season | 0.3449<br>63 | ns | 1.0808<br>17 | 0.3552<br>68 | ns |
| L<br>M<br>A | Prunus nigra | Young | Week_covariate-All_season | 0.9182<br>23 | ns | 1.0808<br>17 | 0.3552<br>68 | ns |
| L<br>M<br>A | Prunus nigra | Young | Week_covariate-Measured | 0.5658<br>72 | ns | 1.0808<br>17 | 0.3552<br>68 | ns |
| L<br>M<br>A | Prunus nigra | Mature | Measured-All_season | 0.8747<br>7 | ns | 0.1361<br>15 | 0.8734<br>11 | ns |
| L<br>M<br>A | Prunus nigra | Mature | Week_covariate-All_season | 0.9942<br>83 | ns | 0.1361<br>15 | 0.8734<br>11 | ns |
| L<br>M<br>A | Prunus nigra | Mature | Week_covariate-Measured | 0.9190<br>14 | ns | 0.1361<br>15 | 0.8734<br>11 | ns |
| L<br>M<br>A | Prunus nigra | Old/Senes-<br>cent | Measured-All_season | 0.0484<br>74 | * | 4.5637<br>36 | 0.0249<br>33 | * |
| L<br>M<br>A | Prunus nigra | Old/Senes-<br>cent | Week_covariate-All_season | 0.9950<br>69 | ns | 4.5637<br>36 | 0.0249<br>33 | * |
| L<br>M<br>A | Prunus nigra | Old/Senes-<br>cent | Week_covariate-Measured | 0.0400<br>98 | * | 4.5637<br>36 | 0.0249<br>33 | * |
| L<br>M<br>A | Acer rubrum | Young | Measured-All_season | 0.3910<br>42 | ns | 1.3429<br>32 | 0.2800<br>01 | ns |
| L<br>M<br>A | Acer rubrum | Young | Week_covariate-All_season | 0.9862<br>59 | ns | 1.3429<br>32 | 0.2800<br>01 | ns |
| L<br>M<br>A | Acer rubrum | Young | Week_covariate-Measured | 0.3124<br>44 | ns | 1.3429<br>32 | 0.2800<br>01 | ns |
| L<br>M<br>A | Acer rubrum | Mature | Measured-All_season | 0.9984<br>13 | ns | 0.1049<br>22 | 0.9008<br>06 | ns |
| L<br>M<br>A | Acer rubrum | Mature | Week_covariate-All_season | 0.9286<br>12 | ns | 0.1049<br>22 | 0.9008<br>06 | ns |
| L<br>M<br>A | Acer rubrum | Mature | Week_covariate-Measured | 0.9073<br>73 | ns | 0.1049<br>22 | 0.9008<br>06 | ns |
| L<br>M<br>A | Acer rubrum | Old/Senes-<br>cent | Measured-All_season | 0.8961<br>58 | ns | 0.1946<br>51 | 0.8244<br>08 | ns |
| L<br>M<br>A | Acer rubrum | Old/Senes-<br>cent | Week_covariate-All_season | 0.9871<br>17 | ns | 0.1946<br>51 | 0.8244<br>08 | ns |
| L<br>M<br>A | Acer rubrum | Old/Senes-<br>cent | Week_covariate-Measured | 0.8211<br>91 | ns | 0.1946<br>51 | 0.8244<br>08 | ns |
| L<br>M<br>A | Rhododendron<br>maximum | Young | Measured-All_season | 0.9148<br>09 | ns | 0.1157<br>8 | 0.8911<br>66 | ns |
| L<br>M<br>A | Rhododendron<br>maximum | Young | Week_covariate-All_season | 0.9996<br>04 | ns | 0.1157<br>8 | 0.8911<br>66 | ns |
| L<br>M<br>A | Rhododendron<br>maximum | Young | Week_covariate-Measured | 0.9037<br>27 | ns | 0.1157<br>8 | 0.8911<br>66 | ns |
| L<br>M<br>A | Rhododendron<br>maximum | Mature | Measured-All_season | 0.8467<br>95 | ns | 0.1908<br>25 | 0.8275<br>18 | ns |
| L<br>M<br>A | Rhododendron<br>maximum | Mature | Week_covariate-All_season | 0.9993<br>54 | ns | 0.1908<br>25 | 0.8275<br>18 | ns |
| L<br>M<br>A | Rhododendron<br>maximum | Mature | Week_covariate-Measured | 0.8637<br>93 | ns | 0.1908<br>25 | 0.8275<br>18 | ns |
| L<br>M<br>A | Rhododendron<br>maximum | Old/Senes<br>cent | Measured-All_season | 0.8774<br>64 | ns | 0.1754<br>72 | 0.8401<br>27 | ns |
| L<br>M<br>A | Rhododendron<br>maximum | Old/Senes<br>cent | Week_covariate-All_season | 0.9988<br>25 | ns | 0.1754<br>72 | 0.8401<br>27 | ns |
| L<br>M<br>A | Rhododendron<br>maximum | Old/Senes<br>cent | Week_covariate-Measured | 0.8551<br>97 | ns | 0.1754<br>72 | 0.8401<br>27 | ns |
| L<br>M<br>A | Rhododendron<br>catawbiense | Young | Measured-All_season | 0.9138<br>22 | ns | 0.1025<br>73 | 0.9029<br>06 | ns |
| L<br>M<br>A | Rhododendron<br>catawbiense | Young | Week_covariate-All_season | 0.9995<br>93 | ns | 0.1025<br>73 | 0.9029<br>06 | ns |
| L<br>M<br>A | Rhododendron<br>catawbiense | Young | Week_covariate-Measured | 0.9245<br>29 | ns | 0.1025<br>73 | 0.9029<br>06 | ns |
| L<br>M<br>A | Rhododendron<br>catawbiense | Mature | Measured-All_season | 0.5191<br>51 | ns | 0.8448<br>42 | 0.4420<br>08 | ns |
| L<br>M<br>A | Rhododendron<br>catawbiense | Mature | Week_covariate-All_season | 0.9992<br>59 | ns | 0.8448<br>42 | 0.4420<br>08 | ns |
| L<br>M<br>A | Rhododendron<br>catawbiense | Mature | Week_covariate-Measured | 0.4974<br>13 | ns | 0.8448<br>42 | 0.4420<br>08 | ns |
| L<br>M<br>A | Rhododendron<br>catawbiense | Old/Senes<br>cent | Measured-All_season | 0.9911<br>61 | ns | 0.0092<br>19 | 0.9908<br>27 | ns |
| L<br>M<br>A | Rhododendron<br>catawbiense | Old/Senes<br>cent | Week_covariate-All_season | 0.9997<br>41 | ns | 0.0092<br>19 | 0.9908<br>27 | ns |
| L<br>M<br>A | Rhododendron<br>catawbiense | Old/Senes<br>cent | Week_covariate-Measured | 0.9939<br>13 | ns | 0.0092<br>19 | 0.9908<br>27 | ns |
| L<br>M<br>A | Acer platanoides | Young | Measured-All_season | 0.3325<br>22 | ns | 1.6184<br>57 | 0.2190<br>99 | ns |
| L<br>M<br>A | Acer platanoides | Young | Week_covariate-All_season | 0.9785<br>24 | ns | 1.6184<br>57 | 0.2190<br>99 | ns |
| L<br>M<br>A | Acer platanoides | Young | Week_covariate-Measured | 0.2456<br>84 | ns | 1.6184<br>57 | 0.2190<br>99 | ns |
| L<br>M<br>A | Acer platanoides | Mature | Measured-All_season | 0.3939<br>36 | ns | 1.2108<br>59 | 0.3155<br>02 | ns |
| L<br>M<br>A | Acer platanoides | Mature | Week_covariate-All_season | 0.9991<br>75 | ns | 1.2108<br>59 | 0.3155<br>02 | ns |
| L<br>M<br>A | Acer platanoides | Mature | Week_covariate-Measured | 0.3737<br>2 | ns | 1.2108<br>59 | 0.3155<br>02 | ns |
| L<br>M<br>A | Acer platanoides | Old/Senes<br>cent | Measured-All_season | 0.9470<br>46 | ns | 0.0835<br>77 | 0.9200<br>87 | ns |
| L<br>M<br>A | Acer platanoides | Old/Senes<br>cent | Week_covariate-All_season | 0.9973<br>84 | ns | 0.0835<br>77 | 0.9200<br>87 | ns |
| L<br>M<br>A | Acer platanoides | Old/Senescent | Week_covariate-Measured | 0.922403 | ns | 0.083577 | 0.920087 | ns |
| E<br>W<br>T | Quercus rubra | Young | Measured-All_season | 0.046668 | * | 3.781599 | 0.037356 | * |
| E<br>W<br>T | Quercus rubra | Young | Week_covariate-All_season | 0.941512 | ns | 3.781599 | 0.037356 | * |
| E<br>W<br>T | Quercus rubra | Young | Week_covariate-Measured | 0.091768 | ns | 3.781599 | 0.037356 | * |
| E<br>W<br>T | Quercus rubra | Mature | Measured-All_season | 0.943493 | ns | 0.054693 | 0.946894 | ns |
| E<br>W<br>T | Quercus rubra | Mature | Week_covariate-All_season | 0.974959 | ns | 0.054693 | 0.946894 | ns |
| E<br>W<br>T | Quercus rubra | Mature | Week_covariate-Measured | 0.993275 | ns | 0.054693 | 0.946894 | ns |
| E<br>W<br>T | Quercus rubra | Old/Senescent | Measured-All_season | 0.86859 | ns | 0.15297 | 0.858985 | ns |
| E<br>W<br>T | Quercus rubra | Old/Senescent | Week_covariate-All_season | 0.997859 | ns | 0.15297 | 0.858985 | ns |
| E<br>W<br>T | Quercus rubra | Old/Senescent | Week_covariate-Measured | 0.897152 | ns | 0.15297 | 0.858985 | ns |
| E<br>W<br>T | Betula papyrifera | Young | Measured-All_season | 0.999985 | ns | 0.110018 | 0.896267 | ns |
| E<br>W<br>T | Betula papyrifera | Young | Week_covariate-All_season | 0.9144<br>6 | ns | 0.1100<br>18 | 0.8962<br>67 | ns |
| E<br>W<br>T | Betula papyrifera | Young | Week_covariate-Measured | 0.9123<br>09 | ns | 0.1100<br>18 | 0.8962<br>67 | ns |
| E<br>W<br>T | Betula papyrifera | Mature | Measured-All_season | 0.9985<br>36 | ns | 0.0013<br>29 | 0.9986<br>72 | ns |
| E<br>W<br>T | Betula papyrifera | Mature | Week_covariate-All_season | 0.9996<br>24 | ns | 0.0013<br>29 | 0.9986<br>72 | ns |
| E<br>W<br>T | Betula papyrifera | Mature | Week_covariate-Measured | 0.9996<br>43 | ns | 0.0013<br>29 | 0.9986<br>72 | ns |
| E<br>W<br>T | Betula papyrifera | Old/Senes-<br>cent | Measured-All_season | 0.5690<br>53 | ns | 0.5654<br>86 | 0.5754<br>71 | ns |
| E<br>W<br>T | Betula papyrifera | Old/Senes-<br>cent | Week_covariate-All_season | 0.9621<br>49 | ns | 0.5654<br>86 | 0.5754<br>71 | ns |
| E<br>W<br>T | Betula papyrifera | Old/Senes-<br>cent | Week_covariate-Measured | 0.7308<br>31 | ns | 0.5654<br>86 | 0.5754<br>71 | ns |
| E<br>W<br>T | Prunus nigra | Young | Measured-All_season | 0.4954<br>34 | ns | 0.9746<br>02 | 0.3917<br>84 | ns |
| E<br>W<br>T | Prunus nigra | Young | Week_covariate-All_season | 0.9926<br>26 | ns | 0.9746<br>02 | 0.3917<br>84 | ns |
| E<br>W<br>T | Prunus nigra | Young | Week_covariate-Measured | 0.4292<br>04 | ns | 0.9746<br>02 | 0.3917<br>84 | ns |
| E<br>W<br>T | Prunus nigra | Mature | Measured-All_season | 0.9385<br>74 | ns | 0.0961<br>76 | 0.9086<br>52 | ns |
| E<br>W<br>T | Prunus nigra | Mature | Week_covariate-All_season | 0.9117<br>78 | ns | 0.0961<br>76 | 0.9086<br>52 | ns |
| E<br>W<br>T | Prunus nigra | Mature | Week_covariate-Measured | 0.9972<br>58 | ns | 0.0961<br>76 | 0.9086<br>52 | ns |
| E<br>W<br>T | Prunus nigra | Old/Senes-<br>cent | Measured-All_season | 0.7266<br>67 | ns | 0.3172<br>62 | 0.7321<br>29 | ns |
| E<br>W<br>T | Prunus nigra | Old/Senes-<br>cent | Week_covariate-All_season | 0.9777<br>77 | ns | 0.3172<br>62 | 0.7321<br>29 | ns |
| E<br>W<br>T | Prunus nigra | Old/Senes-<br>cent | Week_covariate-Measured | 0.8394<br>97 | ns | 0.3172<br>62 | 0.7321<br>29 | ns |
| E<br>W<br>T | Acer rubrum | Young | Measured-All_season | 2.69E-09 | *** | 57.701<br>26 | 6.78E-10 | *** |
| E<br>W<br>T | Acer rubrum | Young | Week_covariate-All_season | 0.6941<br>98 | ns | 57.701<br>26 | 6.78E-10 | *** |
| E<br>W<br>T | Acer rubrum | Young | Week_covariate-Measured | 1.44E-08 | *** | 57.701<br>26 | 6.78E-10 | *** |
| E<br>W<br>T | Acer rubrum | Mature | Measured-All_season | 0.9789<br>13 | ns | 0.0233<br>46 | 0.9769<br>46 | ns |
| E<br>W<br>T | Acer rubrum | Mature | Week_covariate-All_season | 0.9831<br>17 | ns | 0.0233<br>46 | 0.9769<br>46 | ns |
| E<br>W<br>T | Acer rubrum | Mature | Week_covariate-Measured | 0.9997<br>59 | ns | 0.0233<br>46 | 0.9769<br>46 | ns |
| E<br>W<br>T | Acer rubrum | Old/Senes<br>cent | Measured-All_season | 0.7868<br>04 | ns | 0.2276<br>88 | 0.7980<br>73 | ns |
| E<br>W<br>T | Acer rubrum | Old/Senes<br>cent | Week_covariate-All_season | 0.9729<br>51 | ns | 0.2276<br>88 | 0.7980<br>73 | ns |
| E<br>W<br>T | Acer rubrum | Old/Senes<br>cent | Week_covariate-Measured | 0.8992<br>51 | ns | 0.2276<br>88 | 0.7980<br>73 | ns |
| E<br>W<br>T | Rhododendron<br>maximum | Young | Measured-All_season | 0.6720<br>72 | ns | 0.3718<br>11 | 0.6933<br>87 | ns |
| E<br>W<br>T | Rhododendron<br>maximum | Young | Week_covariate-All_season | 0.9371<br>71 | ns | 0.3718<br>11 | 0.6933<br>87 | ns |
| E<br>W<br>T | Rhododendron<br>maximum | Young | Week_covariate-Measured | 0.8656<br>83 | ns | 0.3718<br>11 | 0.6933<br>87 | ns |
| E<br>W<br>T | Rhododendron<br>maximum | Mature | Measured-All_season | 0.9925<br>44 | ns | 0.0070<br>64 | 0.9929<br>63 | ns |
| E<br>W<br>T | Rhododendron<br>maximum | Mature | Week_covariate-All_season | 0.9966<br>02 | ns | 0.0070<br>64 | 0.9929<br>63 | ns |
| E<br>W<br>T | Rhododendron<br>maximum | Mature | Week_covariate-Measured | 0.9992<br>07 | ns | 0.0070<br>64 | 0.9929<br>63 | ns |
| E<br>W<br>T | Rhododendron<br>maximum | Old/Senes<br>cent | Measured-All_season | 0.0326<br>52 | * | 5.5536<br>95 | 0.0104<br>17 | * |
| E<br>W<br>T | Rhododendron<br>maximum | Old/Senes<br>cent | Week_covariate-All_season | 0.9349<br>76 | ns | 5.5536<br>95 | 0.0104<br>17 | * |
| E<br>W<br>T | Rhododendron<br>maximum | Old/Senes<br>cent | Week_covariate-Measured | 0.0148<br>4 | * | 5.5536<br>95 | 0.0104<br>17 | * |
| E<br>W<br>T | Rhododendron<br>catawbiense | Young | Measured-All_season | 0.1376<br>65 | ns | 2.5027<br>19 | 0.1029<br>87 | ns |
| E<br>W<br>T | Rhododendron<br>catawbiense | Young | Week_covariate-All_season | 0.9946<br>88 | ns | 2.5027<br>19 | 0.1029<br>87 | ns |
| E<br>W<br>T | Rhododendron<br>catawbiense | Young | Week_covariate-Measured | 0.1643<br>03 | ns | 2.5027<br>19 | 0.1029<br>87 | ns |
| E<br>W<br>T | Rhododendron<br>catawbiense | Mature | Measured-All_season | 0.9209<br>79 | ns | 0.0755<br>22 | 0.9274<br>79 | ns |
| E<br>W<br>T | Rhododendron<br>catawbiense | Mature | Week_covariate-All_season | 0.9729<br>42 | ns | 0.0755<br>22 | 0.9274<br>79 | ns |
| E<br>W<br>T | Rhododendron<br>catawbiense | Mature | Week_covariate-Measured | 0.9853<br>02 | ns | 0.0755<br>22 | 0.9274<br>79 | ns |
| E<br>W<br>T | Rhododendron<br>catawbiense | Old/Senes<br>cent | Measured-All_season | 0.1729<br>31 | ns | 1.8651<br>67 | 0.1766<br>34 | ns |
| E<br>W<br>T | Rhododendron<br>catawbiense | Old/Senes<br>cent | Week_covariate-All_season | 0.8858<br>54 | ns | 1.8651<br>67 | 0.1766<br>34 | ns |
| E<br>W<br>T | Rhododendron<br>catawbiense | Old/Senes<br>cent | Week_covariate-Measured | 0.3631<br>85 | ns | 1.8651<br>67 | 0.1766<br>34 | ns |
| E<br>W<br>T | Acer platanoides | Young | Measured-All_season | 0.3704<br>11 | ns | 1.4459<br>12 | 0.2553<br>24 | ns |
| E<br>W<br>T | Acer platanoides | Young | Week_covariate-All_season | 0.9818<br>57 | ns | 1.4459<br>12 | 0.2553<br>24 | ns |
| E<br>W<br>T | Acer platanoides | Young | Week_covariate-Measured | 0.2839<br>59 | ns | 1.4459<br>12 | 0.2553<br>24 | ns |
| E<br>W<br>T | Acer platanoides | Mature | Measured-All_season | 0.2462<br>28 | ns | 1.7760<br>12 | 0.1908<br>5 | ns |
| E<br>W<br>T | Acer platanoides | Mature | Week_covariate-All_season | 0.9995<br>37 | ns | 1.7760<br>12 | 0.1908<br>5 | ns |
| E<br>W<br>T | Acer platanoides | Mature | Week_covariate-Measured | 0.2578<br>23 | ns | 1.7760<br>12 | 0.1908<br>5 | ns |
| E<br>W<br>T | Acer platanoides | Old/Senes-<br>cent | Measured-All_season | 0.4883<br>7 | ns | 1.7236<br>84 | 0.1997<br>68 | ns |
| E<br>W<br>T | Acer platanoides | Old/Senes-<br>cent | Week_covariate-All_season | 0.7791<br>28 | ns | 1.7236<br>84 | 0.1997<br>68 | ns |
| E<br>W<br>T | Acer platanoides | Old/Senes-<br>cent | Week_covariate-Measured | 0.1795<br>93 | ns | 1.7236<br>84 | 0.1997<br>68 | ns |
| N | Quercus rubra | Young | Measured-All_season | 0.9380<br>1 | ns | 2.0176<br>89 | 0.1549<br>02 | ns |
| N | Quercus rubra | Young | Week_covariate-All_season | 0.2890<br>11 | ns | 2.0176<br>89 | 0.1549<br>02 | ns |
| N | Quercus rubra | Young | Week_covariate-Measured | 0.1647<br>43 | ns | 2.0176<br>89 | 0.1549<br>02 | ns |
| N | Quercus rubra | Mature | Measured-All_season | 0.3066<br>1 | ns | 3.4286<br>45 | 0.0490<br>05 | * |
| N | Quercus rubra | Mature | Week_covariate-All_season | 0.5210<br>11 | ns | 3.4286<br>45 | 0.0490<br>05 | * |
| N | Quercus rubra | Mature | Week_covariate-Measured | 0.0394<br>54 | * | 3.4286<br>45 | 0.0490<br>05 | * |
| N | Quercus rubra | Old/Senes-<br>cent | Measured-All_season | 0.3829<br>41 | ns | 1.0582<br>56 | 0.3627<br>04 | ns |
| N | Quercus rubra | Old/Senes-<br>cent | Week_covariate-All_season | 0.4957<br>15 | ns | 1.0582<br>56 | 0.3627<br>04 | ns |
| N | Quercus rubra | Old/Senes-<br>cent | Week_covariate-Measured | 0.9777<br>03 | ns | 1.0582<br>56 | 0.3627<br>04 | ns |
| N | Betula papyrifera | Young | Measured-All_season | 0.0007<br>13 | *** | 12.319<br>21 | 0.0002<br>08 | *** |
| N | Betula papyrifera | Young | Week_covariate-All_season | 0.9998<br>4 | ns | 12.319<br>21 | 0.0002<br>08 | *** |
| N | Betula papyrifera | Young | Week_covariate-Measured | 0.0006<br>84 | *** | 12.319<br>21 | 0.0002<br>08 | *** |
| N | Betula papyrifera | Mature | Measured-All_season | 0.1421<br>72 | ns | 3.8118<br>65 | 0.0365<br>07 | * |
| N | Betula papyrifera | Mature | Week_covariate-All_season | 0.7690<br>26 | ns | 3.8118<br>65 | 0.0365<br>07 | * |
| N | Betula papyrifera | Mature | Week_covariate-Measured | 0.0351<br>64 | * | 3.8118<br>65 | 0.0365<br>07 | * |
| N | Betula papyrifera | Old/Senes-<br>cent | Measured-All_season | 0.7720<br>74 | ns | 0.4624<br>86 | 0.6352<br>12 | ns |
| N | Betula papyrifera | Old/Senes-<br>cent | Week_covariate-All_season | 0.6297<br>52 | ns | 0.4624<br>86 | 0.6352<br>12 | ns |
| N | Betula papyrifera | Old/Senescent | Week_covariate-Measured | 0.969663 | ns | 0.462486 | 0.635212 | ns |
| N | Prunus nigra | Young | Measured-All_season | 0.497668 | ns | 0.928039 | 0.409056 | ns |
| N | Prunus nigra | Young | Week_covariate-All_season | 0.456984 | ns | 0.928039 | 0.409056 | ns |
| N | Prunus nigra | Young | Week_covariate-Measured | 0.997286 | ns | 0.928039 | 0.409056 | ns |
| N | Prunus nigra | Mature | Measured-All_season | 0.986446 | ns | 0.012918 | 0.987172 | ns |
| N | Prunus nigra | Mature | Week_covariate-All_season | 0.998601 | ns | 0.012918 | 0.987172 | ns |
| N | Prunus nigra | Mature | Week_covariate-Measured | 0.99371 | ns | 0.012918 | 0.987172 | ns |
| N | Prunus nigra | Old/Senescent | Measured-All_season | 5.50E-05 | *** | 17.94653 | 5.17E-05 | *** |
| N | Prunus nigra | Old/Senescent | Week_covariate-All_season | 0.373043 | ns | 17.94653 | 5.17E-05 | *** |
| N | Prunus nigra | Old/Senescent | Week_covariate-Measured | 0.001045 | ** | 17.94653 | 5.17E-05 | *** |
| N | Acer rubrum | Young | Measured-All_season | 0.178031 | ns | 1.707984 | 0.202531 | ns |
| N | Acer rubrum | Young | Week_covariate-All_season | 0.720825 | ns | 1.707984 | 0.202531 | ns |
| N | Acer rubrum | Young | Week_covariate-Measured | 0.544741 | ns | 1.707984 | 0.202531 | ns |
| N | Acer rubrum | Mature | Measured-All_season | 0.000155 | *** | 23.3499 | 2.34E-06 | *** |
| N | Acer rubrum | Mature | Week_covariate-All_season | 0.2325<br>61 | ns | 23.349<br>9 | 2.34E-06 | *** |
| N | Acer rubrum | Mature | Week_covariate-Measured | 2.45E-06 | *** | 23.349<br>9 | 2.34E-06 | *** |
| N | Acer rubrum | Old/Senes<br>cent | Measured-All_season | 0.9639<br>54 | ns | 2.7965<br>06 | 0.0809<br>59 | ns |
| N | Acer rubrum | Old/Senes<br>cent | Week_covariate-All_season | 0.1585<br>17 | ns | 2.7965<br>06 | 0.0809<br>59 | ns |
| N | Acer rubrum | Old/Senes<br>cent | Week_covariate-Measured | 0.0980<br>16 | ns | 2.7965<br>06 | 0.0809<br>59 | ns |
| N | Rhododendron<br>maximum | Young | Measured-All_season | 0.1722<br>9 | ns | 3.2586<br>92 | 0.0559<br>72 | ns |
| N | Rhododendron<br>maximum | Young | Week_covariate-All_season | 0.8293<br>71 | ns | 3.2586<br>92 | 0.0559<br>72 | ns |
| N | Rhododendron<br>maximum | Young | Week_covariate-Measured | 0.0558<br>39 | ns | 3.2586<br>92 | 0.0559<br>72 | ns |
| N | Rhododendron<br>maximum | Mature | Measured-All_season | 0.9174<br>04 | ns | 0.3050<br>75 | 0.7398<br>85 | ns |
| N | Rhododendron<br>maximum | Mature | Week_covariate-All_season | 0.9218<br>55 | ns | 0.3050<br>75 | 0.7398<br>85 | ns |
| N | Rhododendron<br>maximum | Mature | Week_covariate-Measured | 0.7179<br>89 | ns | 0.3050<br>75 | 0.7398<br>85 | ns |
| N | Rhododendron<br>maximum | Old/Senes<br>cent | Measured-All_season | 0.9381<br>71 | ns | 1.7389<br>26 | 0.1971<br>25 | ns |
| N | Rhododendron<br>maximum | Old/Senes<br>cent | Week_covariate-All_season | 0.2049<br>88 | ns | 1.7389<br>26 | 0.1971<br>25 | ns |
| N | Rhododendron<br>maximum | Old/Senes<br>cent | Week_covariate-Measured | 0.3480<br>41 | ns | 1.7389<br>26 | 0.1971<br>25 | ns |
| N | Rhododendron catawbiense | Young | Measured-All_season | 0.1002<br>41 | ns | 11.622<br>51 | 0.0002<br>95 | *** |
| N | Rhododendron catawbiense | Young | Week_covariate-All_season | 0.0353<br>53 | * | 11.622<br>51 | 0.0002<br>95 | *** |
| N | Rhododendron catawbiense | Young | Week_covariate-Measured | 0.0001<br>91 | *** | 11.622<br>51 | 0.0002<br>95 | *** |
| N | Rhododendron catawbiense | Mature | Measured-All_season | 0.0228<br>38 | * | 5.8655<br>12 | 0.0084<br>33 | ** |
| N | Rhododendron catawbiense | Mature | Week_covariate-All_season | 0.9763<br>01 | ns | 5.8655<br>12 | 0.0084<br>33 | ** |
| N | Rhododendron catawbiense | Mature | Week_covariate-Measured | 0.0141<br>75 | * | 5.8655<br>12 | 0.0084<br>33 | ** |
| N | Rhododendron catawbiense | Old/Senescent | Measured-All_season | 0.1599<br>81 | ns | 2.0466<br>95 | 0.1511<br>07 | ns |
| N | Rhododendron catawbiense | Old/Senescent | Week_covariate-All_season | 0.2866<br>58 | ns | 2.0466<br>95 | 0.1511<br>07 | ns |
| N | Rhododendron catawbiense | Old/Senescent | Week_covariate-Measured | 0.9340<br>32 | ns | 2.0466<br>95 | 0.1511<br>07 | ns |
| N | Acer platanoides | Young | Measured-All_season | 0.9116<br>48 | ns | 0.1396<br>02 | 0.8704<br>05 | ns |
| N | Acer platanoides | Young | Week_covariate-All_season | 0.8751<br>17 | ns | 0.1396<br>02 | 0.8704<br>05 | ns |
| N | Acer platanoides | Young | Week_covariate-Measured | 0.9962<br>17 | ns | 0.1396<br>02 | 0.8704<br>05 | ns |
| N | Acer platanoides | Mature | Measured-All_season | 0.8951<br>69 | ns | 0.1010<br>42 | 0.9042<br>77 | ns |
| N | Acer platanoides | Mature | Week_covariate-All_season | 0.9696<br>04 | ns | 0.1010<br>42 | 0.9042<br>77 | ns |
| N | Acer platanoides | Mature | Week_covariate-Measured | 0.9754<br>31 | ns | 0.1010<br>42 | 0.9042<br>77 | ns |
| N | Acer platanoides | Old/Senescent | Measured-All_season | 0.0451<br>52 | * | 3.4286<br>58 | 0.0490<br>05 | * |
| N | Acer platanoides | Old/Senescent | Week_covariate-All_season | 0.7422<br>96 | ns | 3.4286<br>58 | 0.0490<br>05 | * |
| N | Acer platanoides | Old/Senescent | Week_covariate-Measured | 0.1892<br>87 | ns | 3.4286<br>58 | 0.0490<br>05 | * |
| C | Quercus rubra | Young | Measured-All_season | 0.0002<br>32 | *** | 90.967<br>91 | 6.28E-12 | *** |
| C | Quercus rubra | Young | Week_covariate-All_season | 2.70E-08 | *** | 90.967<br>91 | 6.28E-12 | *** |
| C | Quercus rubra | Young | Week_covariate-Measured | 4.25E-12 | *** | 90.967<br>91 | 6.28E-12 | *** |
| C | Quercus rubra | Mature | Measured-All_season | 3.24E-05 | *** | 18.242<br>77 | 1.52E-05 | *** |
| C | Quercus rubra | Mature | Week_covariate-All_season | 0.8005<br>11 | ns | 18.242<br>77 | 1.52E-05 | *** |
| C | Quercus rubra | Mature | Week_covariate-Measured | 0.0001<br>6 | *** | 18.242<br>77 | 1.52E-05 | *** |
| C | Quercus rubra | Old/Senescent | Measured-All_season | 3.97E-05 | *** | 20.878<br>62 | 5.59E-06 | *** |
| C | Quercus rubra | Old/Senescent | Week_covariate-All_season | 1.87E-05 | *** | 20.878<br>62 | 5.59E-06 | *** |
| C | Quercus rubra | Old/Senescent | Week_covariate-Measured | 0.9509<br>46 | ns | 20.878<br>62 | 5.59E-06 | *** |
| C | Betula papyrifera | Young | Measured-All_season | 0.0180<br>48 | * | 6.1351<br>3 | 0.0070<br>46 | ** |
| C | Betula papyrifera | Young | Week_covariate-All_season | 0.0130<br>05 | * | 6.1351<br>3 | 0.0070<br>46 | ** |
| C | Betula papyrifera | Young | Week_covariate-Measured | 0.9889<br>52 | ns | 6.1351<br>3 | 0.0070<br>46 | ** |
| C | Betula papyrifera | Mature | Measured-All_season | 0.1210<br>07 | ns | 2.5988<br>44 | 0.0951<br>38 | ns |
| C | Betula papyrifera | Mature | Week_covariate-All_season | 0.9839<br>74 | ns | 2.5988<br>44 | 0.0951<br>38 | ns |
| C | Betula papyrifera | Mature | Week_covariate-Measured | 0.1652<br>7 | ns | 2.5988<br>44 | 0.0951<br>38 | ns |
| C | Betula papyrifera | Old/Senes-<br>cent | Measured-All_season | 1.17E-09 | *** | 51.881<br>69 | 1.93E-09 | *** |
| C | Betula papyrifera | Old/Senes-<br>cent | Week_covariate-All_season | 7.52E-06 | *** | 51.881<br>69 | 1.93E-09 | *** |
| C | Betula papyrifera | Old/Senes-<br>cent | Week_covariate-Measured | 0.0014<br>74 | ** | 51.881<br>69 | 1.93E-09 | *** |
| C | Prunus nigra | Young | Measured-All_season | 3.71E-13 | *** | 232.79<br>17 | 1.93E-16 | *** |
| C | Prunus nigra | Young | Week_covariate-All_season | 2.12E-14 | *** | 232.79<br>17 | 1.93E-16 | *** |
| C | Prunus nigra | Young | Week_covariate-Measured | 9.58E-06 | *** | 232.79<br>17 | 1.93E-16 | *** |
| C | Prunus nigra | Mature | Measured-All_season | 0.9669<br>62 | ns | 0.0731<br>04 | 0.9297<br>11 | ns |
| C | Prunus nigra | Mature | Week_covariate-All_season | 0.9251<br>68 | ns | 0.0731<br>04 | 0.9297<br>11 | ns |
| C | Prunus nigra | Mature | Week_covariate-Measured | 0.9908<br>39 | ns | 0.0731<br>04 | 0.9297<br>11 | ns |
| C | Prunus nigra | Old/Senescent | Measured-All_season | 0.038836 | * | 18.86918 | 3.82E-05 | *** |
| C | Prunus nigra | Old/Senescent | Week_covariate-All_season | 2.48E-05 | *** | 18.86918 | 3.82E-05 | *** |
| C | Prunus nigra | Old/Senescent | Week_covariate-Measured | 0.00767 | ** | 18.86918 | 3.82E-05 | *** |
| C | Acer rubrum | Young | Measured-All_season | 0.030424 | * | 3.727175 | 0.038937 | * |
| C | Acer rubrum | Young | Week_covariate-All_season | 0.323618 | ns | 3.727175 | 0.038937 | * |
| C | Acer rubrum | Young | Week_covariate-Measured | 0.430999 | ns | 3.727175 | 0.038937 | * |
| C | Acer rubrum | Mature | Measured-All_season | 0.000555 | *** | 12.02103 | 0.000242 | *** |
| C | Acer rubrum | Mature | Week_covariate-All_season | 0.950665 | ns | 12.02103 | 0.000242 | *** |
| C | Acer rubrum | Mature | Week_covariate-Measured | 0.001188 | ** | 12.02103 | 0.000242 | *** |
| C | Acer rubrum | Old/Senescent | Measured-All_season | 0.280627 | ns | 4.25767 | 0.02615 | * |
| C | Acer rubrum | Old/Senescent | Week_covariate-All_season | 0.381035 | ns | 4.25767 | 0.02615 | * |
| C | Acer rubrum | Old/Senescent | Week_covariate-Measured | 0.019975 | * | 4.25767 | 0.02615 | * |
| C | Rhododendron maximum | Young | Measured-All_season | 0.298951 | ns | 2.341788 | 0.117743 | ns |
| C | Rhododendron maximum | Young | Week_covariate-All_season | 0.112396 | ns | 2.341788 | 0.117743 | ns |
| C | Rhododendron maximum | Young | Week_covariate-Measured | 0.836106 | ns | 2.341788 | 0.117743 | ns |
| C | Rhododendron maximum | Mature | Measured-All_season | 0.000368 | *** | 15.22375 | 5.38E-05 | *** |
| C | Rhododendron maximum | Mature | Week_covariate-All_season | 0.906412 | ns | 15.22375 | 5.38E-05 | *** |
| C | Rhododendron maximum | Mature | Week_covariate-Measured | 0.000127 | *** | 15.22375 | 5.38E-05 | *** |
| C | Rhododendron maximum | Old/Senescent | Measured-All_season | 5.43E-09 | *** | 44.98046 | 7.61E-09 | *** |
| C | Rhododendron maximum | Old/Senescent | Week_covariate-All_season | 7.59E-06 | *** | 44.98046 | 7.61E-09 | *** |
| C | Rhododendron maximum | Old/Senescent | Week_covariate-Measured | 0.009653 | ** | 44.98046 | 7.61E-09 | *** |
| C | Rhododendron catawbiense | Young | Measured-All_season | 0.034889 | * | 891.7525 | 3.00E-23 | *** |
| C | Rhododendron catawbiense | Young | Week_covariate-All_season | 2.10E-14 | *** | 891.7525 | 3.00E-23 | *** |
| C | Rhododendron catawbiense | Young | Week_covariate-Measured | 2.10E-14 | *** | 891.7525 | 3.00E-23 | *** |
| C | Rhododendron catawbiense | Mature | Measured-All_season | 0.996356 | ns | 0.03355 | 0.967052 | ns |
| C | Rhododendron catawbiense | Mature | Week_covariate-All_season | 0.965202 | ns | 0.03355 | 0.967052 | ns |
| C | Rhododendron catawbiense | Mature | Week_covariate-Measured | 0.98378 | ns | 0.03355 | 0.967052 | ns |
| C | Rhododendron catawbiense | Old/Senescent | Measured-All_season | 0.054144 | ns | 3.5108 | 0.04598 | * |
| C | Rhododendron<br>catawbiense | Old/Senes<br>cent | Week_covariate-All_season | 0.9255<br>32 | ns | 3.5108 | 0.0459<br>8 | * |
| C | Rhododendron<br>catawbiense | Old/Senes<br>cent | Week_covariate-Measured | 0.1144<br>53 | ns | 3.5108 | 0.0459<br>8 | * |
| C | Acer platanoides | Young | Measured-All_season | 0.0581<br>39 | ns | 6.9320<br>65 | 0.0042<br>05 | ** |
| C | Acer platanoides | Young | Week_covariate-All_season | 0.0034<br>23 | ** | 6.9320<br>65 | 0.0042<br>05 | ** |
| C | Acer platanoides | Young | Week_covariate-Measured | 0.4451<br>55 | ns | 6.9320<br>65 | 0.0042<br>05 | ** |
| C | Acer platanoides | Mature | Measured-All_season | 1.58E-09 | *** | 65.104<br>44 | 9.95E-10 | *** |
| C | Acer platanoides | Mature | Week_covariate-All_season | 0.3676<br>22 | ns | 65.104<br>44 | 9.95E-10 | *** |
| C | Acer platanoides | Mature | Week_covariate-Measured | 1.35E-08 | *** | 65.104<br>44 | 9.95E-10 | *** |
| C | Acer platanoides | Old/Senes<br>cent | Measured-All_season | 0.0680<br>95 | ns | 4.3786<br>28 | 0.0239<br>25 | * |
| C | Acer platanoides | Old/Senes<br>cent | Week_covariate-All_season | 0.0300<br>18 | * | 4.3786<br>28 | 0.0239<br>25 | * |
| C | Acer platanoides | Old/Senes<br>cent | Week_covariate-Measured | 0.9216<br>2 | ns | 4.3786<br>28 | 0.0239<br>25 | * |
| Q<br>Y | Quercus rubra | Young | Measured-All_season | 0.0028<br>68 | ** | 13.517<br>21 | 0.0001<br>17 | *** |
| Q<br>Y | Quercus rubra | Young | Week_covariate-All_season | 0.4253<br>09 | ns | 13.517<br>21 | 0.0001<br>17 | *** |
| Q<br>Y | Quercus rubra | Young | Week_covariate-Measured | 0.0001<br>19 | *** | 13.517<br>21 | 0.0001<br>17 | *** |
| Q<br>Y | Quercus rubra | Mature | Measured-All_season | 3.32E-08 | *** | 41.851<br>99 | 1.50E-08 | *** |
| Q<br>Y | Quercus rubra | Mature | Week_covariate-All_season | 0.4388<br>48 | ns | 41.851<br>99 | 1.50E-08 | *** |
| Q<br>Y | Quercus rubra | Mature | Week_covariate-Measured | 5.33E-07 | *** | 41.851<br>99 | 1.50E-08 | *** |
| Q<br>Y | Quercus rubra | Old/Senes-<br>cent | Measured-All_season | 0.0169<br>78 | * | 7.3718<br>33 | 0.0031<br>92 | ** |
| Q<br>Y | Quercus rubra | Old/Senes-<br>cent | Week_covariate-All_season | 0.8233<br>67 | ns | 7.3718<br>33 | 0.0031<br>92 | ** |
| Q<br>Y | Quercus rubra | Old/Senes-<br>cent | Week_covariate-Measured | 0.0041<br>25 | ** | 7.3718<br>33 | 0.0031<br>92 | ** |
| Q<br>Y | Betula papyrifera | Young | Measured-All_season | 0.3742<br>52 | ns | 1.0865<br>34 | 0.3534<br>1 | ns |
| Q<br>Y | Betula papyrifera | Young | Week_covariate-All_season | 0.9777<br>8 | ns | 1.0865<br>34 | 0.3534<br>1 | ns |
| Q<br>Y | Betula papyrifera | Young | Week_covariate-Measured | 0.4856<br>97 | ns | 1.0865<br>34 | 0.3534<br>1 | ns |
| Q<br>Y | Betula papyrifera | Mature | Measured-All_season | 3.38E-10 | *** | 92.333<br>28 | 5.36E-12 | *** |
| Q<br>Y | Betula papyrifera | Mature | Week_covariate-All_season | 0.1915<br>79 | ns | 92.333<br>28 | 5.36E-12 | *** |
| Q<br>Y | Betula papyrifera | Mature | Week_covariate-Measured | 1.43E-11 | *** | 92.333<br>28 | 5.36E-12 | *** |
| Q<br>Y | Betula papyrifera | Old/Senes-<br>cent | Measured-All_season | 0.9998<br>67 | ns | 0.5403 | 0.5894<br>94 | ns |
| Q<br>Y | Betula papyrifera | Old/Senes-<br>cent | Week_covariate-All_season | 0.6406<br>33 | ns | 0.5403 | 0.5894<br>94 | ns |
| Q<br>Y | Betula papyrifera | Old/Senescent | Week_covariate-Measured | 0.6502<br>11 | ns | 0.5403 | 0.5894<br>94 | ns |
| Q<br>Y | Prunus nigra | Young | Measured-All_season | 0.0045<br>28 | ** | 7.7075<br>9 | 0.0025<br>98 | ** |
| Q<br>Y | Prunus nigra | Young | Week_covariate-All_season | 0.9483<br>37 | ns | 7.7075<br>9 | 0.0025<br>98 | ** |
| Q<br>Y | Prunus nigra | Young | Week_covariate-Measured | 0.0095<br>37 | ** | 7.7075<br>9 | 0.0025<br>98 | ** |
| Q<br>Y | Prunus nigra | Mature | Measured-All_season | 0.0003<br>03 | *** | 26.371<br>08 | 8.75E-07 | *** |
| Q<br>Y | Prunus nigra | Mature | Week_covariate-All_season | 0.0466<br>06 | * | 26.371<br>08 | 8.75E-07 | *** |
| Q<br>Y | Prunus nigra | Mature | Week_covariate-Measured | 6.21E-07 | *** | 26.371<br>08 | 8.75E-07 | *** |
| Q<br>Y | Prunus nigra | Old/Senescent | Measured-All_season | 0.3140<br>92 | ns | 1.1388<br>24 | 0.3422<br>11 | ns |
| Q<br>Y | Prunus nigra | Old/Senescent | Week_covariate-All_season | 0.6535<br>69 | ns | 1.1388<br>24 | 0.3422<br>11 | ns |
| Q<br>Y | Prunus nigra | Old/Senescent | Week_covariate-Measured | 0.8159<br>91 | ns | 1.1388<br>24 | 0.3422<br>11 | ns |
| Q<br>Y | Acer rubrum | Young | Measured-All_season | 1.63E-06 | *** | 22.765<br>34 | 2.86E-06 | *** |
| Q<br>Y | Acer rubrum | Young | Week_covariate-All_season | 0.0067<br>24 | ** | 22.765<br>34 | 2.86E-06 | *** |
| Q<br>Y | Acer rubrum | Young | Week_covariate-Measured | 0.0069<br>61 | ** | 22.765<br>34 | 2.86E-06 | *** |
| Q<br>Y | Acer rubrum | Mature | Measured-All_season | 0.0090<br>27 | ** | 5.4075<br>86 | 0.0115<br>16 | * |
| Q<br>Y | Acer rubrum | Mature | Week_covariate-All_season | 0.4432<br>9 | ns | 5.4075<br>86 | 0.0115<br>16 | * |
| Q<br>Y | Acer rubrum | Mature | Week_covariate-Measured | 0.1290<br>57 | ns | 5.4075<br>86 | 0.0115<br>16 | * |
| Q<br>Y | Acer rubrum | Old/Senes-<br>cent | Measured-All_season | 0.8791<br>2 | ns | 0.1177<br>52 | 0.8894<br>27 | ns |
| Q<br>Y | Acer rubrum | Old/Senes-<br>cent | Week_covariate-All_season | 0.9726<br>92 | ns | 0.1177<br>52 | 0.8894<br>27 | ns |
| Q<br>Y | Acer rubrum | Old/Senes-<br>cent | Week_covariate-Measured | 0.9633<br>14 | ns | 0.1177<br>52 | 0.8894<br>27 | ns |
| Q<br>Y | Rhododendron<br>maximum | Young | Measured-All_season | 0.7540<br>25 | ns | 2.0353<br>4 | 0.1525<br>81 | ns |
| Q<br>Y | Rhododendron<br>maximum | Young | Week_covariate-All_season | 0.4241<br>43 | ns | 2.0353<br>4 | 0.1525<br>81 | ns |
| Q<br>Y | Rhododendron<br>maximum | Young | Week_covariate-Measured | 0.1357<br>94 | ns | 2.0353<br>4 | 0.1525<br>81 | ns |
| Q<br>Y | Rhododendron<br>maximum | Mature | Measured-All_season | 0.0016<br>14 | ** | 8.7313<br>19 | 0.0014<br>15 | ** |
| Q<br>Y | Rhododendron<br>maximum | Mature | Week_covariate-All_season | 0.6848<br>03 | ns | 8.7313<br>19 | 0.0014<br>15 | ** |
| Q<br>Y | Rhododendron<br>maximum | Mature | Week_covariate-Measured | 0.0122<br>47 | * | 8.7313<br>19 | 0.0014<br>15 | ** |
| Q<br>Y | Rhododendron<br>maximum | Old/Senes-<br>cent | Measured-All_season | 0.9216<br>8 | ns | 0.5204<br>21 | 0.6008<br>25 | ns |
| Q<br>Y | Rhododendron<br>maximum | Old/Senes-<br>cent | Week_covariate-All_season | 0.5773<br>83 | ns | 0.5204<br>21 | 0.6008<br>25 | ns |
| Q<br>Y | Rhododendron<br>maximum | Old/Senes-<br>cent | Week_covariate-Measured | 0.8077<br>08 | ns | 0.5204<br>21 | 0.6008<br>25 | ns |
| Q<br>Y | Rhododendron<br>catawbiense | Young | Measured-All_season | 0.1053<br>82 | ns | 4.4531<br>71 | 0.0226<br>56 | * |
| Q<br>Y | Rhododendron<br>catawbiense | Young | Week_covariate-All_season | 0.7371<br>99 | ns | 4.4531<br>71 | 0.0226<br>56 | * |
| Q<br>Y | Rhododendron<br>catawbiense | Young | Week_covariate-Measured | 0.0218<br>35 | * | 4.4531<br>71 | 0.0226<br>56 | * |
| Q<br>Y | Rhododendron<br>catawbiense | Mature | Measured-All_season | 4.50E-07 | *** | 35.194<br>07 | 7.30E-08 | *** |
| Q<br>Y | Rhododendron<br>catawbiense | Mature | Week_covariate-All_season | 0.9974<br>1 | ns | 35.194<br>07 | 7.30E-08 | *** |
| Q<br>Y | Rhododendron<br>catawbiense | Mature | Week_covariate-Measured | 5.27E-07 | *** | 35.194<br>07 | 7.30E-08 | *** |
| Q<br>Y | Rhododendron<br>catawbiense | Old/Senes-<br>cent | Measured-All_season | 0.9320<br>72 | ns | 1.1931<br>92 | 0.3206<br>09 | ns |
| Q<br>Y | Rhododendron<br>catawbiense | Old/Senes-<br>cent | Week_covariate-All_season | 0.3178<br>09 | ns | 1.1931<br>92 | 0.3206<br>09 | ns |
| Q<br>Y | Rhododendron<br>catawbiense | Old/Senes-<br>cent | Week_covariate-Measured | 0.5098<br>53 | ns | 1.1931<br>92 | 0.3206<br>09 | ns |
| Q<br>Y | Acer platanoides | Young | Measured-All_season | 0.9803<br>18 | ns | 2.1433<br>59 | 0.1391<br>7 | ns |
| Q<br>Y | Acer platanoides | Young | Week_covariate-All_season | 0.2293<br>2 | ns | 2.1433<br>59 | 0.1391<br>7 | ns |
| Q<br>Y | Acer platanoides | Young | Week_covariate-Measured | 0.1660<br>68 | ns | 2.1433<br>59 | 0.1391<br>7 | ns |
| Q<br>Y | Acer platanoides | Mature | Measured-All_season | 0.0056<br>22 | ** | 10.302<br>85 | 0.0005<br>89 | *** |
| Q<br>Y | Acer platanoides | Mature | Week_covariate-All_season | 0.6935<br>18 | ns | 10.302<br>85 | 0.0005<br>89 | *** |
| Q<br>Y | Acer platanoides | Mature | Week_covariate-Measured | 0.0007<br>37 | *** | 10.302<br>85 | 0.0005<br>89 | *** |
| Q<br>Y | Acer platanoides | Old/Senes<br>cent | Measured-All_season | 0.8820<br>9 | ns | 0.4237<br>01 | 0.6594<br>21 | ns |
| Q<br>Y | Acer platanoides | Old/Senes<br>cent | Week_covariate-All_season | 0.8984<br>07 | ns | 0.4237<br>01 | 0.6594<br>21 | ns |
| Q<br>Y | Acer platanoides | Old/Senes<br>cent | Week_covariate-Measured | 0.6330<br>11 | ns | 0.4237<br>01 | 0.6594<br>21 | ns |

**Table S7.**
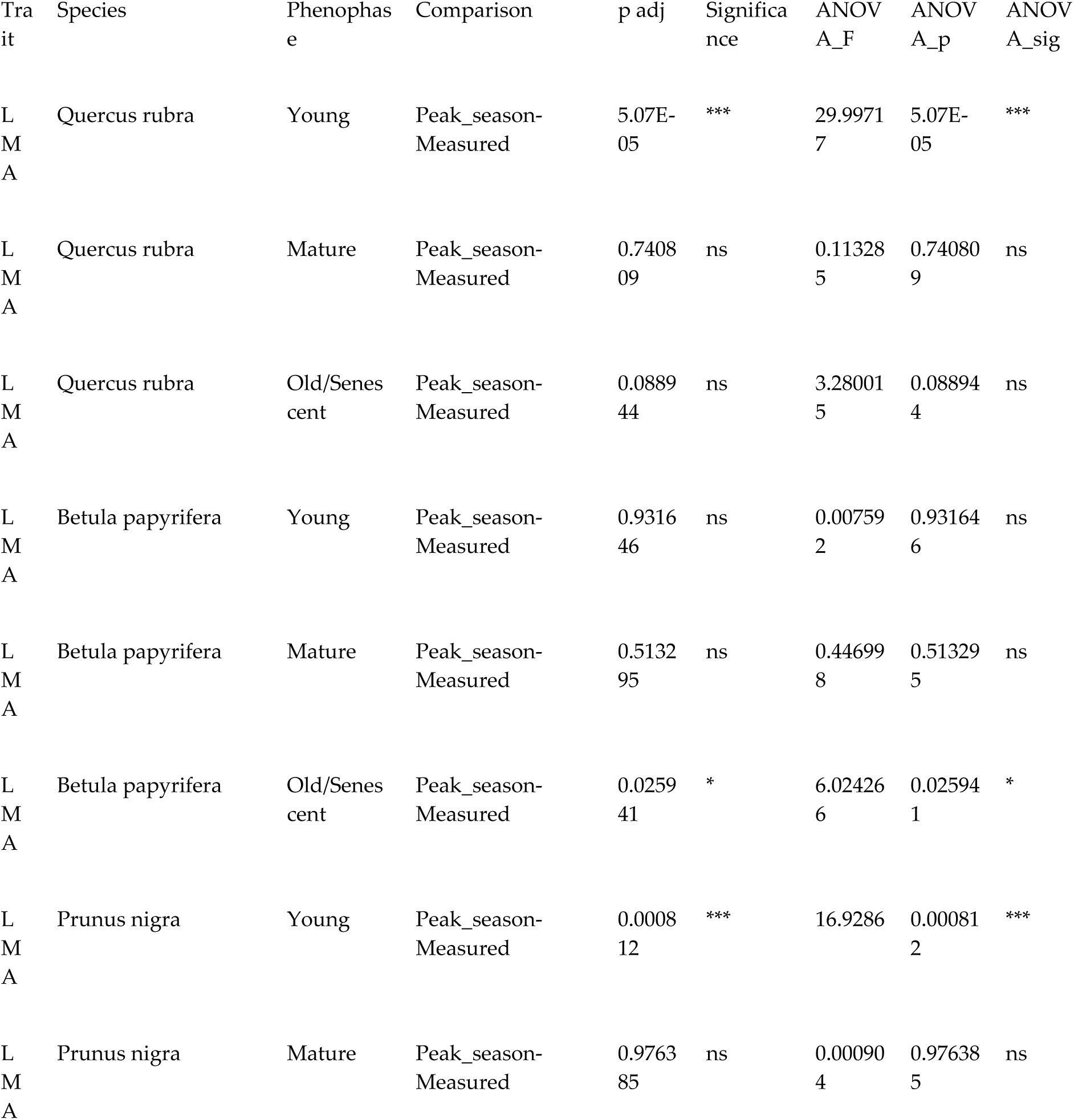
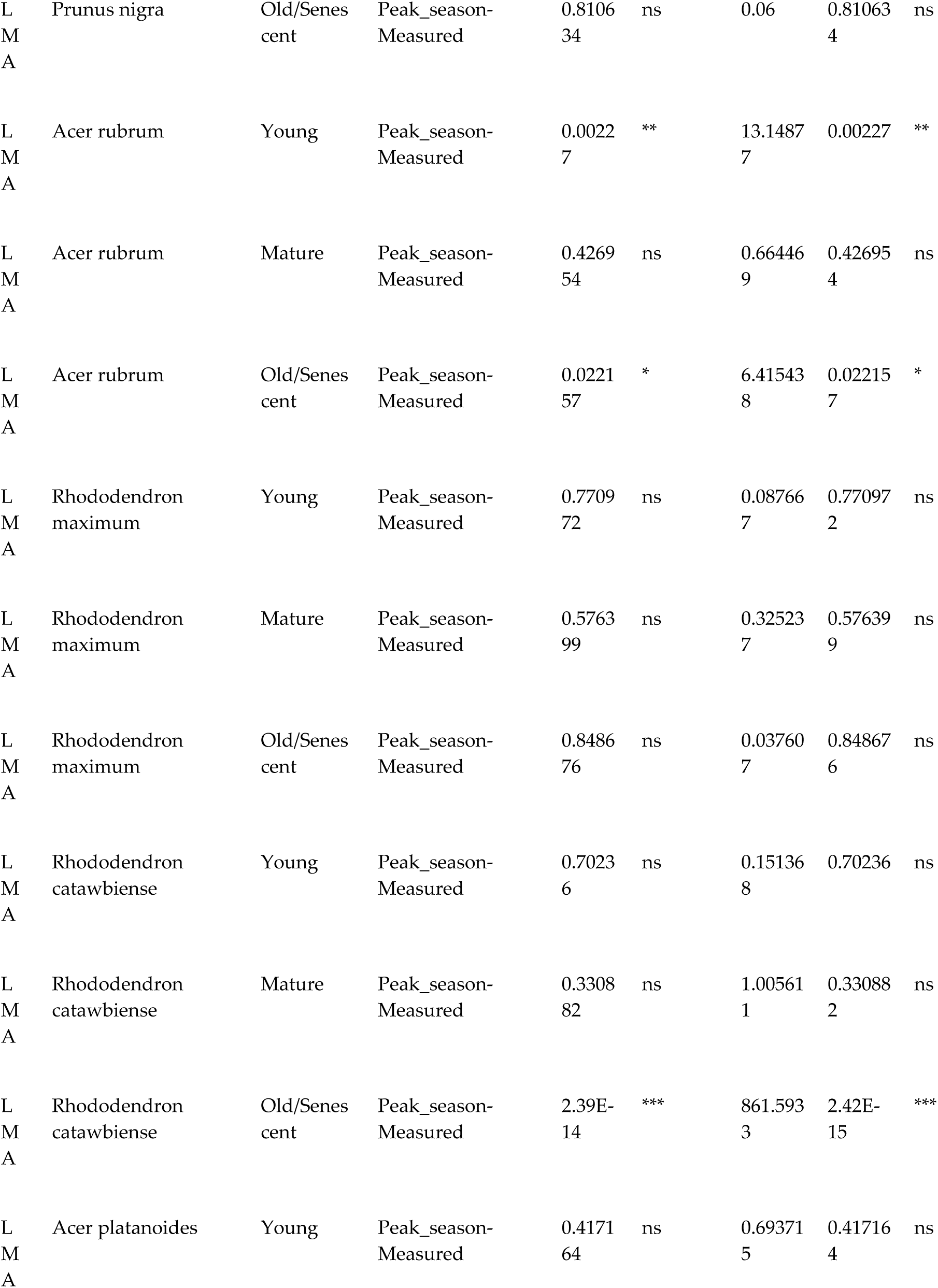
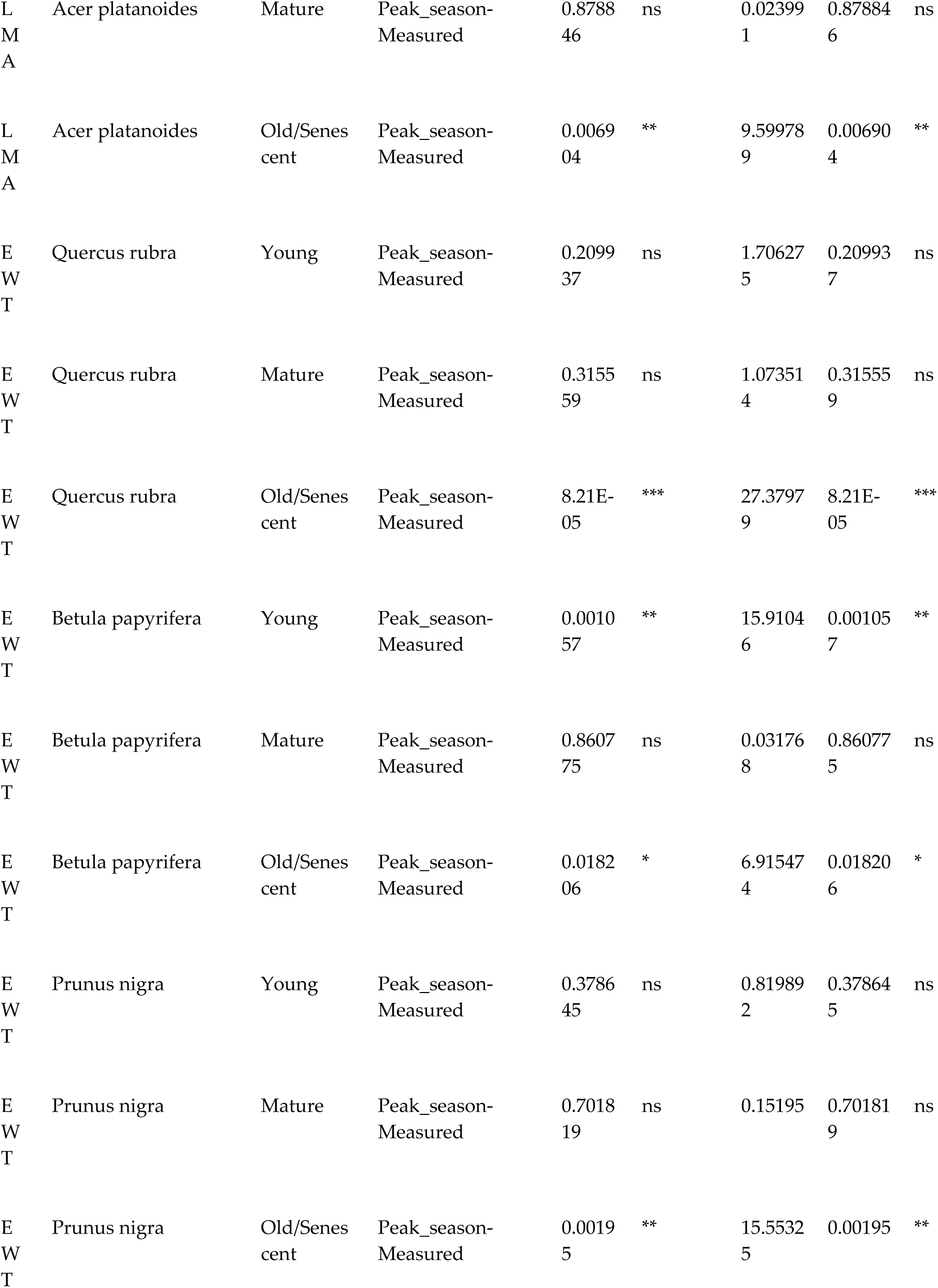
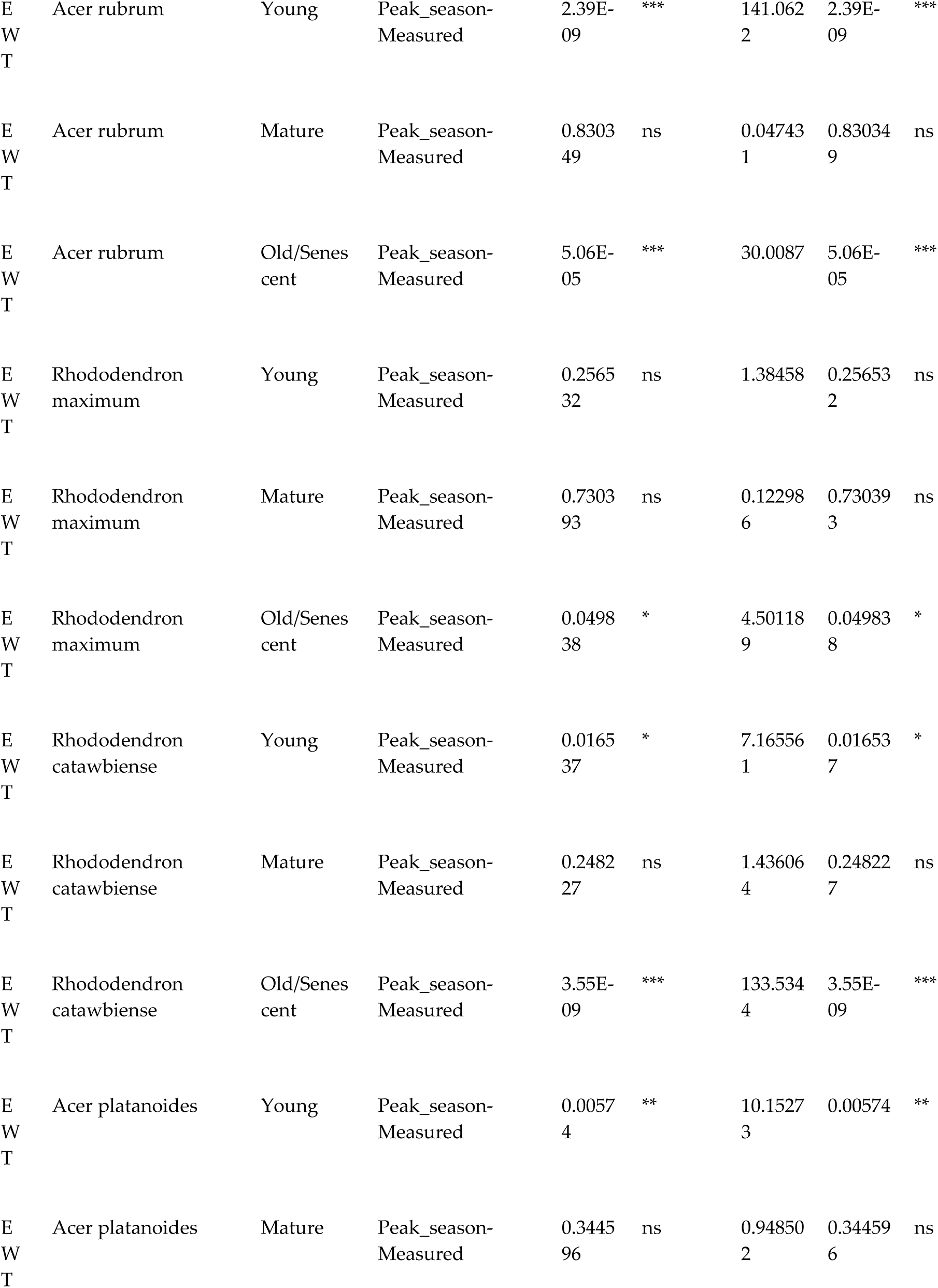
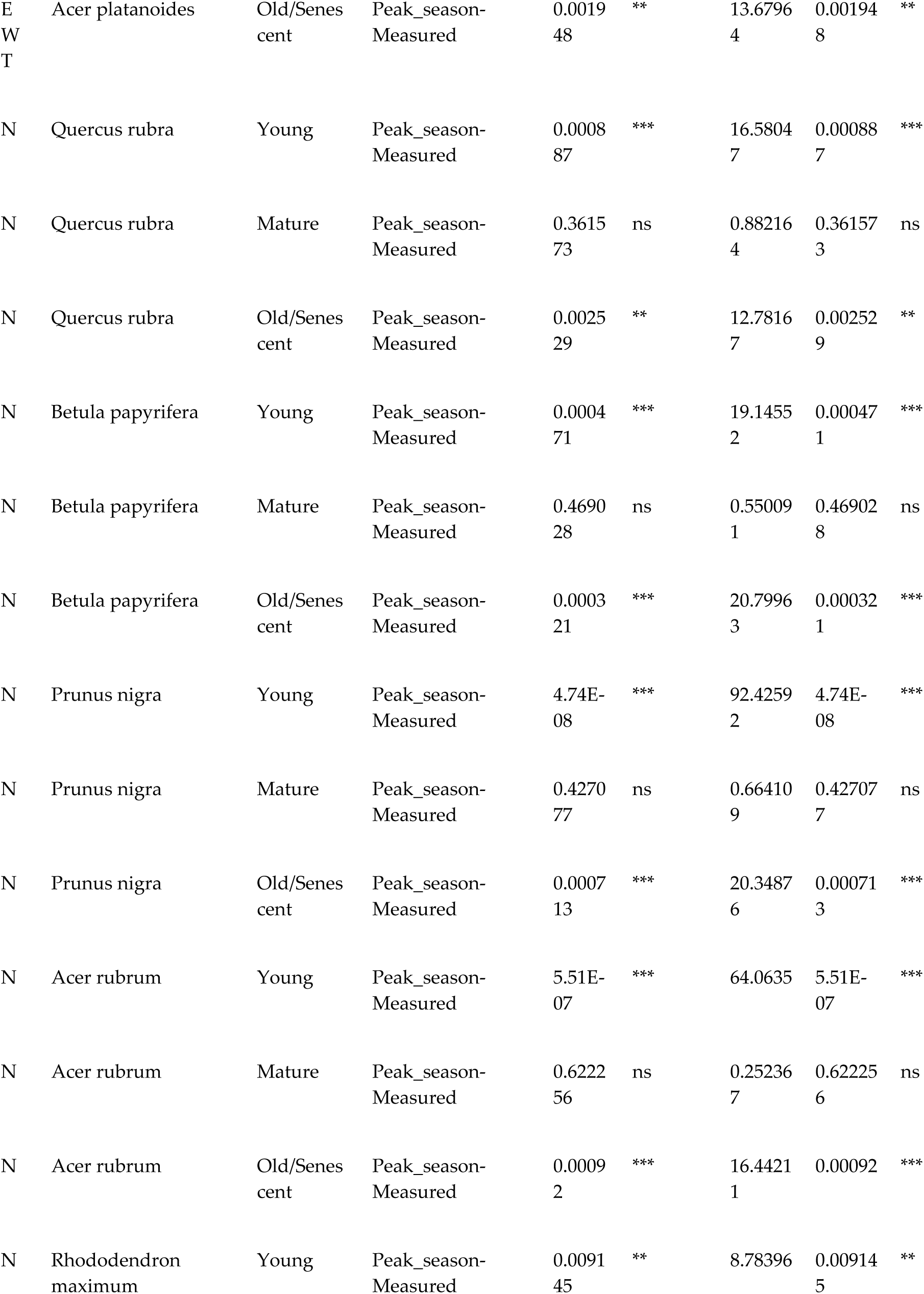
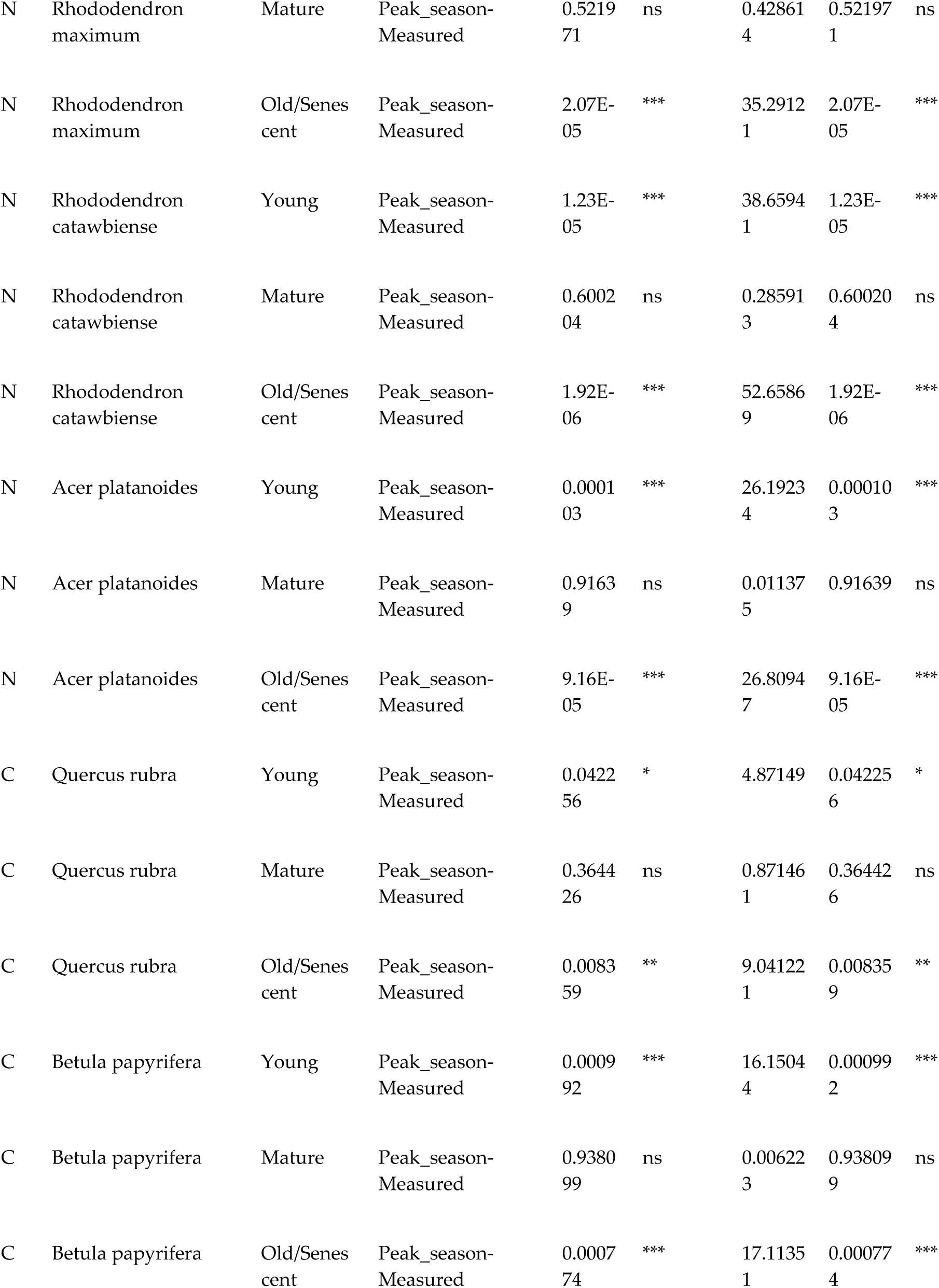
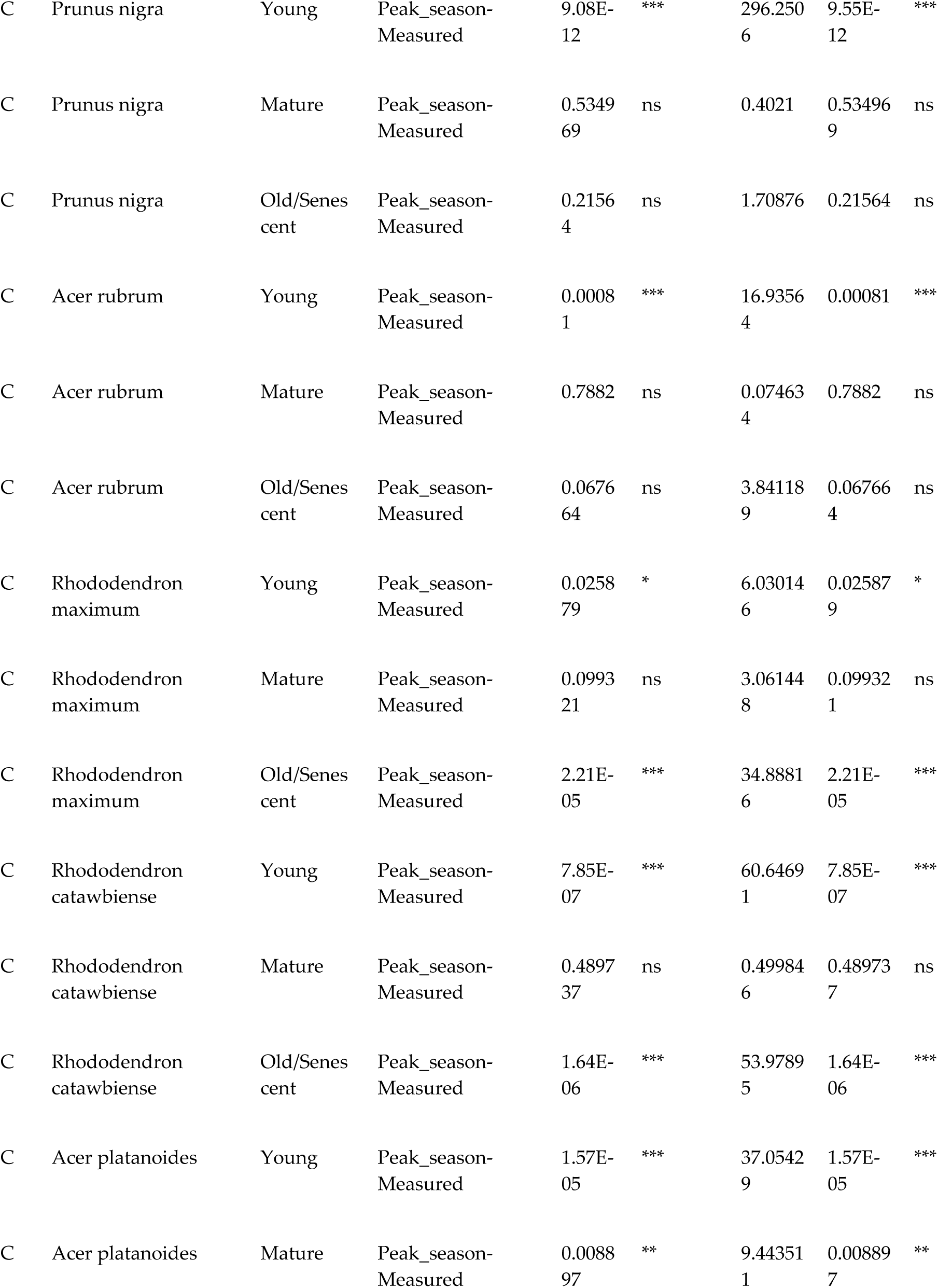
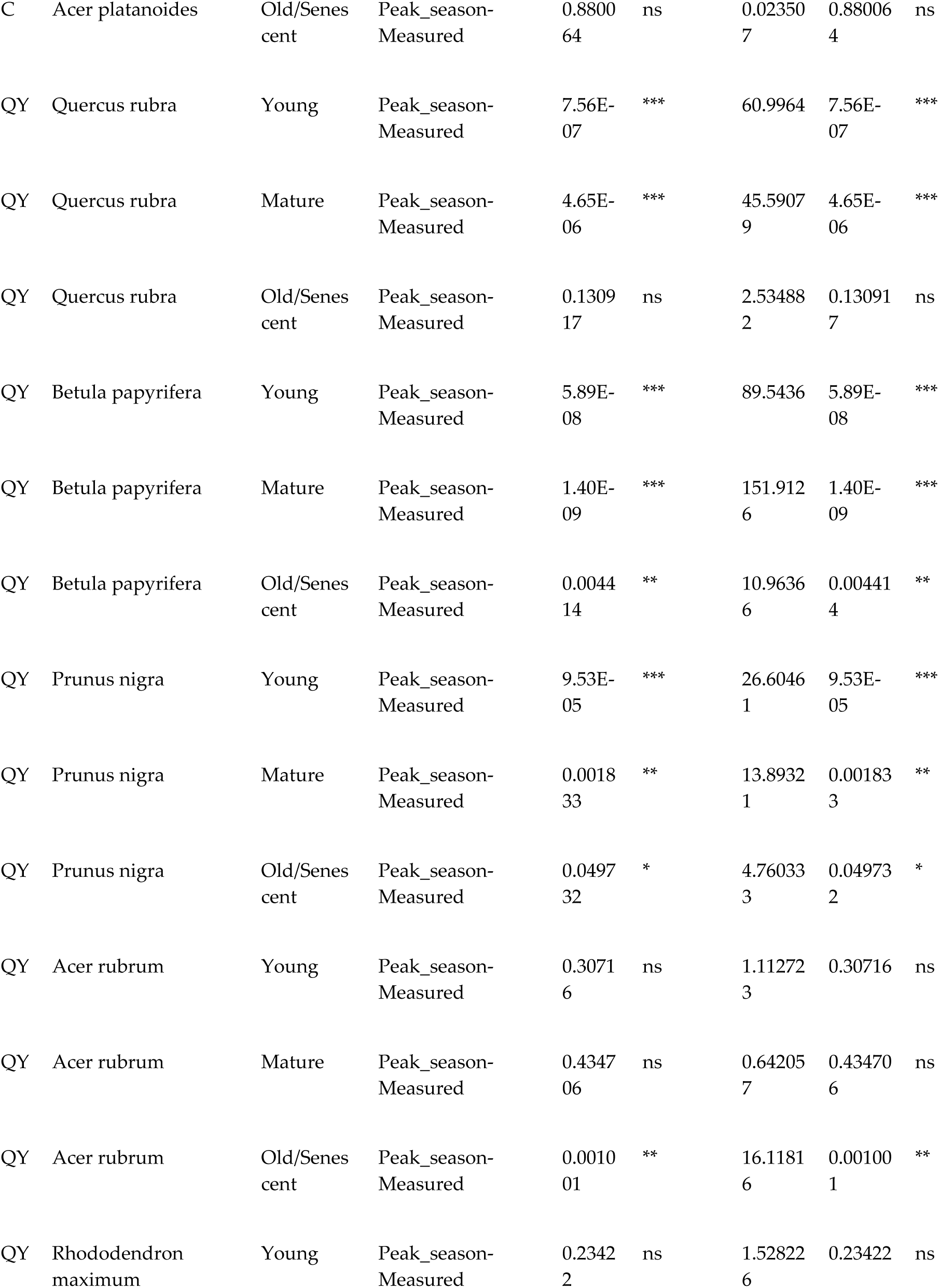
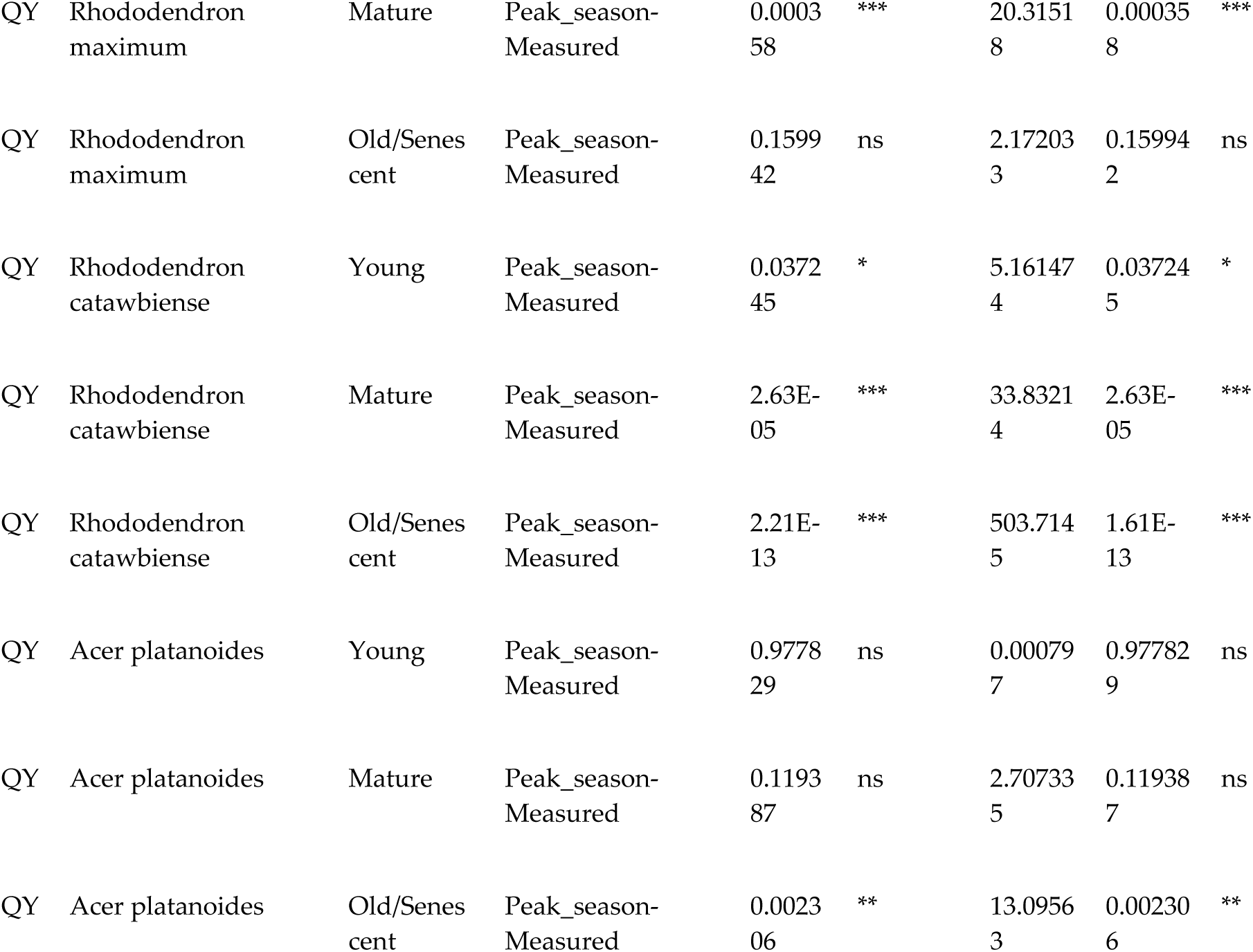
Significant differences across leaf phenophases between measured traits and predicted traits from models trained on one phenophase - Peak season model.

**Table S8.**
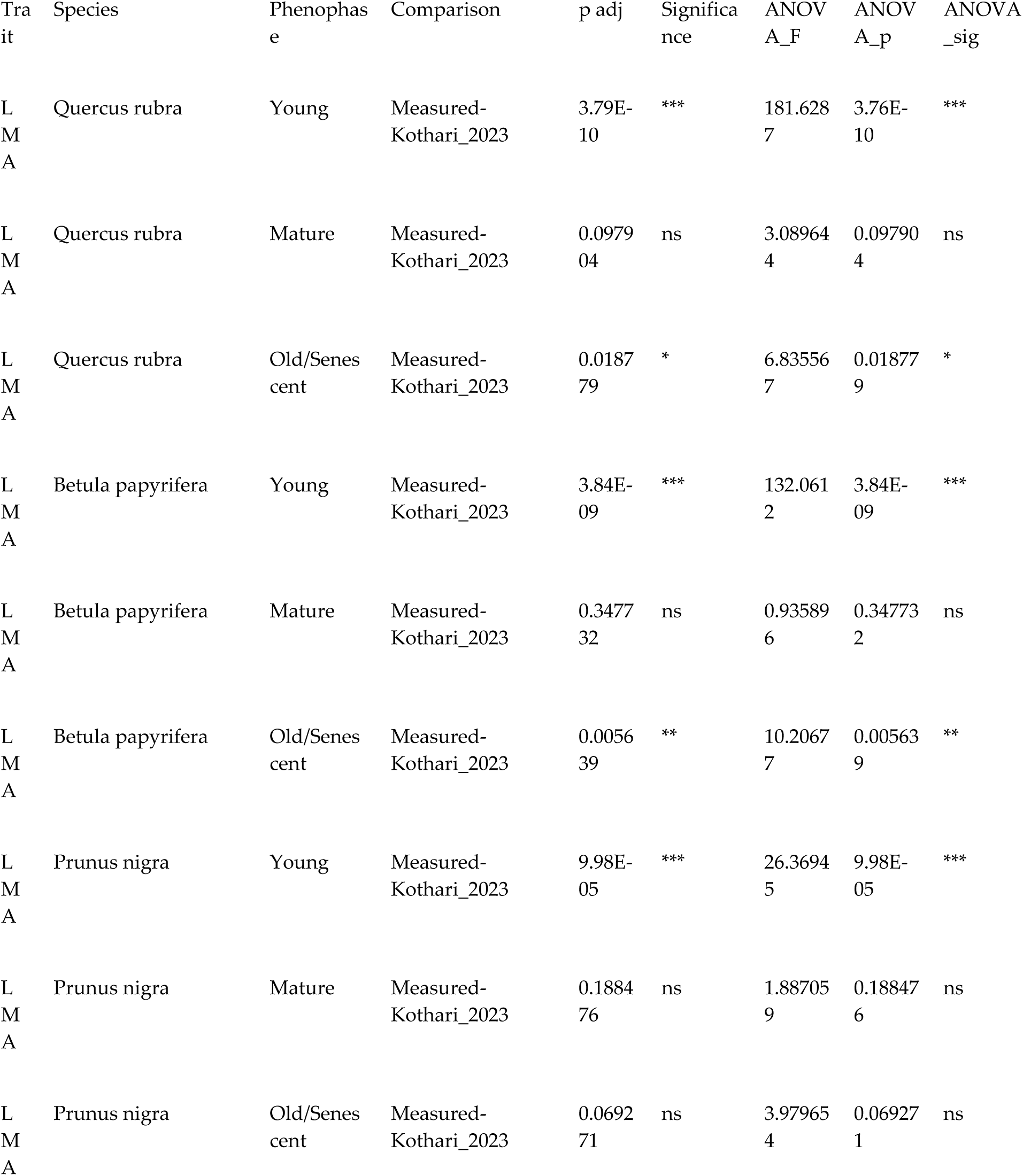
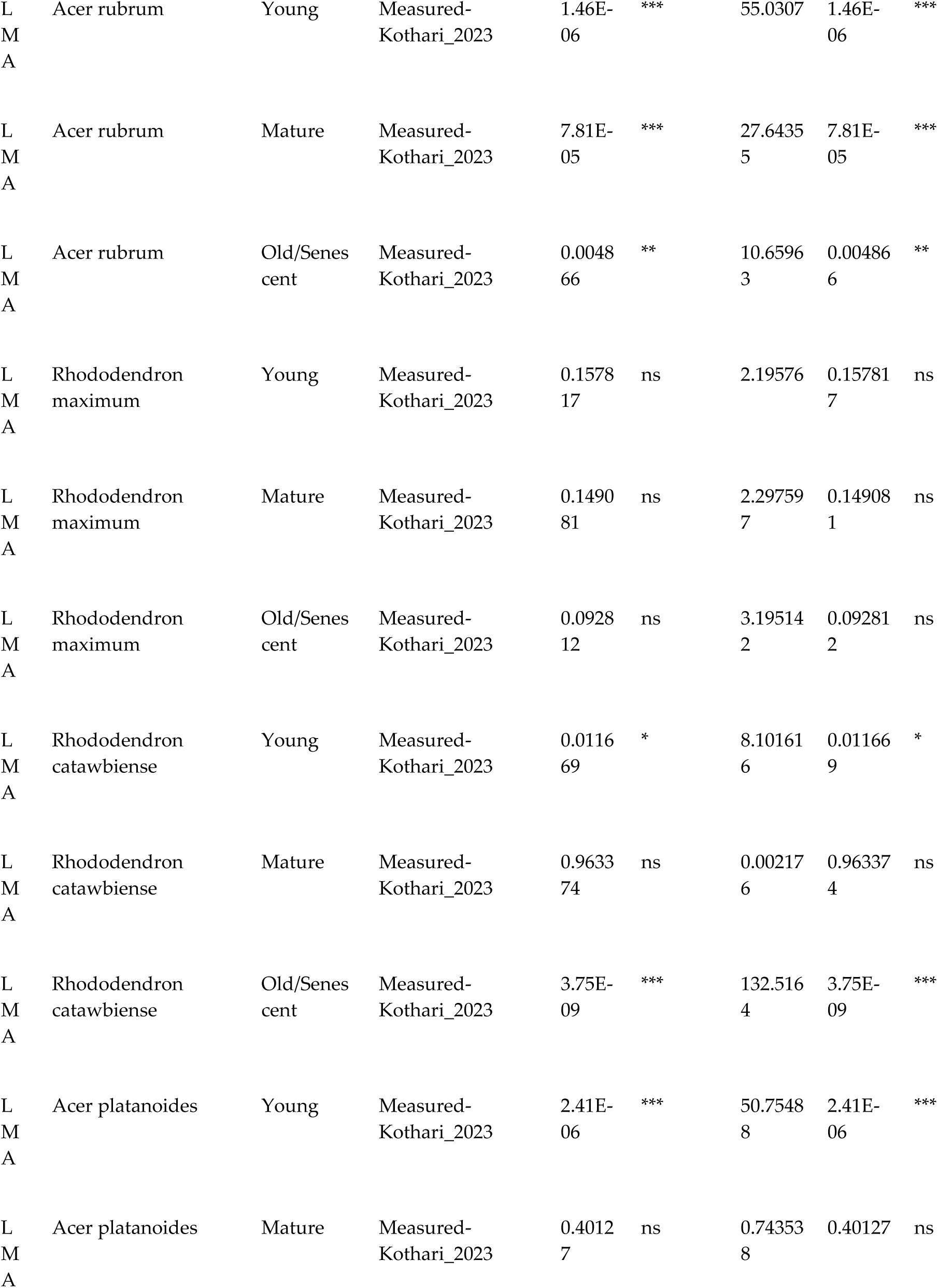
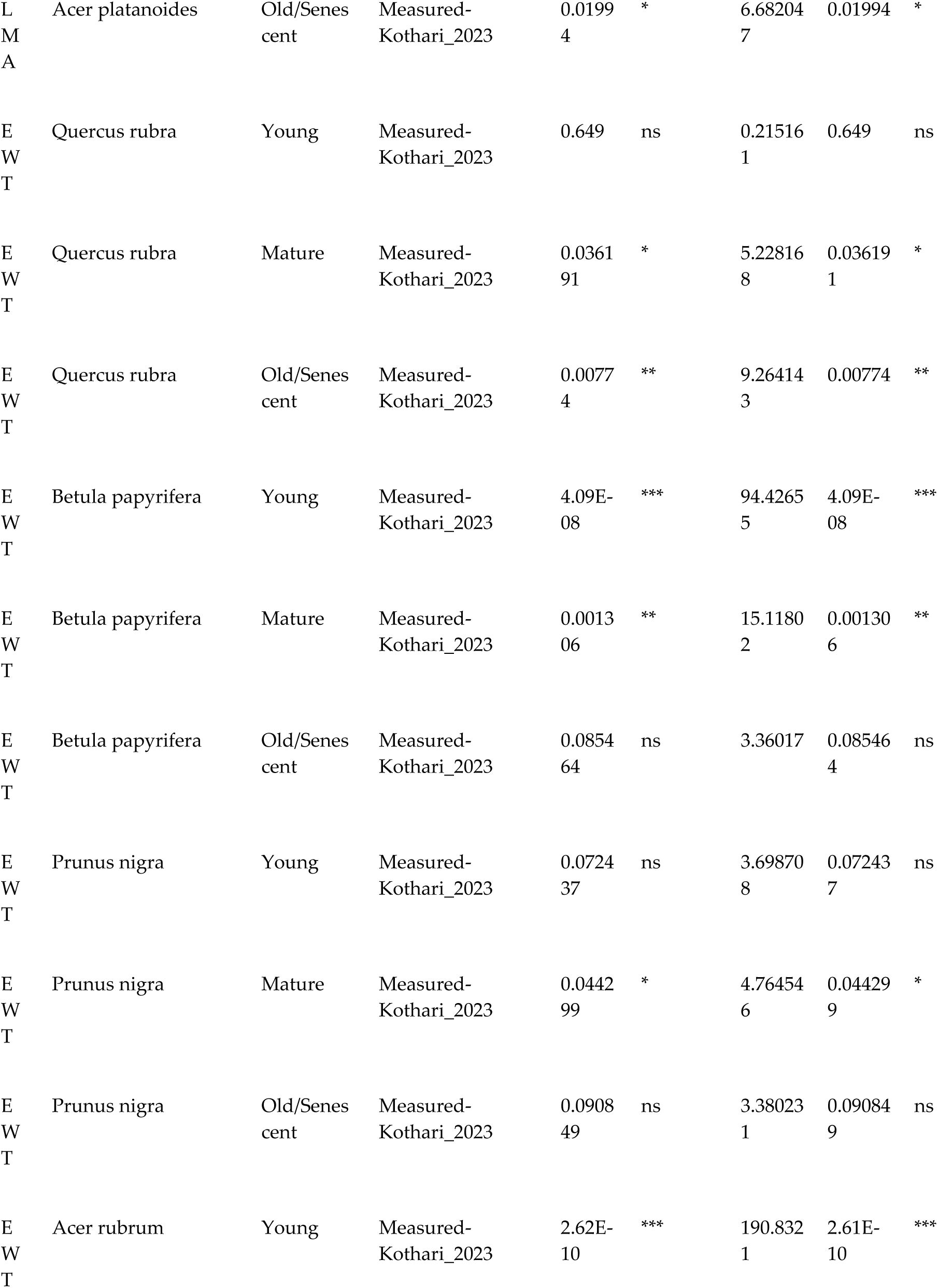
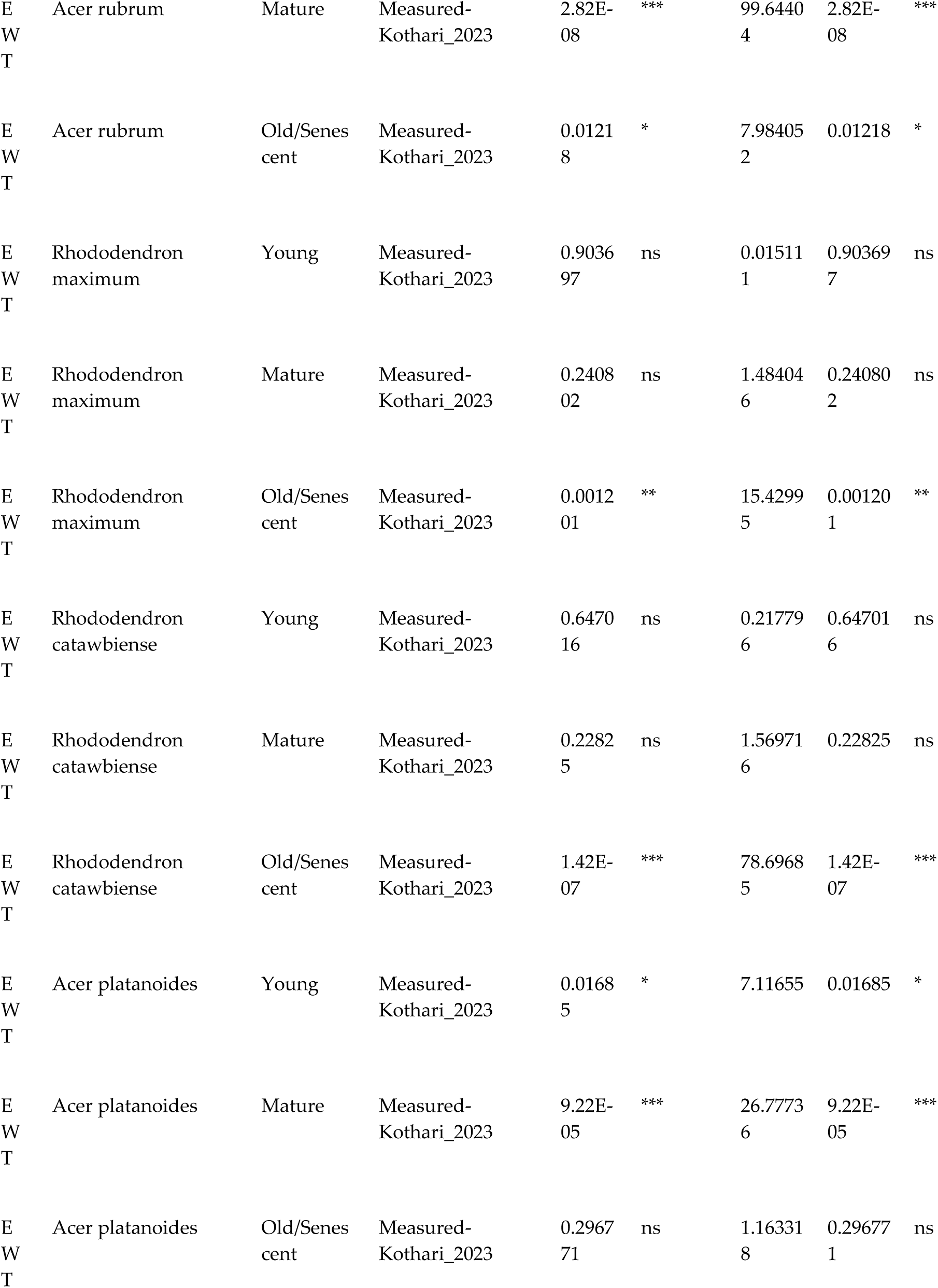
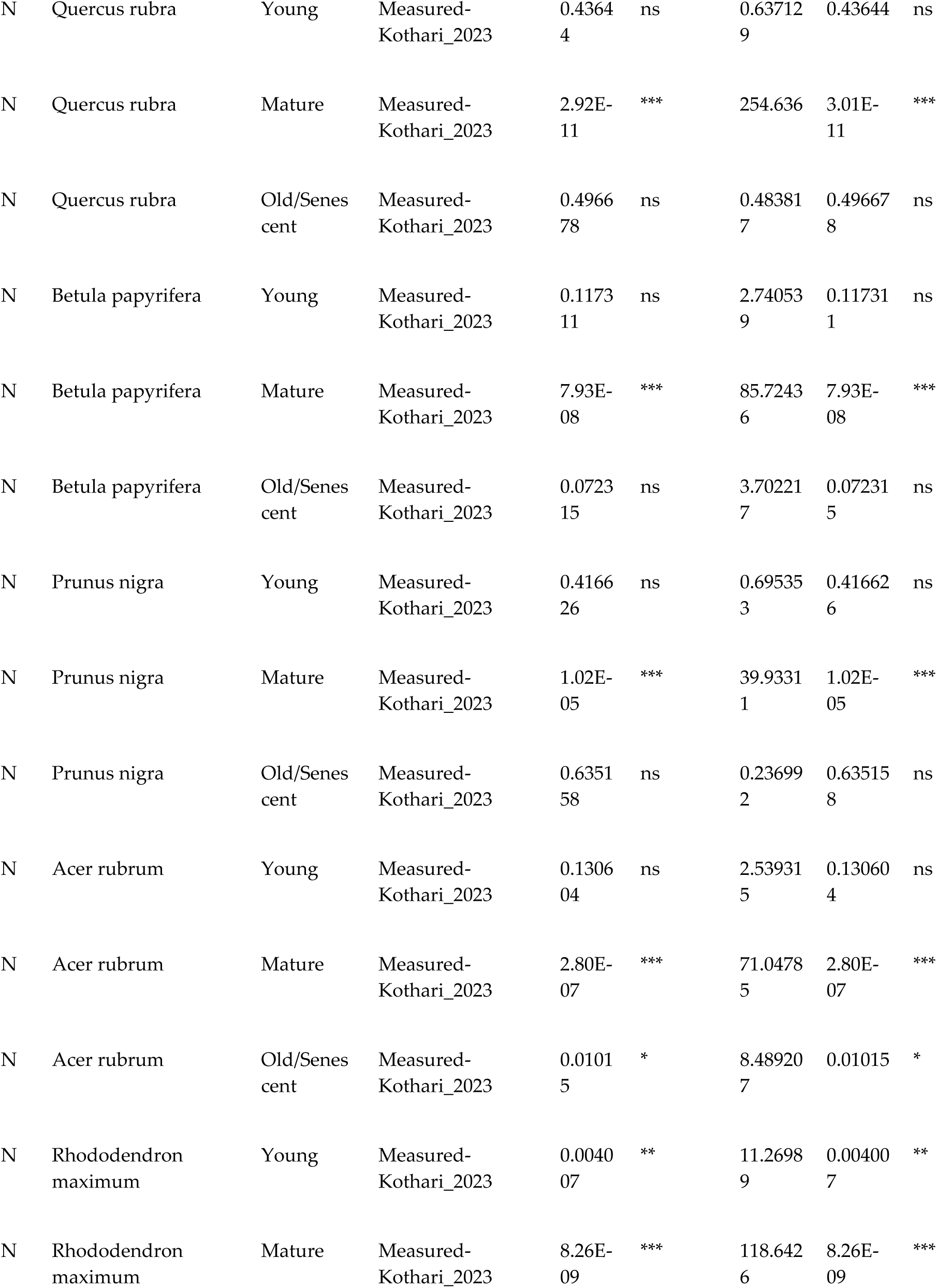
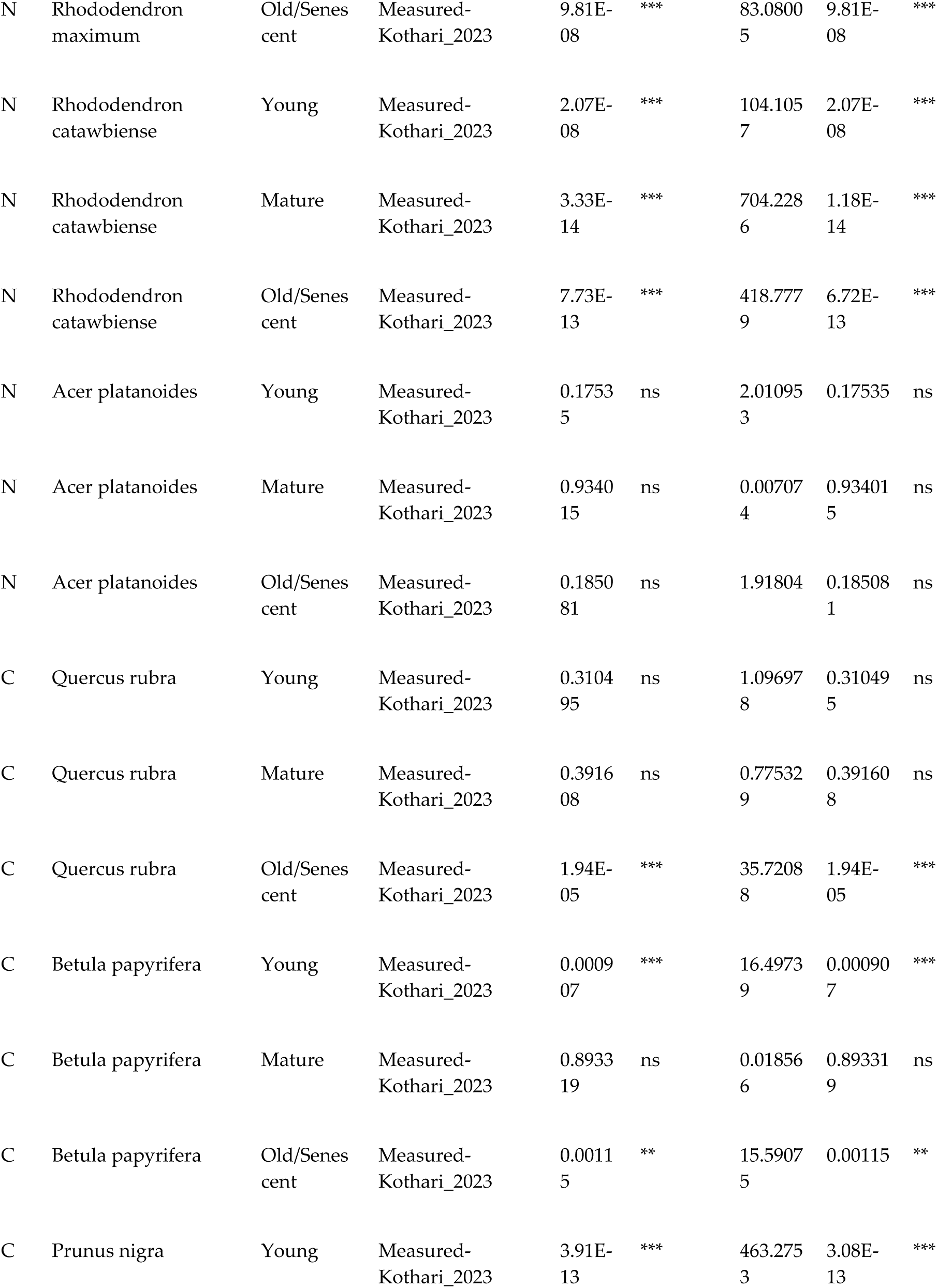
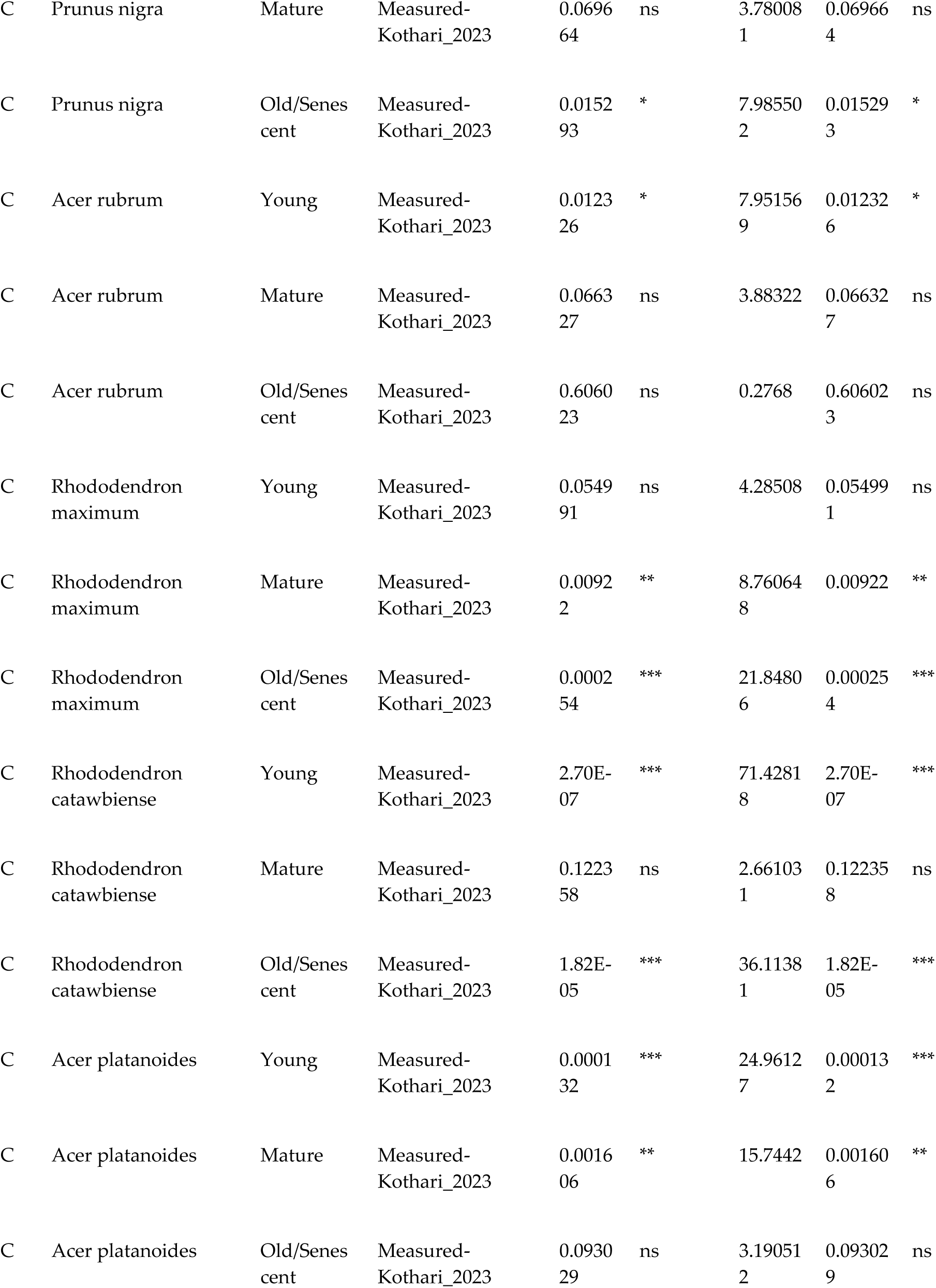
Significant differences across leaf phenophases between measured traits and predicted traits from models trained on one phenophase - Kothari 2023 model.

## REFERENCES

Aguirre-Gutiérrez J, Rifai SW, Deng X, Ter Steege H, Thomson E, Corral-Rivas JJ, Guimaraes AF, Muller S, Klipel J, Fauset S, et al. 2025. Canopy functional trait variation across Earth’s tropical forests. Nature 641: 129–136.

Asner GP, Martin RE, Carranza-Jiménez L, Sinca F, Tupayachi R, Anderson CB, Martinez P. 2014. Functional and biological diversity of foliar spectra in tree canopies throughout the Andes to Amazon region. The New Phytologist 204: 127–139.

de Bello F, Lavorel S, Díaz S, Harrington R, Cornelissen JHC, Bardgett RD, Berg MP, Cipriotti P, Feld CK, Hering D, et al. 2010. Towards an assessment of multiple ecosystem processes and services via functional traits. Biodiversity and Conservation 19: 2873–2893.

Burnett AC, Anderson J, Davidson KJ, Ely KS, Lamour J, Li Q, Morrison BD, Yang D, Rogers A, Serbin SP. 2021a. A best-practice guide to predicting plant traits from leaf-level hyperspectral data using partial least squares regression. Journal of experimental botany 72: 6175–6189.

Burnett AC, Serbin SP, Lamour J, Anderson J, Davidson KJ, Yang D, Rogers A. 2021b. Seasonal trends in photosynthesis and leaf traits in scarlet oak. Tree Physiology 41: 1413–1424.

Caparros-Santiago JA, Rodriguez-Galiano V, Dash J. 2021. Land surface phenology as indicator of global terrestrial ecosystem dynamics: A systematic review. ISPRS Journal of Photogrammetry and Remote Sensing: Official Publication of the International Society for Photogrammetry and Remote Sensing (ISPRS) 171: 330–347.

Cavender-Bares J, Meireles JE, Pinto-Ledezma J, Reich PB, Schuman MC, Townsend PA, Trowbridge A. 2025. Spectral biology across scales in changing environments. Ecology 106: e70078.

Cavender-Bares J, White DM, Ahlstrand NI, Austin MW, Bastianelli D, Bazan S, Boughalmi K, Cardinal-McTeague W, Chacón-Madrigal E, Couvreur TLP, et al. 2026. Next-generation specimen digitization: capturing reflectance spectra from the world’s herbaria for modeling plant biology across time, space, and taxa. The New Phytologist 251: 645–665.

Chadwick K, Asner G. 2016. Organismic-scale remote sensing of canopy foliar traits in lowland tropical forests. Remote Sensing 8: 87.

Chavana-Bryant C, Malhi Y, Anastasiou A, Enquist BJ, Cosio EG, Keenan TF, Gerard FF. 2019. Leaf age effects on the spectral predictability of leaf traits in Amazonian canopy trees. The Science of the total environment 666: 1301–1315.

Chavana-Bryant C, Malhi Y, Wu J, Asner GP, Anastasiou A, Enquist BJ, Cosio Caravasi EG, Doughty CE, Saleska SR, Martin RE, et al. 2017. Leaf aging of Amazonian canopy trees as revealed by spectral and physiochemical measurements. The New Phytologist 214: 1049–1063.

Chlus A, Townsend PA. 2022. Characterizing seasonal variation in foliar biochemistry with airborne imaging spectroscopy. Remote sensing of environment 275: 113023.

Cleland EE, Wolkovich EM. 2024. Effects of phenology on plant community assembly and structure. Annual review of ecology, evolution, and systematics 55: 471–492.

Cope OL, Burkle LA, Croy JR, Mooney KA, Yang LH, Wetzel WC. 2022. The role of timing in intraspecific trait ecology. Trends in ecology & evolution 37: 997–1005.

Cornwell WK, Westoby M, Falster DS, FitzJohn RG, O’Meara BC, Pennell MW, McGlinn DJ, Eastman JM, Moles AT, Reich PB, et al. 2014. Functional distinctiveness of major plant lineages. The Journal of Ecology 102: 345–356.

Couture JJ, Singh A, Rubert-Nason KF, Serbin SP, Lindroth RL, Townsend PA. 2016. Spectroscopic determination of ecologically relevant plant secondary metabolites. Methods in Ecology and Evolution 7: 1402–1412.

Curran PJ. 1989. Remote sensing of foliar chemistry. Remote sensing of environment 30: 271–278.

Damián X, Fornoni J, Domínguez CA, Boege K. 2018. Ontogenetic changes in the phenotypic integration and modularity of leaf functional traits. Functional Ecology 32: 234–246.

Díaz S, Kattge J, Cornelissen JHC, Wright IJ, Lavorel S, Dray S, Reu B, Kleyer M, Wirth C, Colin Prentice I, et al. 2016. The global spectrum of plant form and function. Nature 529: 167–171.

Ely KS, Burnett AC, Lieberman-Cribbin W, Serbin SP, Rogers A. 2019. Spectroscopy can predict key leaf traits associated with source–sink balance and carbon–nitrogen status. Journal of experimental botany 70: 1789–1799.

Escudero A, Mediavilla S. 2003. Decline in photosynthetic nitrogen use efficiency with leaf age and nitrogen resorption as determinants of leaf life span: Leaf life span and nitrogen use efficiency. The Journal of ecology 91: 880–889.

Fajardo A, Siefert A. 2016. Phenological variation of leaf functional traits within species. Oecologia 180: 951–959.

Féret J-B, le Maire G, Jay S, Berveiller D, Bendoula R, Hmimina G, Cheraiet A, Oliveira JC, Ponzoni FJ, Solanki T, et al. 2019. Estimating leaf mass per area and equivalent water thickness based on leaf optical properties: Potential and limitations of physical modeling and machine learning. Remote sensing of environment 231: 110959.

Gomulkiewicz R, Kirkpatrick M. 1992. Quantitative genetics and the evolution of reaction norms. Evolution; International Journal of Organic Evolution 46: 390–411.

Gong Z, Ge W, Guo J, Liu J. 2024. Satellite remote sensing of vegetation phenology: Progress, challenges, and opportunities. ISPRS Journal of Photogrammetry and Remote Sensing: Official Publication of the International Society for Photogrammetry and Remote Sensing (ISPRS) 217: 149–164.

Hörtensteiner S. 2006. Chlorophyll degradation during senescence. Annual Review of Plant Biology 57: 55–77.

Jiang JY, Wang CW, Chen CI, Chen CW, Wong SL, Chen SP, Huang MY, Weng JH. 2024. Photosystem II efficiency in response to diurnal and seasonal variations in photon flux density and air temperature for green, yellow-green, and purple-leaved cultivars of sweet potato [Ipomoea batatas (L.) Lam]. Photosynthetica 62: 116–125.

Ji F, Li F, Hao D, Shiklomanov AN, Yang X, Townsend PA, Dashti H, Nakaji T, Kovach KR, Liu H, et al. 2024. Unveiling the transferability of PLSR models for leaf trait estimation: lessons from a comprehensive analysis with a novel global dataset. The new phytologist 243: 111–131.

Ji F, Zheng T, Shiklomanov AN, Yang R, Townsend PA, Li F, Hao D, Dashti H, Kovach KR, You H, et al. 2026. Tracking seasonal variability in plant traits from spaceborne PRISMA and NEON AOP across forest types and ecoregions. Remote Sensing of Environment 333: 115149.

Kattge J, Bönisch G, Díaz S, Lavorel S, Prentice IC, Leadley P, Tautenhahn S, Werner GDA, Aakala T, Abedi M, et al. 2020. TRY plant trait database - enhanced coverage and open access. Global change biology 26: 119–188.

Kokaly RF, Asner GP, Ollinger SV, Martin ME, Wessman CA. 2009. Characterizing canopy biochemistry from imaging spectroscopy and its application to ecosystem studies. Remote Sensing of Environment 113: S78–S91.

Kothari S, Beauchamp-Rioux R, Blanchard F, Crofts AL, Girard A, Guilbeault-Mayers X, Hacker PW, Pardo J, Schweiger AK, Demers-Thibeault S, et al. 2023. Predicting leaf traits across functional groups using reflectance spectroscopy. The New phytologist 238: 549–566.

Kothari S, Schweiger AK. 2022. Plant spectra as integrative measures of plant phenotypes. The Journal of Ecology 110: 2536–2554.

Kraft NJB, Godoy O, Levine JM. 2015. Plant functional traits and the multidimensional nature of species coexistence. Proceedings of the National Academy of Sciences 112: 797–802.

Kuhn M. 2008. Building Predictive Models inRUsing thecaretPackage. Journal of Statistical Software 28: 1–26.

Laughlin DC, Lusk CH, Bellingham PJ, Burslem DFRP, Simpson AH, Kramer-Walter KR. 2017. Intraspecific trait variation can weaken interspecific trait correlations when assessing the whole-plant economic spectrum. Ecology and Evolution 7: 8936–8949.

Liland KH, Mevik B-H, Wehrens R, Hiemstra P. 2026. *Partial least squares and principal component regression*. CRAN.

Luo Z, Guan H, Zhang X, Liu N. 2017. Photosynthetic capacity of senescent leaves for a subtropical broadleaf deciduous tree species Liquidambar formosana Hance. Scientific reports 7: 6323.

Mason CM, LaScaleia MC, De La Pascua DR, Monroe JG, Goolsby EW. 2020. Learning from dynamic traits: Seasonal shifts yield insights into ecophysiological trade-offs across scales from macroevolutionary to intraindividual. International Journal of Plant Sciences 181: 88–102.

Mayfield MM, Levine JM. 2010. Opposing effects of competitive exclusion on the phylogenetic structure of communities: Phylogeny and coexistence. Ecology Letters 13: 1085–1093.

McGill BJ, Enquist BJ, Weiher E, Westoby M. 2006. Rebuilding community ecology from functional traits. Trends in ecology & evolution 21: 178–185.

McHugh ML. 2011. Multiple comparison analysis testing in ANOVA. Biochemia medica 21: 203–209.

McKown AD, Guy RD, Azam MS, Drewes EC, Quamme LK. 2013. Seasonality and phenology alter functional leaf traits. Oecologia 172: 653–665.

Meireles JE, Cavender-Bares J, Townsend PA, Ustin S, Gamon JA, Schweiger AK, Schaepman ME, Asner GP, Martin RE, Singh A, et al. 2020. Leaf reflectance spectra capture the evolutionary history of seed plants. The New phytologist 228: 485–493.

Meireles JE, Schweiger AK, Cavender-Bares JM. 2017. spectrolab: Class and Methods for Hyperspectral Data.

Mevik B-H, Wehrens R. 2007. TheplsPackage: Principal component and partial least squares regression inR. Journal of Statistical Software 18: 1–23.

Morais MC, Cabral JA, Gonçalves B. 2022. Seasonal variation in selected biochemical traits in the leaves of co-occurring invasive and native plant species under Mediterranean conditions. Plants 11: 1171.

Muller O, Hirose T, Werger MJA, Hikosaka K. 2011. Optimal use of leaf nitrogen explains seasonal changes in leaf nitrogen content of an understorey evergreen shrub. Annals of Botany 108: 529–536.

Nichodemus CO, Meireles JE. 2026. Data for ‘Reflectance spectra capture temporal variation in functional traits and leaf phenology’.

Niittynen P, Merkurjeva M, Kemppinen J. 2026. Seasonal variation of leaf functional traits in sub-Arctic plants. *Oikos (Copenhagen*, Denmark): e11849.

Nunes MH, Davey MP, Coomes DA. 2017. On the challenges of using field spectroscopy to measure the impact of soil type on leaf traits. Biogeosciences 14: 3371–3385.

Ooms J. 2026. Advanced graphics and image-processing in R. ropensci.

Pahadi P, Wason J, Annis S, McGill B, Zhang Y-J. 2024. A new dimension of leaf economic spectrum: temporal instability of relationships among genotypes. The new phytologist.

Palm E, Guidi Nissim W, Gagnon-Fee D, Labrecque M. 2022. Photosynthetic patterns during autumn in three different Salix cultivars grown on a brownfield site. Photosynthesis research 154: 155–167.

Pigliucci M. 2003. Phenotypic integration: studying the ecology and evolution of complex phenotypes. Ecology letters 6: 265–272.

Poorter H, Niinemets Ü, Poorter L, Wright IJ, Villar R. 2009. Causes and consequences of variation in leaf mass per area (LMA): a meta-analysis. The New phytologist 182: 565–588.

Puglielli G, Varone L. 2018. Inherent variation of functional traits in winter and summer leaves of Mediterranean seasonal dimorphic species: evidence of a ‘within leaf cohort’ spectrum. AoB Plants 10: ly027.

Rao Q, Chen J, Chou Q, Ren W, Cao T, Zhang M, Xiao H, Liu Z, Chen J, Su H, et al. 2023. Linking trait network parameters with plant growth across light gradients and seasons. Functional Ecology 37: 1732–1746.

Reich PB. 2014. The world-wide ‘fast–slow’ plant economics spectrum: a traits manifesto. The Journal of ecology 102: 275–301.

Rudolf VHW. 2019. The role of seasonal timing and phenological shifts for species coexistence. Ecology Letters 22: 1324–1338.

Serbin SP, Singh A, McNeil BE, Kingdon CC, Townsend PA. 2014. Spectroscopic determination of leaf morphological and biochemical traits for northern temperate and boreal tree species. Ecological applications: a publication of the Ecological Society of America 24: 1651–1669.

Siefert A, Violle C, Chalmandrier L, Albert CH, Taudiere A, Fajardo A, Aarssen LW, Baraloto C, Carlucci MB, Cianciaruso MV, et al. 2015. A global meta-analysis of the relative extent of intraspecific trait variation in plant communities. Ecology Letters 18: 1406–1419.

Šímová I, Violle C, Svenning J-C, Kattge J, Engemann K, Sandel B, Peet RK, Wiser SK, Blonder B, McGill BJ, et al. 2018. Spatial patterns and climate relationships of major plant traits in the New World differ between woody and herbaceous species. Journal of biogeography 45: 895–916.

Skillman JB. 2008. Quantum yield variation across the three pathways of photosynthesis: not yet out of the dark. Journal of experimental botany 59: 1647–1661.

Slaton MR, Hunt ER, Smith WK. 2001. Estimating near-infrared leaf reflectance from leaf structural characteristics. American Journal of Botany 88: 278–284.

Stasinski L, White DM, Nelson PR, Ree RH, Meireles JE. 2021. Reading light: leaf spectra capture fine-scale diversity of closely related, hybridizing arctic shrubs. The New phytologist 232: 2283–2294.

Sultan SE. 2000. Phenotypic plasticity for plant development, function and life history. Trends in Plant Science 5: 537–542.

Tanaka A, Ito H. 2025. Chlorophyll degradation and its physiological function. Plant & Cell Physiology 66: 139–152.

Turner W, Spector S, Gardiner N, Fladeland M, Sterling E, Steininger M. 2003. Remote sensing for biodiversity science and conservation. Trends in Ecology & Evolution 18: 306–314.

Ustin SL, Gitelson AA, Jacquemoud S, Schaepman M, Asner GP, Gamon JA, Zarco-Tejada P. 2009. Retrieval of foliar information about plant pigment systems from high resolution spectroscopy. Remote Sensing of Environment 113: S67–S77.

Ustin SL, Jacquemoud S. 2020. How the optical properties of leaves modify the absorption and scattering of energy and enhance leaf functionality. In: Remote Sensing of Plant Biodiversity. Cham: Springer International Publishing, 349–384.

Violle C, Enquist BJ, McGill BJ, Jiang L, Albert CH, Hulshof C, Jung V, Messier J. 2012. The return of the variance: intraspecific variability in community ecology. Trends in Ecology & Evolution 27: 244–252.

Violle C, Navas M-L, Vile D, Kazakou E, Fortunel C, Hummel I, Garnier E. 2007. Let the concept of trait be functional! Oikos 116: 882–892.

Vogelmann TC. 1993. Plant tissue optics. Annual Review of Plant Physiology and Plant Molecular Biology 44: 231–251.

Wang Z, Féret J-B, Liu N, Sun Z, Yang L, Geng S, Zhang H, Chlus A, Kruger EL, Townsend PA. 2023. Generality of leaf spectroscopic models for predicting key foliar functional traits across continents: A comparison between physically- and empirically-based approaches. Remote sensing of environment 293: 113614.

Wickham H. 2016. Ggplot2: Elegant graphics for data analysis. Cham, Switzerland: Springer International Publishing.

Wold S, Sjöström M, Eriksson L. 2001. PLS-regression: a basic tool of chemometrics. Chemometrics and Intelligent Laboratory Systems 58: 109–130.

Wright IJ, Reich PB, Cornelissen JHC, Falster DS, Garnier E, Hikosaka K, Lamont BB, Lee W, Oleksyn J, Osada N, et al. 2005. Assessing the generality of global leaf trait relationships. The new phytologist 166: 485–496.

Wright IJ, Reich PB, Westoby M, Ackerly DD, Baruch Z, Bongers F, Cavender-Bares J, Chapin T, Cornelissen JHC, Diemer M, et al. 2004. The worldwide leaf economics spectrum. Nature 428: 821–827.

Yang X, Tang J, Mustard JF, Wu J, Zhao K, Serbin S, Lee J-E. 2016. Seasonal variability of multiple leaf traits captured by leaf spectroscopy at two temperate deciduous forests. Remote sensing of environment 179: 1–12.

Yu L, Luo X, Zhao R, Satriawan TW, Tian J. 2025. The spatiotemporal variations in ecosystem photosynthetic quantum yield and their drivers. Agricultural and Forest Meteorology 365: Not Available.

Zhao K, Valle D, Popescu S, Zhang X, Mallick B. 2013. Hyperspectral remote sensing of plant biochemistry using Bayesian model averaging with variable and band selection. Remote sensing of environment 132: 102–119.

Zhao Y-W, Wang C-K, Huang X-Y, Hu D-G. 2021. Anthocyanin stability and degradation in plants. Plant Signaling & Behavior 16: 1987767.

